# Systematic Analysis of Network-driven Adaptive Resistance to CDK4/6 and Estrogen Receptor Inhibition using Meta-Dynamic Network Modelling

**DOI:** 10.1101/2023.01.24.525460

**Authors:** Anthony Hart, Sung-Young Shin, Lan K. Nguyen

**Author notes:** To whom correspondence should be addressed (L.K.N).

## Abstract

Drug resistance inevitably emerges during the treatment of cancer by targeted therapy. Adaptive resistance is a major form of drug resistance, wherein the rewiring of protein signalling networks in response to drug perturbation allows drug-targeted protein activity to recover. This can occur in the continuous presence of the drug and enables cells to survive/grow. Simultaneously, molecular heterogeneity enables the selection of drug-resistant cancer clones that can survive an initial drug insult, proliferate, and eventually cause disease relapse. Despite their importance, the link between heterogeneity and adaptive resistance, specifically how heterogeneity influences protein signalling dynamics to drive adaptive resistance, remains poorly understood. Here, we have explored the relationship between heterogeneity, protein signalling dynamics and adaptive resistance through the development of a novel modelling technique coined Meta Dynamic Network (MDN) modelling. We use MDN modelling to characterise how heterogeneity influences the drug-response signalling dynamics of the proteins that regulate early cell cycle progression and demonstrate that heterogeneity can robustly facilitate adaptive resistance associated dynamics for key cell cycle regulators. We determined the influence of heterogeneity at the level of both reaction coefficients and protein abundance and show that reaction coefficients are a much stronger driver of adaptive resistance. Owing to the mechanistic nature of the underpinning ODE framework, we then identified a full spectrum of subnetworks capable of driving adaptive resistance dynamics in the key early cell cycle regulators. Finally, we show that single-cell dynamic data supports the validity of our MDN modelling technique and a comparison between our predicted resistance mechanisms and known CDK4/6 and Estrogen Receptor inhibitor resistance mechanisms suggests MDN can be deployed to robustly predict network-level resistance mechanisms for novel drugs and additional protein signalling networks.

## INTRODUCTION

Drug resistance is a widespread phenomenon across all cancer types and is a major obstacle to the development of curative therapeutic strategies (1–3). Cellular heterogeneity is a known driver of drug resistance, wherein the treatment of a heterogenous population of cancer cells with cytostatic or cytotoxic drugs creates a selective pressure that results in the survival and expansion of any cells that are capable of overcoming the effects of said drugs (4–6). While this general phenomenon is well established (7–9), it is usually less clear exactly how and why some cells are able to overcome a drug treatment and others are not. Heterogeneity is a catch-all term used to describe any and all differences between cells, however, it could be argued that the majority of the differences between cells ultimately converge on differences in how their constituent proteins behave, i.e. their protein dynamics. To deepen our understanding of how tumour heterogeneity drives drug resistance, we must explore the relationship between cellular heterogeneity, protein dynamics and drug resistance.

The efficacy of a targeted therapy largely comes down to how well and how long the targeted protein is suppressed, and how frequently and reliably this suppression results in either cytostasis or apoptosis. A cell can therefore be considered resistant if the target protein is insufficiently suppressed or is initially suppressed but later recovers, or if the suppression of the protein is insufficient to stimulate cytostasis or apoptosis (10, 11). Proteins are embedded in complex networks and protein dynamics are dictated by the properties of the networks in which they reside. While resistance is often due to direct effects, such as reduced drug-target binding affinity (12, 13) or excessive drug efflux from tumour cells (14, 15), it can also occur due to changes in the state of the biochemical networks of a cell that counteract the effects of the drug; a phenomenon referred to as adaptive resistance (16–18). PI3K, EGFR and CDK4/6 are just few prime examples of proteins that have been targeted in the treatment of cancer that display acute and robust adaptive resistance (19–21). These studies also support the notion that it is rarely the behaviour of a single ‘gatekeeper’ protein that drives adaptive resistance, but an entire protein network that acts and is acted upon by the target protein (22, 23). Frequently however, we possess a limited knowledge and appreciation of the network-level mechanistic relationships that underpin adaptative resistance.

Understanding the relationship between cellular heterogeneity and adaptive drug resistance requires an ability to explore and characterise the heterogeneity that exists both within a tumour and between patients. This can be achieved *in silico* by investigating the behaviour of a network, i.e. the dynamics of its constituent proteins, over a broad range of network conditions, allowing us to observe the full spectrum of possible dynamics that can be displayed by a given network topology. A particularly useful technique for investigating protein signalling networks is Ordinary Differential Equation (ODE) modelling (24). An ODE model is a mathematical representation of how the protein species within a network interact and evolve over time. ODE models have been extensively used to simulate and predict network-level responses to perturbations, such as growth-factor stimulation or drug treatment (25–28). In an ODE model, the interactions between the network’s constituent proteins are converted into mathematical formulations using well-established biochemical rate laws (29–33). The end result is a set of ODEs – the model – that can be numerically solved using specialised ODE solvers, allowing one to simulate, and thus predict, how the concentrations of the protein species will change over time or in response to perturbations (29–33).

An ODE model is comprised of state variables, initial conditions, model parameters, and defined inputs and outputs (29–31). Typically, the parameters, or reaction coefficients, represent the strength of protein interactions and state variables correspond to the concentrations of the protein species. Initial conditions (ICs) are the values of the state variables at the starting point of a simulation and represent the total abundance levels of the protein species within the model. Model input(s) are usually a perturbation to the modelled network, such as a growth factor stimulation or a drug inhibition. We can model network heterogeneity by producing a suite of ODE models, each with their own set of parameter values and/or initial conditions. In this manner, each model ‘instance’ represents a unique cellular context and produces a unique set of dynamic behaviours for its constituent proteins. By varying the parameters and initial conditions over wide ranges, we can delineate an extreme upper limit to the heterogeneity in dynamics facilitated by a network topology.

The process of creating models almost always involves a degree of abstraction, either explicitly, due to conscious decisions by the creator, or implicitly, due to imperfect knowledge of the system being modelled. This abstraction forces a model’s parameters to capture information beyond that which is being explicitly modelled and can potentially render the concept of a ‘true’ or ‘biologically accurate’ parameter value less meaningful. There is an array of factors that can influence reaction coefficients: post-translational modifications, catalysts/scaffolds, pH, etc., and they can all individually influence reaction coefficients over several orders of magnitude. In the context of cancer, this variability in reaction coefficients can become extreme, with mutations altering the properties of proteins and dramatically changing how strongly they interact with other proteins. By varying the parameters and initial conditions over wide ranges, we also potentially capture protein dynamics driven by factors we haven’t explicitly modelled.

Drug resistance is a problem almost as old as drug treatments; and many computational modelling frameworks have been developed to investigate the emergence of drug resistance during cancer treatment (34–36). A comprehensive review of the modelling frameworks developed to explore the relationship between cellular heterogeneity and drug resistance, in particular, was written by Chisholm et al. in 2016 (35). Most of the models covered within focused on population dynamics and were used to identify key processes that govern how resistance emerges within a heterogenous cell population. Notably, none of the models in this review attempted to link heterogenous intracellular processes with adaptive resistance mechanisms, or resistance in general. Perhaps most similar to the ensemble approach developed within this project, He et al. developed an ODE-based modelling technique termed RACIPE (random circuit perturbation) (37). In contrast to our work, RACIPE was a quasi-Monte Carlo method developed to alleviate the necessity of identifying precise kinetic parameter values on which model predictions are supposedly dependent. While their technique does involve the exploration of a range of parameter values, it does so to make predictions about broadly robust outcomes in gene regulatory circuits, not to link parameter variation to heterogenous protein network dynamics. Thus, there is a need for a novel computational framework capable of systematically characterising network-dynamic heterogeneity and its relationship with adaptive resistance.

To address this need, we have developed a new modelling framework, coined Meta-Dynamic Network (MDN) modelling, which enables us to study the effects of network heterogeneity on protein signalling dynamics. In this paper we have applied MDN modelling to an Early Cell Cycle (ECC) network model to systematically explore and illuminate adaptive resistance mechanisms that arise in response to targeted CDK4/6 and Estrogen Receptor (ER) inhibition. The ECC network model developed herein incorporates the two major mitogenic signalling pathways that are largely responsible for cell cycle initiation, and the G1-S phase cell cycle regulatory network.

Using MDN and the ECC network model, we have demonstrated that variation in both the strength of reaction coefficients (parameters) and the total abundance of proteins (initial condition values) can affect the qualitative shape and features of a network’s protein dynamics. Furthermore, we provide evidence that parameter variation induces a much greater variety in the qualitative shape and features of the network’s protein dynamics and is therefore much more likely to give rise to adaptive resistance dynamics. We also show that there is a surprising large number of contexts in which the network topology of the ECC network can facilitate resistance dynamics, and that we can dissect these contexts to systematically characterise their underpinning kinetic drivers. Finally, we analysed existing publicly available biological data to qualitatively validate the existence of a spectrum of signalling dynamics in response to CDK4/6 inhibitors and propose a novel relationship between population signalling dynamics and population level drug sensitivity.

## MATERIALS AND METHODS

### ODE model construction, modelling, and simulations

We converted the ECC network (**Figure 1**) into a system of ODEs using standard biochemical rate laws: catalysed reactions (e.g. phosphorylations) as pseudo-first-order, complex formations as mass action, and transcription, translation and degradation as first-order kinetics. We used these simpler forms to reduce the number of poorly constrained parameters and enable a broader sampling of parameter values in the meta-dynamic analysis. The model comprises 50 state variables, 94 parameters, two mitogenic inputs and two drug inputs. A detailed description of model scope and justification is given in the **Supplementary Information**; and the biochemical reaction lists, ODEs and SBML model files are provided as **Model Files S1 - S3**.

**Figure 1:**
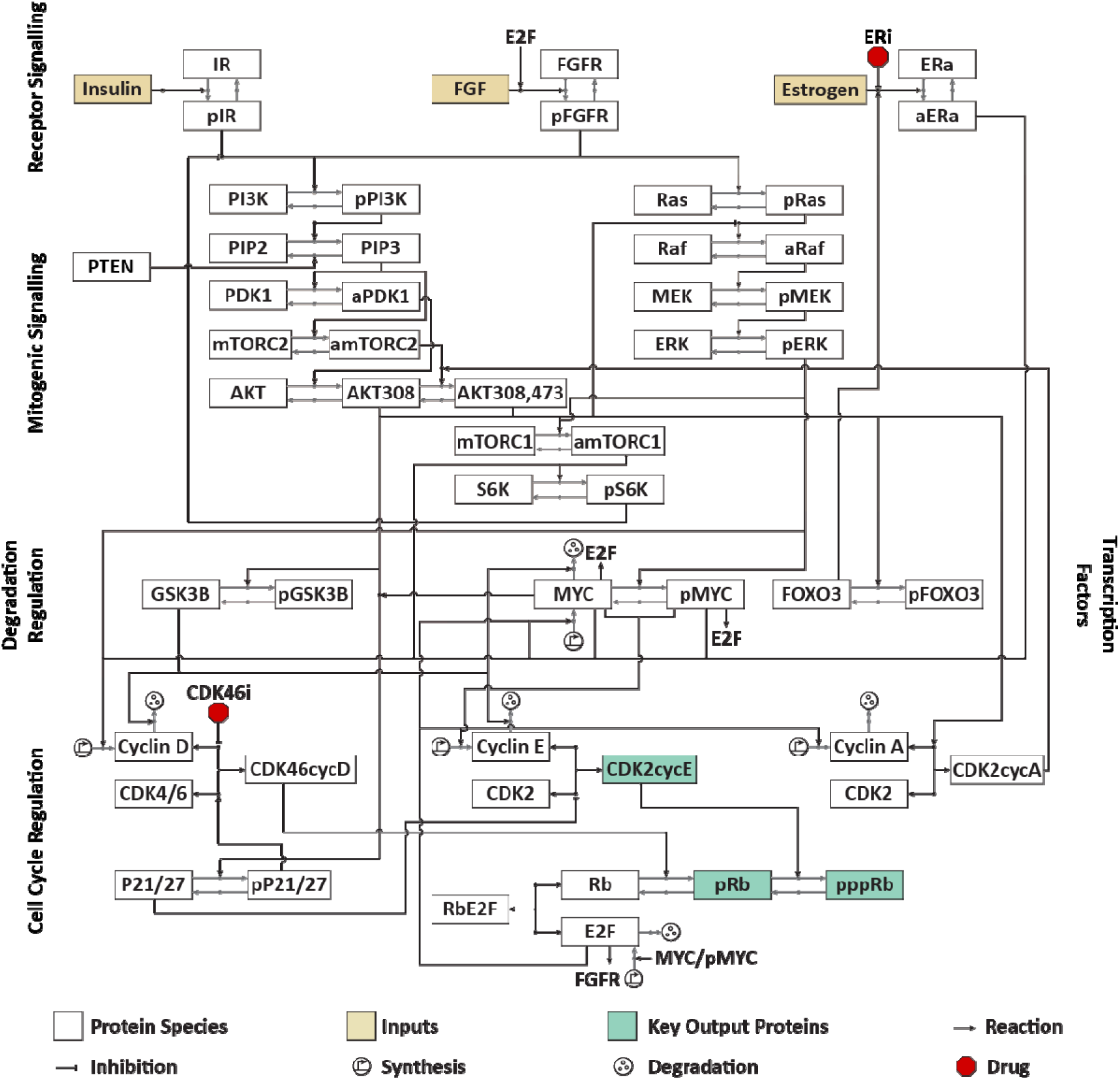
**Network Schematic of the Early Cell Cycle (ECC) Network**. Network schematic showing the detailed biochemical reactions that regulate the initiation of the G1 phase and transition through to the S phase of the cell cycle. The network includes three mitogenic inputs, two mitogenic signalling pathways, PI3K and MAPK, key transcription factors that promote G1 phase transition, key CDK-cyclin complexes and their regulators, a CDK4/6 inhibitor, and an estrogen receptor inhibitor. Canonically, mitogenic stimulation leads to the cascading activation of the mitogenic signalling pathways, which converge on the synthesis of cyclin D. The synthesis of cyclin D then promotes the activation of CDK4/6, which consequently drives the synthesis of cyclin E. Cylin E activates CDK2, leading to the synthesis of cyclin A and the formation of CDK2-cyclin A complexes signals the initiation of S phase. ER and CDK4/6 inhibitors are believed to disrupt this linear chain of events and thus prevent proliferation.

Simulations were implemented in MATLAB (The MathWorks. Inc. 2019a) with IQM toolbox (http://www.intiquan.com/intiquan-tools/) to auto-generate ODEs and compile a MEX file for accelerated simulation speed. ODEs were solved using the SUNDIALS CVODES solver, which employs a variable-order, variable-step Backward Differentiation Formula (BDF) method, designed to solve stiff ODE systems.

Each simulation of our model consisted of three phases: (i) **starvation** phase to ensure a steady state equilibrium was reached before mitogenic stimulation; (ii) **fed** with constant mitogenic stimulation (100 nM of insulin and FGF) until quasi-steady state; and (iii) treatment with CDK4/6i and ER inhibitor. Mitogens and drugs were modelled as step inputs at the start of their respective phases and then held constant for the remaining simulation time, isolating network-intrinsic adaptation rather than responses to input removal. Drug inhibitions were modelled by reducing the relevant parameter values by a fixed fraction.

### Sampling and filtering of model instances

MDN modelling involves random sampling of parameter sets to create unique model instances. We drew random parameter sets and conserved-total initial conditions and continued sampling until we obtained N=100,000 accepted instances for each analysis. To ensure sampled parameter sets reflected plausible biological contexts while retaining rare network states, we applied several filtering steps. An instance was accepted if it met the following: (i) **Quasi–steady state** reached in the starvation and fed phases, defined by very small changes across species within a time cap (1% change in concentration over the final 100 time steps); (ii) **Non-trivial on-target drug effect** during treatment (e.g. ≥5% reduction in the CDK4/6 activity proxy or [CDK4/6·CycD] level for CDK4/6i, and in [active ER] level for ERi). Moreover, we note that multi-timescale signalling models are often stiff (38, 39), and random parameter draws can create extreme early transients that are numerically hard to integrate even with stiff solvers. We therefore excluded these parameter instances, which accounts for only <1% of total sampled sets.

Applying these filters, ∼40–45% of instances did not reach steady state within the allotted time, and ∼50–55% did not meet the minimum drug-response criterion. Approximately 10% satisfied all criteria and were retained for analysis. Importantly, we employed ‘rejection sampling’ and continued drawing until we had N = 100,000 accepted instances that satisfied all the criteria.

### Quantitative definitions of dynamic patterns

A major component of MDN modelling is the analysis of the qualitative shape of individual protein dynamics. This enables us to group together model instances that produce broadly similar behaviours for any proteins being investigated. We can then analyse these groups to identify and elucidate shared network structures that drive the behaviour. To this end, we defined six qualitative protein dynamic categories: increasing, decreasing, biphasic, rebound, no-response and other.

To assess the qualitative shape of each protein dynamic, we developed a series of mathematical restraints that were capable of sorting protein simulations into their appropriate categories.

First, each protein dynamic was normalised to its initial value. A protein’s dynamic was categorised as **increasing (INC)** if (*i*) the final concentration exceeded 20% of its initial concentration, (*ii*) over the entire simulated time period it never dropped below 10% of its initial concentration, and (*iii*) the final concentration was within 10% of the maximum concentration. It was categorised as **decreasing (DEC)** if (*i*) the final concentration was at least 20% lower than the initial concentration, (*ii*) never went above 10% of its initial concentration, and (*iii*) the final concentration was within 10% of the minimum concentration. It was categorised as **biphasic (BIP)** if (*i*) the maximum concentration exceeded the initial concentration by 20% and (*ii*) the final concentration was at least 10% lower than the maximum concentration. It was categorised as **rebound (REB)** if (*i*) the minimum concentration was at least 20% lower than the initial concentration and (*ii*) the final concentration was at least 10% bigger than the minimum concentration. It was categorised as **no-response (NRP)** if the concentration never exceeded more or less than 10% of the initial concentration. Finally, it was categorised as **other (ETC)** if the protein’s dynamic did not fit one of the previously described categories.

### Key model output proteins

There is strong evidence to suggest that the linear cascade of cell cycle regulator progression, wherein the activation of CDK4/6 leads to the activation of CDK2-cyclin E complex, which in turn leads to the activation of CDK2-cyclin A, is overly simplistic (40, 41). If this traditional understanding was correct, the inhibition of CDK4/6 activity would robustly inhibit cell cycle progression, yet this is not the case (42). A number of studies have suggested that the activation or reactivation of proteins downstream of CDK4/6 can enable a cell to overcome the cell cycle inhibitory effects of CDK4/6 targeting drugs and enable cell cycle progression. Proteins that have been shown to demonstrate this phenomenon include phosphorylated Rb and the active CDK2-cyclin E complex (21, 43, 44). As such, we have focused on the dynamics of monophosphorylated Rb (**pRb**), hyperphosphorylated Rb (**pppRb**) and the CDK2-cyclin E complex (**CDK2cycE**) throughout this project, as key mediators of adaptive resistance to CDK4/6 and ER inhibitors.

### Protein knockdown perturbation analyses

To assess the effects of parameters on protein dynamics we performed high-throughput perturbation analyses wherein each parameter was knocked down one at a time (OAT) and the effect on the nominated output assessed. We chose the OAT design intentionally to obtain causal, first-order attribution of control points across a broad parameter ensemble without confounding from simultaneous co-inhibition. This provides an interpretable ranking of primary drivers that is consistent with the paper’s mechanistic focus.

## RESULTS

### An Overview of Meta-Dynamic Network (MDN) Modelling

To explore the link between cellular heterogeneity, protein dynamics and drug resistance, we first developed a novel modelling framework coined Meta-Dynamic Network (MDN) modelling. The principal purpose of the MDN modelling framework is to explore how parametric and initial condition variation influence the qualitative shape of protein dynamics within a given network. By varying model parameters and initial condition (**IC**) values we are able capture many sources of cellular heterogeneity and characterise their influence on protein signalling dynamics. Varying parameter values captures heterogeneity in the interactions between proteins that arises due to phenomena such as cell-specific mutations, post-translational modifications, and epigenetic modifications. Varying initial condition values captures heterogeneity in protein concentration that arises due to phenomena such as stochastic protein expression, copy number variation and tissue-specific expression programmes.

Once a large number of unique model instances have been generated, each with their own unique parameter and/or initial condition values, we can simulate each model’s response to drug treatment and analyse the resulting shape of each protein’s dynamic. From this analysis we can determine if there are contexts where key output proteins, such as downstream targets of the drug perturbation, demonstrate protein dynamics associated with adaptive resistance. Finally, we can group model instances that display similar dynamics together and perform further analyses to systematically identify the mechanistic causes capable of driving adaptive resistance dynamics. An overview of the MDN modelling process can be seen in **Figure 2**.

**Figure 2.**
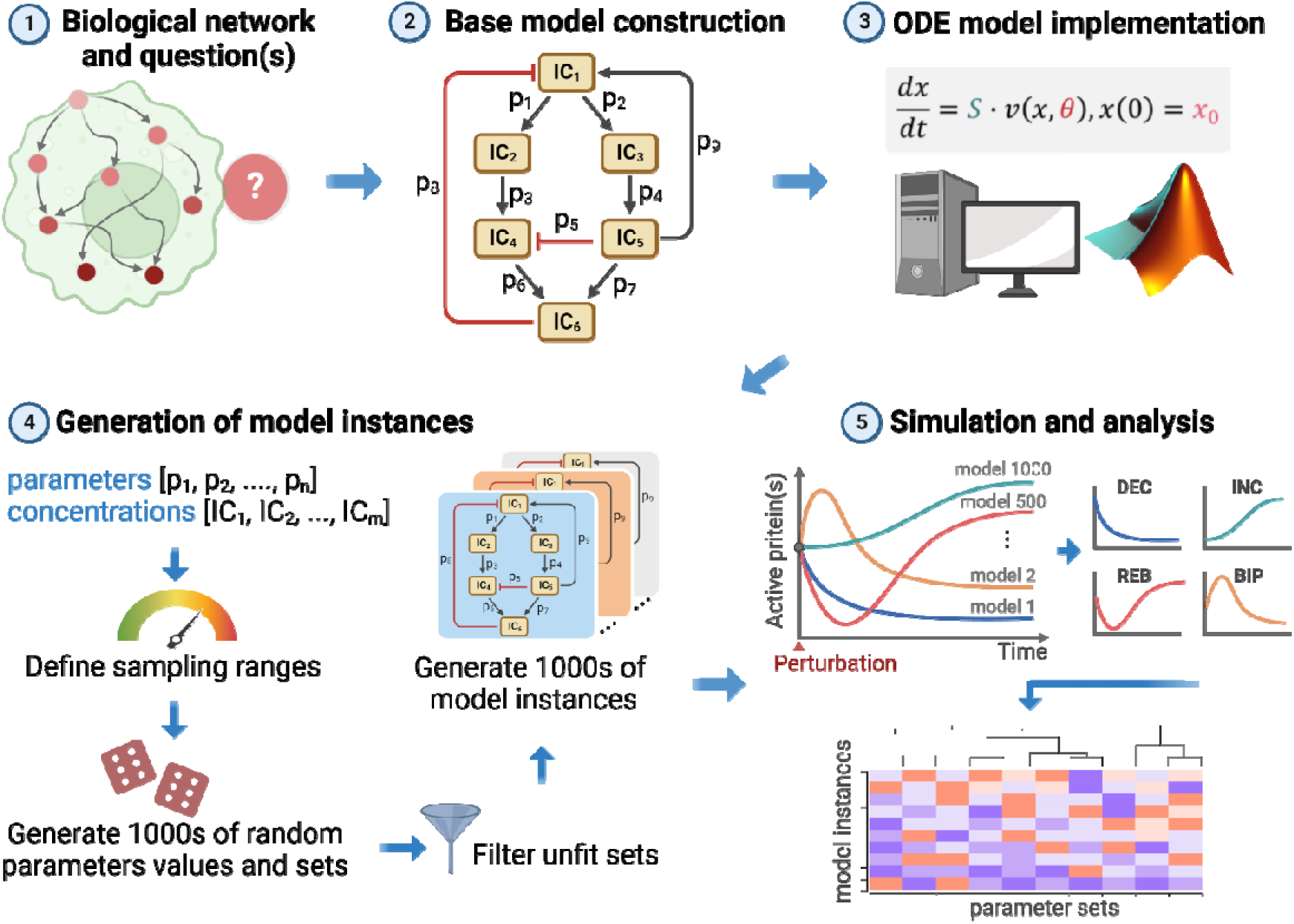
A workflow of the Meta Dynamic Network (MDN) modelling pipeline. As with all modelling, the first step is asking a useful question. In our case this is usually related to understanding complex protein signalling dynamics. The second step is defining the network; this involves selecting nodes and identifying relationships between them. The third step is translating the network schematic to a mathematical representation, which is typically a combination of ordinary differential equations and biochemical rate laws. The fourth step is the generation of thousands of model instances, each with a unique set of properties e.g. parameter and initial condition values. The fifth step is the simulation of each model instance and the bulk analysis of the simulations. In this work, we analyse the distribution of the qualitative dynamics of the network’s protein species.

Protein dynamics that are associated with adaptive resistance are those where the protein is insufficiently suppressed over a sustained time period, i.e. increasing, biphasic and rebound (18–20, 22, 45). Increasing is the most dramatic adaptive-resistance response, where the drug perturbation not only fails to suppress key output proteins, but actually increases their activity/expression. Biphasic is similar to increasing in its initial phase, where the outputs actually increase, however the degree of the second decreasing phase likely influences the degree of resistance conferred by this particular behaviour. Rebound is similar to biphasic wherein the degree of resistance conferred by this dynamic is likely influenced by how strongly the protein recovers. While rebound is probably the most commonly observed adaptive-resistance driving dynamic (18–23), for the purpose of this project we consider all three of these behaviours as adaptive-resistance associated behaviours and consider decreasing as the only sensitivity associated behaviour.

### Construction of a mechanistic model of the Early Cell Cycle (ECC) network

The ECC network model generated in this project was built to capture the signalling events driving entry into G1 and progression through the early cell cycle events, up to the G1-S phase transition. A detailed network schematic of the ECC’s biochemical reactions can be seen in **Figure 1**. We nominally include two Tyrosine Kinase Receptors (TKRs), the Insulin Receptor (INSR) and the Fibroblast Growth Factor Receptor (FGFR), as they strongly stimulate the PI3K and MAPK pathways respectively but are both capable of stimulating each pathway.

The two major mitogenic signalling pathways, PI3K and MAPK, were included for three reasons. The first is that they are known to potently stimulate the production of cyclin D, the primary activator of CDK4/6 (46, 47). The second is that activating mutations in these pathways are some of the most common mutations across all cancer types and the final reason is that these same mutations have been shown to facilitate resistance to CDK4/6 inhibitors (48, 49). The inclusion of proteins within this network is not exhaustive, due to computational limitations, but our model captures the core signalling relationships and network structure. The final component included in this model was the network regulating G1 entry and progression into S phase. This includes the relevant CDKs, cyclins and their inhibitors, as well as the transcription machinery regulating cyclin expression. Clinical approval for the use of CDK4/6 inhibitors has traditionally been limited to patients with ER+ (estrogen receptor-positive) breast cancer who were concurrently receiving ER (estrogen receptor) antagonists but were still experiencing disease progression (50–52). Due to the ER’s ability to promote CDK4/6 inhibition efficacy and the clinical practice of simultaneously prescribing ER antagonists with CDK4/6 inhibitors (50–52), the ER and its downstream signalling events were also included in the model.

The ECC network model was constructed using ODEs that represent biochemical interactions as a series of ordinary differential equations based on established kinetic laws. More detailed description of the model is given in the **Methods and Materials**. The full ODE equations and reaction rates are given in supplementary **Model Files S1 - S3**.

### The ECC network robustly facilitates resistance-associated protein dynamics

To investigate the meta-dynamics of the ECC network, i.e. the range of dynamics that can be facilitated by the topology of the ECC network, we first generated 100,000 unique model instances. Each model instance possessed an identical set of ICs along with a unique set of parameter values. The parameter values for each model instance were generated by randomly selecting values for each parameter between the range 10^-5^ to 10^4^. See supplementary **Figure S1A** for a visual representation of the model generation process. Each model instance was simulated as laid out in the *ODE model construction, modelling, and simulations* (see **Methods and Materials**), and the resulting time-course drug response of each of the network’s proteins was stored. This was repeated for all model instances. Alongside this process we also performed an experiment to test if novel, randomly drawn sets of model instances produced altered distributions of dynamics. Selecting new sets of 100,000 model instances produced near identical distributions of dynamics giving us confidence the distribution of protein dynamics converges at this set size. Details of this experiment can be found in the **Supplementary Information**.

We found that parameter variation resulted in a wide range of drug perturbation responses for many of the ECC network proteins. For example, **Figure 3A** displays a clustered subset of monophosphorylated Rb (pRb) time-course drug responses, showing a fairly extreme variety of drug-response dynamics that are possible in response to parameter variation. pRb not only possesses a *quantitative* spectrum of intra-category responses, which is probably expected, but it can also display a range of *qualitative* dynamics, and in fact, can demonstrate all six categories of qualitative drug response dynamics (see **Methods and Materials**).

**Figure 3:**
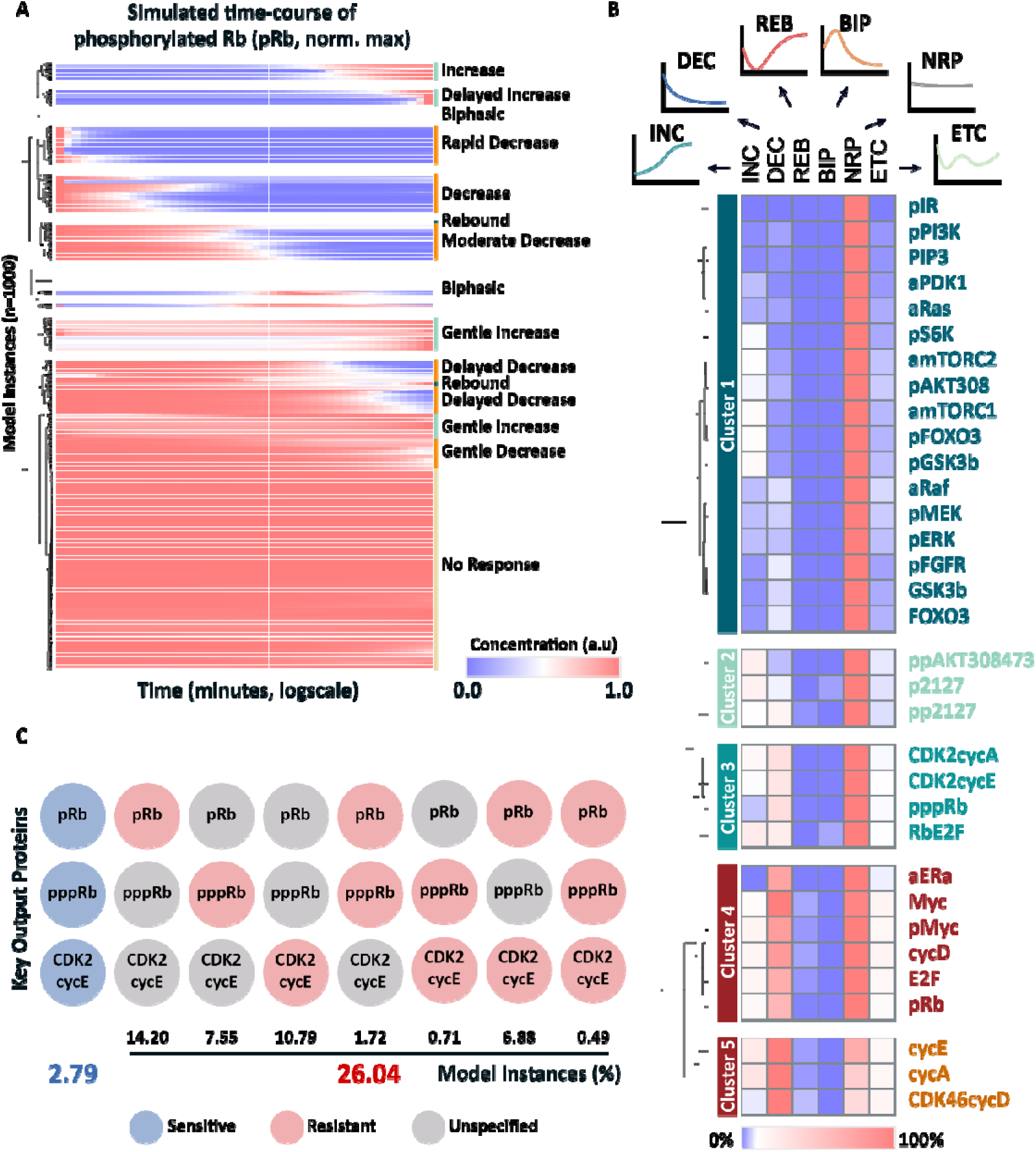
The ECC network robustly facilitates resistance-associated protein dynamics. **A)** Clustered heatmap of a the time-course profiles of a representative subgroup model instances, normalised to maximum value. Time-course profiles are of mono-phosphorylated Rb (pRb), the downstream target of active CDK4/6, and in response to CDK4/6 and ER inhibition. Blue represents low concentration and red represents high concentration. **B)** The frequency of the 6 dynamic categories for the model’s active protein forms, across 100,000 model instances. Blue represents low frequency and red represents high frequency. Species have been clustered to highlight species that have highly correlated distributions of dynamics. **C)** The frequency of model instances displaying simultaneous sensitivity (blue), or resistance in at least one of the key output proteins (red). Only 2.79% of model instances show simultaneous sensitivity across all three key output proteins. The percentage of resistant model instances was calculated by measuring the number of unique model instances that show resistance-associated dynamics for at least one of the key output proteins.

Categorising the dynamics of each protein across all of the model instances allowed us to calculate the distribution/frequency of each dynamic for each protein. The distribution of behaviours for the ‘active’ forms of the model’s protein species can be seen in the heatmap in **Figure 3B**. This overall distribution of protein behaviours is akin to a map that shows how the network state can affect the response of individual proteins to a drug treatment, in this case, the response to simultaneous CDK4/6 and ER inhibition. For many proteins, particularly proteins far upstream of the drug targets, the network state has only a little influence on drug response. This is particularly well demonstrated by the proteins in cluster 1, which show close to 100% no-response (NRP) dynamics.

Other proteins are much more strongly influenced by network state, where under the right conditions, a protein’s response can flip from sensitivity to resistance. The proteins in clusters 4 and 5 show this phenomenon to a moderate degree but also show a predilection towards decreasing dynamics. The predilection to a decreasing dynamic may be due to their proximity to the drug targets CDK4/6 and ER and/or the linearity of the interactions between these proteins and the direct drug targets. It is particularly interesting to observe that most of the major promoters of cell cycle progression downstream of CDK4/6 cluster together (cluster 3, **Figure 3B**) and are all strongly influenced by the network state. This suggests that heterogeneity can strongly influence whether CDK4/6 inhibition will translate to an effective network-level response and prevent cell cycle progression.

To further assess the capability of the ECC network to facilitate adaptive resistance, we dug into the dynamic distributions of the key output proteins: pRb, pppRb (hyper-phosphorylated Rb) and CDK2cycE. Recall that we have defined resistance dynamics as increasing, biphasic and rebound, and sensitive behaviours as decreasing. While the frequency of sensitive dynamics was much higher than even the sum of resistance dynamics for all three proteins, the ratio of model instances displaying sensitivity to those displaying resistance was much lower than we expected. The ratios of sensitive to resistant behaviours, as percentages of 100,000 model instances, were 30:14, 22:8, 19:11 (rounded) for pRb, pppRb and CDK2cycE, respectively (see supplementary **Figure S2A**). These results suggest that for every 2 model instances/network states that facilitate sensitivity, there is 1 that facilitates resistance.

We then analysed the model instances further to identify how frequently sensitive and resistant behaviours occurred simultaneously across the three key output proteins. Our results suggest that broad sensitivity across all three output proteins is quite rare, and that resistance is remarkably common (**Figure 3C)**. We observed that only 2.79% of model instances show simultaneous sensitivity across all three output proteins (**Figure 3C)**. In contrast, when evaluating model instances where at least 1 of the output proteins displays a resistance associated dynamic, we observe that this occurs in just over a quarter, 26.04%, of all model instances (**Figure 3C)**.

Given that CDK46cycD is only strongly suppressed in just under 60% of the model instances (supplementary **Figure 2A)**, we hypothesised that the resistance displayed by the key output proteins may be largely due to insufficient suppression of CDK46cycD. To explore this, we filtered out the model instances that showed strong suppression of CDK46cycD and re-generated the distribution of protein dynamics. The resulting heatmap can be seen in supplementary **Figure S2B**. Surprisingly, the distribution of behaviours did not appear to change significantly, with the key output proteins showing very similar levels of adaptive resistance dynamics. This result suggests that the resistance dynamics of the key output proteins are *bona fide* adaptive resistance mechanisms and not just a result of insufficient suppression of the drug target.

Taken together, these results show that in the face of heterogeneity, the ECC network is robustly capable of facilitating adaptive resistance and that key output proteins display adaptive resistance even when the drug targets are being robustly suppressed.

### Reaction coefficients are a stronger driver of adaptive-resistance dynamics than protein abundance

Having investigated the influence of parametric variation on drug response, we then wanted to explore the effect of IC variation in a similar fashion. To this end, we generated a further 100,000 model instances. This time, however, each model instance possessed an identical ‘nominal’ parameter set (see supplementary **Table S1)**, along with a unique set of ICs. Each unique set of ICs was generated by randomly selecting values between the range 10^0^ and 10^4^, for the inactive form of each protein. See supplementary **Figure 1B** for a visual representation of the model generation process. We then repeated the simulation and analysis of protein dynamics as performed previously. The resulting distribution of dynamics can be seen in supplementary **Figure S3A**, and the clustering of this distribution can be seen in supplementary **Figure S3B**.

Interestingly, we found that compared to parametric variation, IC variation induced much less variety in protein response dynamics. Most proteins demonstrated a singular dominant behaviour and very few proteins exhibited both sensitive and resistance dynamics. This seems to suggest that IC variation is less capable of influencing a proteins drug response. However, we thought that this may have been due to the selected nominal parameter set and may not be a universal phenomenon. To explore this hypothesis, we investigated the ability of parametric variation and IC variation to induce adaptive resistance in our key output proteins. Essentially, we selected sensitive model instances, then randomly varied either parameter values or ICs and measured how frequently the model instance switched from drug-sensitive to drug-resistant.

To achieve this, we first selected 1000 unique parameter sets and 1000 unique sets of ICs that demonstrated a decreasing dynamic for one of our key output proteins, e.g. pppRb. Each parameter set was combined with each IC set, resulting in the creation of 1,000,000 unique model instances. These model instances were arranged in a 1000 x 1000 matrix, where the rows represent the unique parameter sets and the columns represent the unique ICs. See supplementary **Figure 1C** for a visual representation. Each model instance was simulated, and the dynamic of the key output protein recorded in the matrix. By measuring the frequency of resistant dynamics across the rows and columns, we could compare the ability of parameter variation and IC variation to induce resistance. An overview of this process can be seen in **Figure 4A**.

**Figure 4:**
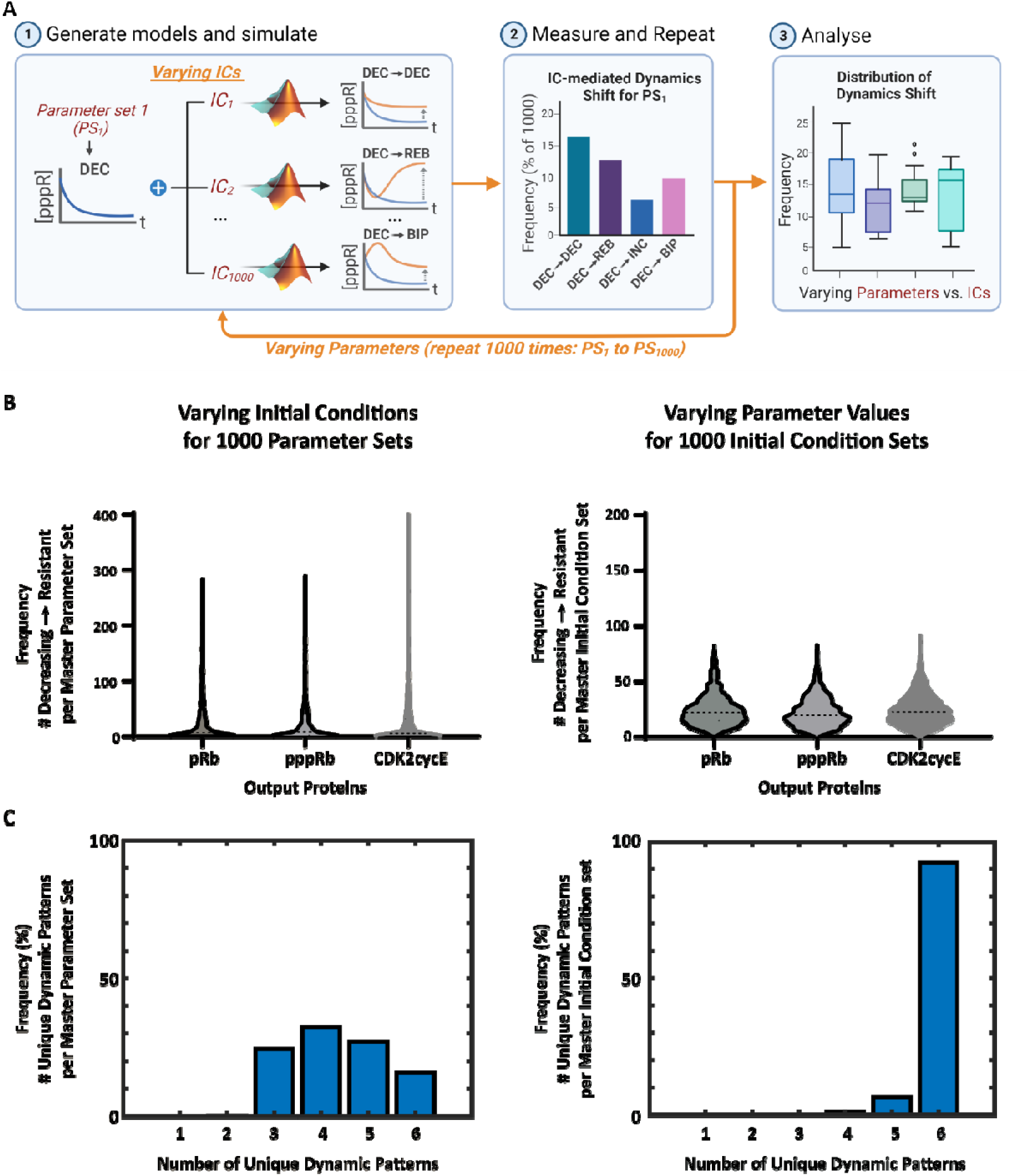
Reaction coefficients are a stronger driver of adaptive-resistance dynamics than protein abundance. **A)** Overview of the computational pipeline undertaken to compare the effects of parametric variation with initial condition variation. 1000 unique initial condition sets are applied to a base parameter set that demonstrates a decreasing dynamic for the key output protein being analysed. Each of the new model instances are simulated and the category of the output protein’s dynamic is determined. Then the number of times the model instances produce a resistance-associated dynamic is measured. This entire process is repeated for 1000 base parameter sets. The final step is measuring the distribution of resistance behaviours when the parameter values are changed and comparing the distribution when the initial condition values are changed. **B)** The frequency of which initial condition variation (left) induces resistance, in each key output protein. Varying initial condition values usually does not induce resistance, however, for select parameter sets, most initial conditions produce resistance associated dynamics. This analysis is repeated for parameter variation (right). Contrary to varying initial condition values, varying parameters usually induces resistance to a small degree, but there are no sets of initial conditions that are universally resistant. **C)** The number of unique dynamic categories produced by initial condition variation (left), versus parametric variation (right). Varying initial conditions produces a moderate variety of qualitative dynamics, and can produce all 6 dynamic categories, but varying parameters almost always induces every single possible dynamic category for the key output proteins.

We found that for most parameter sets, varying the initial conditions *never* induced resistance in any of the three output proteins (**Figure 4B**, left). There was however, a very small number of parameter sets where altering the initial conditions frequently induced resistance. In contrast, altering the parameter values *always* induced resistance in a small number of the IC sets; but there were no IC sets wherein parameter variation strongly induced resistance, independent of the protein chosen (**Figure 4B**, right).

To further characterise the difference between parameter and IC variation, we then investigated the variety of dynamics induced by either parameter or IC variation. For each master parameter or IC set, we measured how many unique dynamic categories were produced across their respective 1000 model instances. We found that varying the ICs still induced a moderate amount of variety in protein dynamics (**Figure 4C**, left). However, parameter variation almost always produced all six dynamic categories (**Figure 4C**, right).

These results provide clear evidence that changes in both parameter values and initial condition values can facilitate the emergence of adaptive resistance. Because conserved totals shift the steady-state values even with fixed kinetics, expression heterogeneity (via totals) can alter pre-treatment equilibria and subsequent drug responses. Supplementary **Figure S4** demonstrates this shift. However, changes in parameter values appear to be much more capable of inducing large, qualitative shifts in protein dynamics compared to changes in initial condition values.

The ability of expression variation to induce resistance also seems to be dependent on the master parameter set and there also appears to be no IC set that can induce resistance independent of the parameter values. These results imply that heterogeneity in reaction coefficients is much more likely, and independently able to facilitate the emergence of adaptive resistance compared to heterogeneity in protein abundance. Due to the *independent* ability of parameter variation to *strongly* induce adaptive resistance, we chose to focus on this particular form of heterogeneity in the following studies.

### Reaction coefficients co-ordinate to drive adaptive resistance, independent of their individual strengths

The data produced by our initial MDN pipeline enabled us to identify thousands of model instances where the key output proteins displayed adaptive resistance dynamics. Using these model instances, we next wanted to explore whether there were shared network features driving resistance, or if each model instance was unique in its ability to generate resistance. To explore this in a systematic manner, we extracted subsets of model instances, one for each key output protein and resistance behaviour combination (i.e. protein-dynamic combination) and performed a series of analyses on their parameters.

First, we investigated the mean and standard deviation of each parameter in each protein-dynamic combination to see if there were any individual parameter values underpinning resistance. We found that the mean of most parameters was between 0.1 and 1, but the standard deviation usually spanned 6 orders of magnitude (supplementary **Figure S5)**. We did observe that all three receptor-activation associated parameters (kc1f1-INSR, kc10f1-FGFR, kc15f1-ER) were generally much lower in value than the rest, but further investigation ruled that this was generally due to higher values causing the model’s system of ODEs to become too stiff, and therefore unsolvable, and were thus filtered out. We then undertook hierarchical clustering of each protein-dynamic combination’s parameter values to try and identify if there were groups of parameter values driving resistance. However, we were unable to find any meaningful clusters (supplementary **Figure S6)**. Together, these results led us to conclude that there were no particular individual parameters or groups of parameter values that were responsible for driving resistance.

Having found no patterns in the raw, or absolute, parameter values, we decided to investigate how frequently each parameter contributed to the increase or recovery of protein activity following drug perturbation, for each output protein. To this end, we investigated the effects of parameter knockdowns on the following dynamic features: maximum concentration, final concentration and/or rebound, hereafter collectively referred to as ‘resistance features’. These features were chosen as they represent either an increase or recovery of protein activity, post drug perturbation, that could potentially enable a cell to overcome the drug insult. The relevant resistance features for biphasic and increasing dynamics were maximum concentration and final concentration, and the relevant features for rebound dynamics were rebound and final concentration. Rebound is not applicable to biphasic and increasing dynamics as it simply hasn’t occurred, and the maximum concentration is only applicable to rebounding dynamics if the rebound overshoots the initial concentration, which is then covered by final concentration.

To measure the influence of parameters on the resistance features of the key output proteins we performed a high-throughput perturbation analysis wherein we reduced the value of each parameter by 20% in the third simulation phase i.e. the drug treatment phase. Parameters were perturbed one parameter at a time, and we measured the effect on the output protein’s resistance features. A breakdown of the measured dynamic features, including the resistance features just described, is given in supplementary **Figure S7**.

If the knockdown of a parameter resulted in a reduction in the resistance features of the output proteins’ dynamic, it scored a 1; otherwise it scored a 0. Repeating this process for each parameter and model instance resulted in an M x N matrix, where M is the number of model instances (protein and dynamic dependent), and N is the number of perturbed parameters (94), for each protein-dynamic combination. See **Figure 5A** for a visual representation of this, and the remaining overall process.

**Figure 5:**
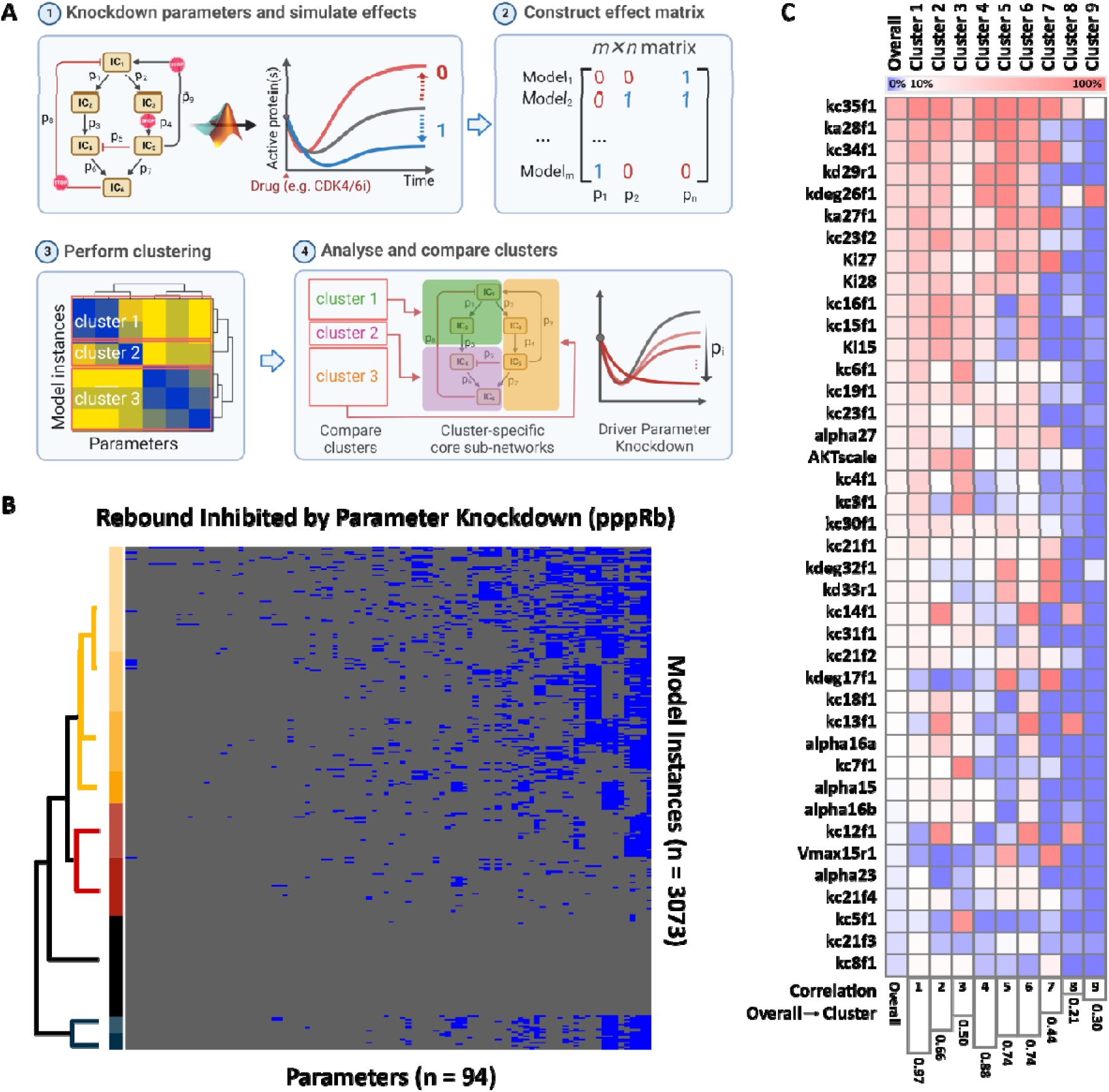
MDN analysis identifies core sub-networks that facilitate resistance. **A)** Overview of the process undertaken to investigate the relationship between parameter knockdown and resistance-dynamic features to identify resistance driving parameter signatures. Each parameter is knocked down by 20%, one at a time, and the effect on the protein dynamics is measured. If the knockdown decreases resistance it scores a 1, otherwise it scores a 0. This process is repeated for each parameter, one at a time, for each model instance. The end result is a matrix where the columns represent parameters, and the rows represent the model instances. This matrix can then be analysed using clustering methods to identify parametric resistance signatures. **B)** Heatmap produced by hierarchical clustering of the parameters that contribute to rebounding pppRb, for model instances that display rebounding pppRb. **C)** Comparing parametric resistance signatures with the parameters that most frequently contribute to resistance overall. Parameters are ranked by how frequently they contribute to rebounding pppRb. Clusters are aligned with overall ranking. The lower bar graph represents the correlation between the overall ranking and each cluster specific ranking. This heatmap shows that some parametric resistance signatures align closely with the overall ranking, but some clusters are quite different, emphasising the context specificity of resistance.

Initially, we wanted to determine how frequently each parameter contributed to the resistance features of each output protein. To achieve this, we summed up the number of times a parameter scored a 1 in the preceding analysis across all model instances, for each protein-dynamic combination. The results were plotted as bar graphs (top and bottom 10 parameters), which can be seen in supplementary **Figure S8**. The graphs showed clear rankings in how frequently each parameter contributed to the adaptive resistance features of each protein dynamic. Each protein-dynamic combination displayed a unique ranking but there also appeared to be a significant overlap in the top-scoring parameters. There also appeared to be more intra-protein overlap than inter-protein overlap, and pppRb and CDK2cycE shared much more similarity with each other than either with pRb. It was also interesting to observe that for biphasic pppRb and CDK2cycE, their top-scoring parameters accounted for approximately 90% of all model instances, indicating that these parameters must be critical in driving this particular dynamic for these proteins. The remaining top scoring parameters for each protein-dynamic subset only accounted for between 50-80% of model instances, suggesting each protein-dynamic combination possesses a degree of heterogeneity in the parameters that drive their adaptive resistance dynamics.

Together, the above results show that it is not the value of individual parameters that drive adaptive resistance, but rather it is the manner in which parameters co-ordinate to shape the quantitative and qualitative features of a protein’s dynamic that drives adaptive resistance.

### MDN can identify a full spectrum of core sub-networks that facilitate resistance

The observation that the top-scoring parameters generally do not broadly drive adaptive resistance dynamics suggests that there may be a number of different network states that are capable of driving a given resistance dynamic. To investigate this possibility, we subjected the matrices produced in the previous analysis to hierarchical clustering to try and find groups of model instances with shared resistance-driving parameter signatures. As a prime example, we first focused on rebounding hyperphosphorylation of Rb; both as the hyperphosphorylation of Rb strongly promotes cell cycle progression, and because previous studies have demonstrated the role of this protein-dynamic combination in CDK4/6 inhibitor resistance (21, 53, 54). **Figure 5B** displays a heatmap showing how rebounding pppRb model instances cluster, with respect to the contribution of their parameters to the rebounding dynamic.

The clustering of the pppRb-rebound model instances produced 9 robust clusters, where robust clusters are defined as those that contain at least 1% of the total protein-dynamic combination model instances, and the top-scoring parameter in the cluster accounts for at least 80% of its constituent model instances. Simply put, the clusters accounted for a high enough proportion of the model instances and the top scoring parameter was robustly representative of the model instances within the cluster. The left-most column of the heatmap in **Figure 5C** shows the overall ranking for how frequently each parameter contributes to the rebound dynamic of pppRb, and the remaining 9 columns represent the 9 clusters, aligned with the overall ranking. We observed a fairly strong consensus amongst the top-scoring parameters, with the majority of the cluster differences coming from the middle and lower parameters (**Figure 5C**). Of note, this pattern appeared to be broadly true when we analysed all the 9 protein-dynamic groups (supplementary **Figure S9**).

Despite broad similarities between the clusters, many of the individual clusters did not correlate particularly well with the overall ranking and possessed clear differences in their resistance-driving parameter signatures (**Figure 5C**). These results suggest that the same drug response dynamic for the same protein in two different cells can be driven by different network states. To highlight the similarities and differences between the subnetworks underpinning resistance driven by pppRb rebound, we overlayed the parameter signatures of the five largest clusters on the ECC network schematic (**Figure 6A**). This revealed significant overlap between the clusters around the formation of the CDK2-cyclin E complex and the hyperphosphorylation of Rb itself. These results were to be expected as the CDK2-cyclin E complex is directly responsible for the hyperphosphorylation of Rb and confirms the logical hypothesis that CDK2 activation can drive the reactivation of Rb phosphorylation following CDK4/6 and ER inhibition. However, that almost all of the clusters shared this core network suggests that this subnetwork alone is likely sufficient to enable the reactivation of Rb phosphorylation on its own and would be strongly selected for by cancer during treatment. Similarly, but less expectedly, we also found strong overlap at the level of Myc (**Figure 6A**), suggesting a strong likelihood for its involvement in the development of resistance.

**Figure 6:**
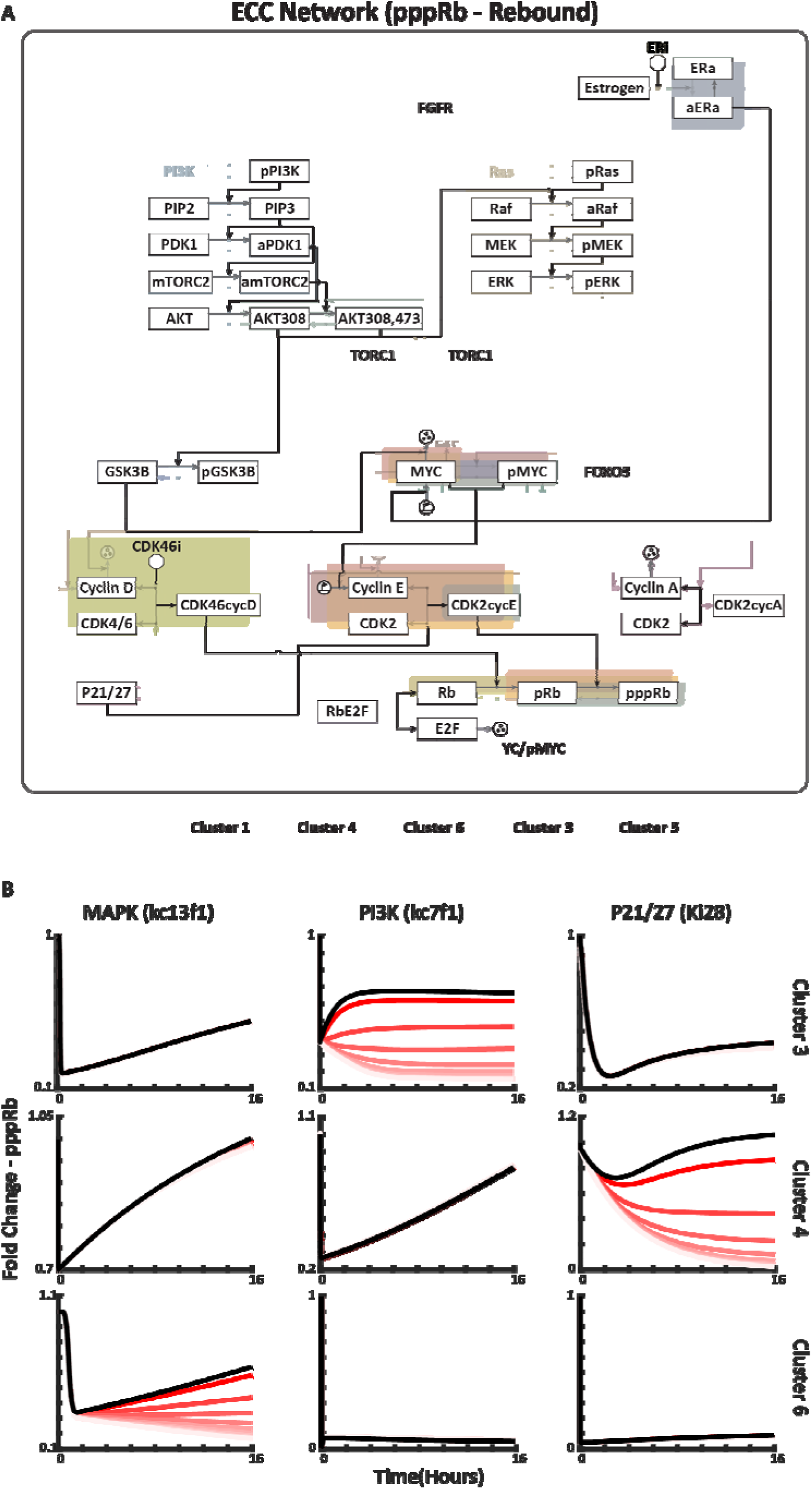
Identification of subnetworks driving pppRb rebound-mediated resistance within the ECC network. **A)** Cluster-specific parameter signatures overlaid with the ECC network to highlight subnetworks that drive resistance through rebound of pppRb. Only the five largest clusters (out of 9) are displayed. **B)** Parameter knockdowns for the top-scoring parameters in three select clusters that display divergent resistance driving parameter signatures. Black represents treatment with CDK4/6 and ER inhibitors alone, decreasingly-red lines represent the addition of an increasingly potent parameter knockdown. These results highlight the context specificity of parameter knockdown when attempting to prevent adaptive resistance.

On the other hand, clusters that have less overlap highlight potential differences in underlying resistance mechanisms between patients experiencing seemingly similar drug responses. For the pppRb-rebound group, these clusters included those driven by the PI3K and MAPK pathways, protein degradation regulation (GSK3B), and p21/27 inhibition of the CDK2-cyclin E complex (**Figure 6A**). Consistently, when we targeted (inhibited) parameters related to these signalling nodes across different clusters we did not observe universal suppression of pppRb; instead, we saw cluster-specific suppression of pppRb (**Figure 6B**).

Expanding this analysis to include all 9 protein-dynamic combinations, we observed that resistance-promoting subnetworks were widespread throughout the ECC network (supplementary **Figure S10** and **Figures S10A-I)**. Further, we identified just under 100 robust clusters/subnetworks capable of facilitating resistance dynamics (supplementary **Table S2A – J**). Finally, to highlight the overlap between protein-dynamic clusters, we combined all of the individual clusters of all of the protein-dynamic combinations (supplementary **Figure S11**).

This revealed resistance most frequently converges on the activation of CDK2. Mechanisms driving this activation include: the promotion of cyclin E synthesis, reduction in cyclin E degradation, promotion of the formation of the CDK2cyclin E complex and suppression of p21 and p27 mediated CDK2 inhibition. There were also moderate contributions to resistance spread throughout both upstream mitogenic pathways. The contribution of the ER to resistance seemed to be context dependent. A strong deactivation rate of the ER contributed to resistance quite frequently, yet paradoxically strong activation of the ER also somewhat frequently contributed.

The above results suggest three things. First, while there is a moderately large number of network states that can drive resistance to dual CDK4/6-ER inhibition, there is very likely only a **finite** number of network states that cancer can exploit to overcome drug perturbation.

Second, the moderately large number of network states that facilitate resistance and the observation that any given resistance dynamic can be driven by many different mechanisms highlight the low likelihood of finding broadly-applicable resistance mechanisms. Third, some protein nodes and interactions contribute far more frequently to resistance than others. It is quite possible that these frequencies are related to the likelihood and tumorigenicity of mutations affecting these nodes/interactions.

### Qualitative support for MDN modelling-based simulations and predictions Monoculture cell populations demonstrate a significant degree of heterogeneous signalling dynamics in response to CDK4/6 inhibitors

This study provides a theoretical framework that connects heterogeneity, protein signalling dynamics and adaptive resistance mechanisms. To validate this framework, we analysed the literature to pinpoint biological evidence for a number of our key predictions. First, we wanted to observe if and how much drug-response signalling heterogeneity exists in cellular monocultures. Second, we wanted to find evidence that cells possess differential resistance-mediating subnetworks. And third, we wanted to determine if the resistance-mediating subnetworks we identified agree with known resistance mechanisms.

Recently, Yang et al. constructed a novel reporter system that enabled them to investigate the activity of CDK4/6 and CDK2 in individual MCF10A cells in response to three clinically approved CDK4/6 inhibitors: abemaciclib, palbociclib and ribociclib (55). Plotting the single-cell time-course data for each drug and protein revealed that, at the single-cell level, there is a wide variety of protein signalling dynamics (**Figure 7A**). We then subjected this time-course data to our category analysis to calculate the distribution of each protein’s drug response dynamics (**Figure 7B**). We found that at the level of CDK4/6, the direct target of the three drugs, there was a moderate degree of signalling heterogeneity. Additionally, even the most potent drug (abemaciclib) failed to induce sustained suppression in approximately 10% of the cell population. The activity levels of CDK2, however, showed a much greater degree of signalling heterogeneity; and all three of the drugs only managed to strongly suppress CDK2 activity in a sustained manner in 40% of the population (**Figure 7B**). We further observed that there were many individual cells that demonstrated *increasing* CDK2 activity in response to CDK4/6 inhibition, a particularly unintuitive result, predicted by our MDN analysis. We then extracted a random sample of model instances from our MDN analysis (parametric variation) and plotted the drug response dynamics of CDK46cycD and CDK2cycE, our model’s equivalent active forms of CDK4/6 and CDK2 **(Figure 7C)**. The resemblance between the distribution of dynamics produced by our model simulations and the single-cell data is striking, both showing very similar proportions of resistant and sensitive dynamics.

**Figure 7.**
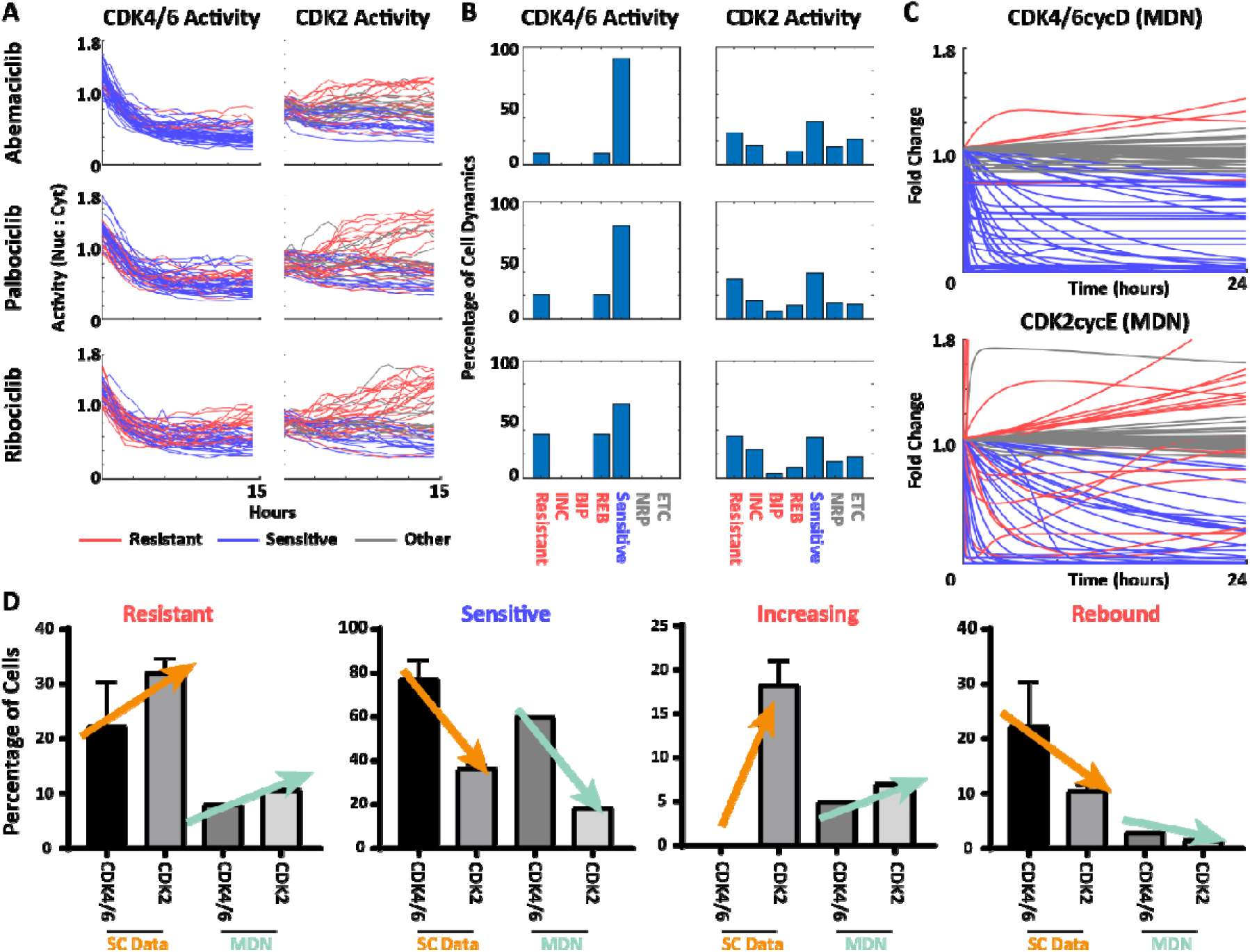
Validation of MDN-based predictions. Qualitative comparisons between single-cell data and our MDN analyses. **A)** Single-cell time-course responses to CDK4/6 inhibition from Yang et al. (55)). The reporter readout reflects phosphorylation-dependent nucleo-cytoplasmic shuttling; the y-axis is the nuclear/cytoplasmic fluorescence ratio (not fold-change to t=0). Baseline values at t=0 therefore vary across cells (typically ∼1–1.8) due to inherent differences in reporter localization prior to treatment. Red traces: resistance-associated dynamics; blue: sensitivity-associated; grey: no/limited response. **B)** Frequency of each dynamic category in panel A. **C)** Representative MDN simulations for CDK4/6·CycD and CDK2·CycE drawn from the model ensemble (parametric variation). Colours denote the same dynamic classes as in A. The distribution of resistant and sensitive dynamics from our simulations are highly correlated with the single-cell data. **D)** Agreement between single-cell data and MDN simulations quantified by the distribution of dynamic classes across both datasets.

Finally, we examined how the distribution of specific dynamics changed when comparing CDK4/6 activity with CDK2 activity. **Figure 7D** shows that the single-cell data and our MDN analysis exhibit the same general trends: there are more resistant dynamics at the level of CDK2 than at the level of CDK4/6; there are fewer sensitive dynamics at the level of CDK2 than CDK4/6; and there are more increasing dynamics at the level of CDK2. Perhaps most unintuitively, there are fewer rebound dynamics at the level of CDK2 in the single cell data, which was also predicted by our MDN analysis. Though this analysis is *qualitative*, we demonstrate the capacity for variation in protein activity within a conserved network architecture to drive a wide variety of signalling dynamics, including those associated with resistance, that mirrors the variation in signalling dynamics seen in isogenic single cells.

### The identified resistance-mediating core subnetworks align with known resistance mechanisms

Previous studies have identified a wealth of mechanisms responsible for driving resistance to CDK4/6 and ER inhibitors. Focusing on mechanisms that have overlap with our model, the vast majority of identified mechanisms align quite well with the predictions made by our MDN analysis, validating the ability of this technique to identify a full spectrum of resistance-driving mechanisms. Moreover, our analysis also makes predictions about additional protein nodes/interactions that have not yet been identified by the literature.

Known mechanisms of resistance to CDK4/6 inhibitors include: Rb loss/mutations (56, 57), CDK4/6 overexpression (58–60), loss of ER expression (58, 61, 62), loss of PTEN (63), reduced expression/activity of p21 and p27 (64–66), activation of mTOR complexes (67, 68), activation of PI3K and PI3K signalling pathways (61, 69), activation of PDK1 (70), upregulation of FGFR (71, 72), AKT amplification and/or over-activation (73), E2F amplification and/or over-activation (74), MAPK pathway activation (75–77), c-Myc activity (78, 79), cyclin E overexpression and increased CDK2-cyclin E complex formation (43, 44, 80, 81). Known mechanisms of resistance to ER inhibitors include mutations that induce the constitutive activation of ER (82, 83), loss of ER activity (84, 85), activation of PI3K and MAPK signalling (86, 87), overexpression of c-Myc (88, 89) and loss of p21 and p27 (90, 91).

The preceding list represents a wealth of known resistance mechanisms that are widely distributed throughout the ECC network. Every single one of the above mechanisms is represented in at least one of our identified resistance-driving subnetworks. To clearly demonstrate how well our MDN analysis captured the known resistance mechanisms we created a table containing the known resistance mechanisms and the respective resistance-driving parameters identified by our MDN analysis (supplementary **Table S3)**. The ability of MDN to largely capture the known resistance mechanisms highlights its potential usefulness in predicting resistance mechanisms to novel drugs and drug targets. Apart from capturing the known resistance mechanisms, our analysis really extends the observation that resistance can emerge from many different nodes within the ECC network and further adds support to the idea that it is critical to treat each patient individually.

## DISCUSSION

Heterogeneity is a defining feature of cancer and a major barrier to durable responses to targeted therapies. Clinical and experimental studies of CDK4/6 inhibitors in ER+ breast cancer consistently show wide inter- and intra-tumour variability in response and multiple, often co-existing resistance mechanisms (92). Yet, despite this wealth of descriptive data, we still lack mechanistic frameworks that can systematically relate underlying molecular heterogeneity to the spectrum of signalling behaviours that enable adaptive drug resistance. In this work, we introduce meta-dynamic network (MDN) modelling as such a framework. By sampling broad ensembles of kinetic parameters and protein abundances on a fixed network topology, MDN maps the full distribution of qualitative signalling dynamics (‘meta-dynamics’) accessible to a network architecture, and identifies which of these trajectories are compatible with sustained target inhibition versus adaptive escape.

Our first key finding is that even within a single, fixed early cell-cycle (ECC) network topology, variation in reaction kinetics and total protein abundances can generate a surprisingly rich repertoire of qualitative signalling behaviours. We use the term ‘meta-dynamics’ to denote the convergent distribution of these behaviours across a large ensemble of model instances, which represents a theoretical upper limit on what the network can manifest dynamically.

Importantly, this repertoire includes trajectories in which continuous CDK4/6 inhibition fails to maintain target suppression and downstream cell-cycle arrest - canonical signatures of adaptive resistance. This provides a concrete mechanistic link between molecular heterogeneity and the well-described phenomenon of rapid signalling rewiring under targeted therapy, in which feedback and crosstalk restore pathway activity despite continued drug exposure (18, 93).

We find that adaptive resistance can emerge from heterogeneity in both kinetic parameters and protein abundances, but not symmetrically. However, across the ECC network, we observe that perturbations to kinetic parameters (interaction strengths) are more potent and consistent drivers of adaptive resistance dynamics than changes in total protein abundance. This supports a hierarchical view of control: network topology and interaction strengths largely determine which qualitative behaviours are possible, while protein levels modulate how frequently particular regimes are accessed. That hierarchy is consistent with both systems-level perspectives on cellular heterogeneity (94) and parameter-sensitivity analyses in whole-cell and signalling models, which have shown that many distinct parameter sets can collapse onto a limited set of functional outputs (95).

Perhaps the most striking observation from our analysis is how rapidly the meta-dynamic distribution converges. Despite sweeping over 90 kinetic parameters and 50 total abundances across nine orders of magnitude, the distribution of qualitative response classes stabilises well before 100,000 accepted model instances. Doubling the ensemble size yields only negligible changes in the frequencies of each dynamic regime, see supplementary **Figure S12**. This reflects a strong filtering effect imposed by network topology rather than simply a statistical inevitability of sampling from fixed priors. When parameters or abundances are pushed to extremes, nodes tend to be driven into saturating states, either effectively ‘always off’ or ‘always on’, and become unresponsive to upstream perturbations. These saturated cases fall into the ‘no response’ category and do not generate new, more complex temporal patterns. Consequently, the ECC topology funnels the near-infinite combinatorial parameter space into a finite, recurring set of qualitative behaviours (e.g. monotonic decrease, rebound, biphasic).

Such topology-constrained behavioural repertoires have also been noted in recent work that treats heterogeneity by assigning probability distributions to model parameters and analysing the resulting ensemble of single-cell behaviours (96, 97). MDN complements these approaches by emphasising qualitative dynamic classes and their frequencies rather than precise parameter posteriors.

At the level of individual nodes and subnetworks, our perturbation analyses reveal that adaptive resistance is a genuinely network-level property. Nearly every node in the ECC model can be implicated in at least one resistance subnetwork when viewed across the full heterogeneity ensemble, echoing the clinical and experimental literature where numerous, sometimes mutually exclusive mechanisms of CDK4/6 inhibitor resistance have been described - ranging from loss of RB1, to cyclin E/CDK2 activation, PI3K–AKT upregulation, FGFR signalling, and alterations in cell-cycle checkpoint control (92). However, MDN does not predict an unstructured ‘anything goes’ landscape. When we rank interactions by how frequently their perturbation participates in resistance-associated trajectories, a small set of hubs and co-regulated modules emerges. In other words, many routes to resistance exist, but they tend to be coordinated through a limited number of recurrent network motifs and subnetworks. This modular picture is consistent with reviews of adaptive resistance in breast cancer, which emphasise convergent rewiring of signalling hubs and feedback nodes despite diverse upstream lesions (18).

As an additional, orthogonal validation of the resistance subnetworks identified by MDN, we compared our perturbation-derived resistance modules with gene-dependency patterns from the Cancer Dependency Map (DepMap) (98). Although DepMap is not specific to CDK4/6 inhibition, hierarchical clustering of knockout dependencies for ECC-network genes revealed distinct vulnerability modules across >1000 cell lines (supplementary **Figure S13**). The existence of such modules supports the MDN prediction that different cellular contexts rely on distinct but recurrent subnetworks to sustain proliferation, reinforcing the view that resistance is governed by coordinated network-level processes rather than isolated nodes.

Our findings also shed light on why CDK4/6 inhibition resistance is so pervasive yet difficult to capture with single biomarkers. Recent work integrating bulk, single-cell, and trial data has shown that tumours resistant to CDK4/6 inhibition are characterised by increased intra-tumoural heterogeneity and multiple, co-existing resistance signatures, including MYC-driven programmes and altered estrogen-response pathways (99). Single-cell imaging and modelling studies have further demonstrated that cell-cycle dynamics and signalling trajectories at the single-cell level are critical determinants of CDK4/6 inhibitor sensitivity (100). Our MDN analysis provides a mechanistic underpinning for these observations: across a broad space of plausible network states, a surprisingly large fraction of configurations yield adaptive resistance-like signalling for key G1-S regulators, and simultaneous durable suppression of all relevant downstream effectors is rare. In such a landscape, static measurements of one or a few nodes are unlikely to robustly stratify patients, because resistance is inherently encoded in distributed, dynamic network behaviour rather than in single static markers.

Despite the theoretical nature of our study, qualitative comparison with single-cell signalling data supports the core results of MDN. Experiments in isogenic breast epithelial cells have shown that CDK4/6 activity reporters display highly heterogeneous dynamics under inhibitor treatment, with some cells remaining durably suppressed, others partially rebounding, and others showing minimal inhibition (55, 100). Our meta-dynamic distributions reproduce this spectrum of behaviours despite being generated from a much broader parameter ensemble than any one cell line is likely to occupy. We emphasise that this comparison is intentionally qualitative: our goal is not to fit a specific cell line with a calibrated model, but to demonstrate that the ECC topology, when subjected to realistic heterogeneity, naturally gives rise to the types of signalling trajectories observed empirically. The close qualitative agreement between simulated and experimental distributions suggests that even nominally isogenic populations may explore a substantial portion of the network’s meta-dynamic landscape.

As with any model-based framework, MDN has limitations that shape how its predictions should be interpreted. First, we use mass-action and first-order kinetics rather than more detailed Michaelis–Menten or cooperative rate laws. This was essential to keep the parameter space tractable for large-scale exploration, but it means that behaviours relying critically on enzyme saturation or higher-order cooperativity may be under-represented. Second, our parameter and abundance priors are deliberately broad and largely uniform in log-space, whereas biochemical parameters and expression levels in real cells are often better approximated by log-normal or other structured distributions (101–103). In this sense, the present work should be viewed as defining what is possible for a given topology rather than estimating what is probable in a specific tumour context. Third, our model is deterministic and does not explicitly capture intrinsic biochemical noise. We argue that sampling across a broad ensemble of parameter and abundance sets can approximate the time-averaged consequences of genetic, epigenetic, and stochastic variation, but intrinsic noise at low copy numbers and rare event dynamics will require explicit stochastic formulations (104). Finally, the ECC model abstracts some multi-protein modules into single nodes to maintain computational tractability. As a result, highly ranked “hub” nodes in our analysis should be interpreted as implicating critical processes or modules, rather than single gene products, as the key levers of resistance.

These limitations point naturally to several avenues for future work. A priority is to constrain MDN ensembles using experimentally inferred parameter and abundance distributions, for example by integrating Bayesian parameter-inference methods that represent cell-to-cell variability as probability distributions over model parameters. Another key direction is to couple MDN-derived resistance subnetworks with data-driven approaches that operate directly on patient-derived single-cell and spatial multi-omics data, which are increasingly used to dissect sample-level heterogeneity and its association with treatment response (105). Such integrated frameworks could bridge the gap between mechanistic prediction of resistance-enabling subnetworks and empirical identification of patient-specific vulnerabilities. On the dynamical side, embedding stochastic simulations or Langevin approximations within the MDN framework would allow us to evaluate how intrinsic noise reshapes the meta-dynamic landscape and whether it introduces qualitatively new resistance regimes or primarily modulates the frequencies of existing ones. Finally, our perturbation analyses suggest that a finite set of recurrent resistance modules underpins a vast number of resistant states. This aligns with translational efforts that design rational combination therapies to pre-empt or overcome adaptive resistance by co-targeting signalling hubs and compensatory feedbacks (106) and MDN could provide a principled way to enumerate and prioritise such combinations for experimental testing.

In summary, MDN modelling offers a systems-level framework that connects molecular heterogeneity to dynamic signalling behaviour and, ultimately, to adaptive drug resistance. By revealing how a fixed network topology can generate a finite but diverse set of resistance-associated trajectories, and by identifying the hubs and modules that recurrently mediate these trajectories, our work complements empirical single-cell and clinical studies and suggests concrete strategies for network-level intervention. Although we have focused on the G1-S transition and CDK4/6 inhibition as a clinically important case study, the MDN approach is broadly applicable to other signalling networks and therapeutic targets. As mechanistic models, single-cell datasets, and computational methods continue to mature, we envisage MDN-like frameworks playing an increasingly important role in predicting how tumours will escape targeted treatments and in guiding combination strategies to keep them one step ahead.

## Supporting information

Model File 1

Model File 2

Model File 3

Model File 4

## ACKNOWLEDGMENTS

L.K.N was funded by a Victorian Cancer Agency Mid-Career Research Fellowship (MCRF18026); a Venture Grant from Cancer Council Victoria, Australia; and an Investigator Initiated Research grant from the National Breast Cancer Foundation and Love Your Sister, Australia (IIRS-20–094). A.H was This research is/was supported by an Australian Government Research Training Program (RTP) Scholarship. This research was supported in part by the Australian Research Council Centre of Excellence for the Mathematical Analysis of Cellular Systems (MACSYS, CE230100001), funded by the Australian Government

## AUTHOR CONTRIBUTIONS

Conceptualization: Anthony Hart, Lan K. Nguyen.

Formal analysis: Anthony Hart, Sung-Young Shin, Lan K. Nguyen. Funding acquisition: Lan K. Nguyen.

Project administration: Lan K. Nguyen. Supervision: Lan K. Nguyen.

Writing – original draft: Anthony Hart, Lan K. Nguyen, Sung-Young Shin.

## Notes

### Competing Interest Statement

The authors have declared no competing interest.

### Summary of Updates

At a high level, we clarified the study design and analysis goals, tightened definitions, and added methodological detail where it most advances interpretability. Importantly, these updates leave the analytical pipelines and major conclusions unchanged. Conceptually, we now make explicit that our objective is coverage of the output space of qualitative dynamics supported by the network topology, not exhaustive enumeration of parameter space. To support this, we added a convergence analysis and clarified that triplicates refers to independent ensembles used to demonstrate reproducibility. We also refined how we describe and implement initial conditions (as conserved total abundances that encode expression heterogeneity) and reframed filtering as minimal numerical/feasibility checks, using rejection sampling to obtain the prespecified ensemble size. Solver choices and input modelling (constant step mitogen/drug) are now spelled out succinctly. We expanded the model specification and rationale (complete reaction list with rate laws and brief biological justifications in the Supplement) and unified terminology throughout. Figures and legends have been overhauled for readability and accuracy, with missing labels added and ordering corrected. For validation, we clarified the nature of the single-cell reporter readout, improved Figure 7s presentation, and emphasised - consistent with our aims - that comparisons are qualitative. Finally, we have rewritten the Discussion to centre on interpretation, implications, and connect our findings to the literature. It now: (i) frames MDN as a systems-level framework that links molecular heterogeneity to qualitative signalling meta-dynamics and adaptive escape under constant drug pressure; (ii) highlights two key findings: an asymmetry in control (interaction kinetics exert stronger, more consistent influence than protein abundance) and a topology-driven convergence whereby a vast parameter space funnels into a finite set of recurrent behaviours; (iii) shows that resistance is a network-level property, with many possible routes but a small set of recurrent hubs/modules dominating; and (iv) provides a qualitative alignment with single-cell reporter data while clarifying the intent and limits of that comparison. Moreover, we now explicitly discuss limitations (rate-law simplifications, broad priors, determinism, and modular abstractions) and outline next steps for future research, including data-constrained priors and stochastic extensions.

## REFERENCES

1. Vasan N, Baselga J, Hyman DM. 2019. A view on drug resistance in cancer. Nature 575:299–309.

2. Wang X, Zhang H, Chen X. 2019. Drug resistance and combating drug resistance in cancer. Cancer Drug Resistance 2:141.

3. Pich O, Bailey C, Watkins TB, Zaccaria S, Jamal-Hanjani M, Swanton C. 2022. The translational challenges of precision oncology. Cancer Cell.

4. Knudsen ES, Kumarasamy V, Nambiar R, Pearson JD, Vail P, Rosenheck H, Wang J, Eng K, Bremner R, Schramek D, Rubin SM, Welm AL, Witkiewicz AK. 2022. CDK/cyclin dependencies define extreme cancer cell-cycle heterogeneity and collateral vulnerabilities. Cell Reports 38:110448.

5. Jubran MR, Vilenski D, Flashner-Abramson E, Shnaider E, Vasudevan S, Rubinstein AM, Meirovitz A, Sharon S, Polak D, Kravchenko-Balasha N. 2022. Overcoming resistance to EGFR monotherapy in HNSCC by identification and inhibition of individualized cancer processes. Theranostics 12:1204–1219.

6. Dagogo-Jack I, Shaw AT. 2018. Tumour heterogeneity and resistance to cancer therapies. Nat Rev Clin Oncol 15:81–94.

7. Turke AB, Zejnullahu K, Wu Y-L, Song Y, Dias-Santagata D, Lifshits E, Toschi L, Rogers A, Mok T, Sequist L. 2010. Preexistence and clonal selection of MET amplification in EGFR mutant NSCLC. Cancer cell 17:77–88.

8. Patel AG, Chen X, Huang X, Clay MR, Komorova N, Krasin MJ, Pappo A, Tillman H, Orr BA, McEvoy J. 2022. The myogenesis program drives clonal selection and drug resistance in rhabdomyosarcoma. Developmental Cell 57:1226–1240. e8.

9. Cassidy J. 2019. Studying the clonal origins of drug resistance in human breast cancersUniversity of Cambridge.

10. Herrera-Abreu MT, Palafox M, Asghar U, Rivas MA, Cutts RJ, Garcia-Murillas I, Pearson A, Guzman M, Rodriguez O, Grueso J, Bellet M, Cortés J, Elliott R, Pancholi S, Baselga J, Dowsett M, Martin LA, Turner NC, Serra V. 2016. Early Adaptation and Acquired Resistance to CDK4/6 Inhibition in Estrogen Receptor-Positive Breast Cancer. Cancer Res 76:2301–13.

11. Ahmed TA, Adamopoulos C, Karoulia Z, Wu X, Sachidanandam R, Aaronson SA, Poulikakos PI. 2019. SHP2 Drives Adaptive Resistance to ERK Signaling Inhibition in Molecularly Defined Subsets of ERK-Dependent Tumors. Cell Rep 26:65–78.e5.

12. Blombery P, Anderson MA, Gong JN, Thijssen R, Birkinshaw RW, Thompson ER, Teh CE, Nguyen T, Xu Z, Flensburg C, Lew TE, Majewski IJ, Gray DHD, Westerman DA, Tam CS, Seymour JF, Czabotar PE, Huang DCS, Roberts AW. 2019. Acquisition of the Recurrent Gly101Val Mutation in BCL2 Confers Resistance to Venetoclax in Patients with Progressive Chronic Lymphocytic Leukemia. Cancer Discov 9:342–353.

13. Yun CH, Mengwasser KE, Toms AV, Woo MS, Greulich H, Wong KK, Meyerson M, Eck MJ. 2008. The T790M mutation in EGFR kinase causes drug resistance by increasing the affinity for ATP. Proc Natl Acad Sci U S A 105:2070–5.

14. Xue X, Liang X-J. 2012. Overcoming drug efflux-based multidrug resistance in cancer with nanotechnology. Chinese journal of cancer 31:100.

15. Smyth MJ, Krasovskis E, Sutton VR, Johnstone RW. 1998. The drug efflux protein, P-glycoprotein, additionally protects drug-resistant tumor cells from multiple forms of caspase-dependent apoptosis. Proceedings of the National Academy of Sciences 95:7024–7029.

16. Jin X, Ge LP, Li DQ, Shao ZM, Di GH, Xu XE, Jiang YZ. 2020. LncRNA TROJAN promotes proliferation and resistance to CDK4/6 inhibitor via CDK2 transcriptional activation in ER+ breast cancer. Mol Cancer 19:87.

17. Li Q, Jiang B, Guo J, Shao H, Del Priore IS, Chang Q, Kudo R, Li Z, Razavi P, Liu B, Boghossian AS, Rees MG, Ronan MM, Roth JA, Donovan KA, Palafox M, Reis-Filho JS, de Stanchina E, Fischer ES, Rosen N, Serra V, Koff A, Chodera JD, Gray NS, Chandarlapaty S. 2022. INK4 Tumor Suppressor Proteins Mediate Resistance to CDK4/6 Kinase Inhibitors. Cancer Discov 12:356–371.

18. Cremers CG, Nguyen LK. 2019. Network rewiring, adaptive resistance and combating strategies in breast cancer. Cancer Drug Resistance 2:1106.

19. Wright SCE, Vasilevski N, Serra V, Rodon J, Eichhorn PJA. 2021. Mechanisms of resistance to PI3K inhibitors in cancer: adaptive responses, drug tolerance and cellular plasticity. Cancers 13:1538.

20. Ma P, Fu Y, Chen M, Jing Y, Wu J, Li K, Shen Y, Gao J-X, Wang M, Zhao X. 2016. Adaptive and acquired resistance to EGFR inhibitors converge on the MAPK pathway. Theranostics 6:1232.

21. Herrera-Abreu MT, Palafox M, Asghar U, Rivas MA, Cutts RJ, Garcia-Murillas I, Pearson A, Guzman M, Rodriguez O, Grueso J. 2016. Early Adaptation and Acquired Resistance to CDK4/6 Inhibition in Estrogen Receptor–Positive Breast CancerEarly Adaption and Acquired Palbociclib Resistance. Cancer research 76:2301–2313.

22. Park S-M, Hwang CY, Choi J, Joung CY, Cho K-H. 2020. Feedback analysis identifies a combination target for overcoming adaptive resistance to targeted cancer therapy. Oncogene 39:3803–3820.

23. Nguyen LK, Kholodenko BN. 2016. Feedback regulation in cell signalling: Lessons for cancer therapeutics. Seminars in Cell & Developmental Biology 50:85–94.

24. Bachmann J, Raue A, Schilling M, Becker V, Timmer J, Klingmüller U. 2012. Predictive mathematical models of cancer signalling pathways. Journal of Internal Medicine 271:155–165.

25. Shankar E, Weis MC, Avva J, Shukla S, Shukla M, Sreenath SN, Gupta S. 2019. Complex Systems Biology Approach in Connecting PI3K-Akt and NF-κB Pathways in Prostate Cancer. Cells 8:201.

26. Clarke MA, Fisher J. 2020. Executable cancer models: successes and challenges. Nature Reviews Cancer 20:343–354.

27. Altrock PM, Liu LL, Michor F. 2015. The mathematics of cancer: integrating quantitative models. Nature Reviews Cancer 15:730–745.

28. Ghomlaghi M, Hart A, Hoang N, Shin S, Nguyen LK. 2021. Feedback, crosstalk and competition: ingredients for emergent non-linear behaviour in the PI3K/mTOR signalling network. International journal of molecular sciences 22:6944.

29. Tyson JJ, Laomettachit T, Kraikivski P. 2019. Modeling the dynamic behavior of biochemical regulatory networks. Journal of theoretical biology 462:514–527.

30. Tandon G, Yadav S, Kaur S. 2022. Chapter 24 - Pathway modeling and simulation analysis, p 409-423. In Singh DB, Pathak RK (ed), Bioinformatics 10.1016/B978-0-323-89775-4.00007-9. Academic Press.

31. Deuflhard P, Röblitz S. 2015. ODE Models for Systems Biological Networks, p 1-32. *In* Deuflhard P, Röblitz S (ed), A Guide to Numerical Modelling in Systems Biology doi:10.1007/978-3-319-20059-0_1. Springer International Publishing, Cham.

32. Kearney AL, Norris DM, Ghomlaghi M, Wong MKL, Humphrey SJ, Carroll L, Yang G, Cooke KC, Yang P, Geddes TA. 2021. Akt phosphorylates insulin receptor substrate to limit PI3K-mediated PIP3 synthesis. Elife 10:e66942.

33. Shin SY, Müller AK, Verma N, Lev S, Nguyen LK. 2018. Systems modelling of the EGFR-PYK2-c-Met interaction network predicts and prioritizes synergistic drug combinations for triple-negative breast cancer. PLoS Comput Biol 14:e1006192.

34. Sun X, Hu B. 2017. Mathematical modeling and computational prediction of cancer drug resistance. Briefings in Bioinformatics 19:1382–1399.

35. Chisholm RH, Lorenzi T, Clairambault J. 2016. Cell population heterogeneity and evolution towards drug resistance in cancer: biological and mathematical assessment, theoretical treatment optimisation. Biochimica et Biophysica Acta (BBA)-General Subjects 1860:2627–2645.

36. Belkhir S, Thomas F, Roche B. 2021. Darwinian approaches for cancer treatment: benefits of mathematical modeling. Cancers 13:4448.

37. Huang B, Jia D, Feng J, Levine H, Onuchic JN, Lu M. 2018. RACIPE: a computational tool for modeling gene regulatory circuits using randomization. BMC Syst Biol 12:74.

38. Städter P, Schälte Y, Schmiester L, Hasenauer J, Stapor PL. 2021. Benchmarking of numerical integration methods for ODE models of biological systems. Scientific Reports 11:2696.

39. Resat H, Petzold L, Pettigrew MF. 2009. Kinetic modeling of biological systems. Methods Mol Biol 541:311–35.

40. Stallaert W, Kedziora KM, Taylor CD, Zikry TM, Ranek JS, Sobon HK, Taylor SR, Young CL, Cook JG, Purvis JE. 2022. The structure of the human cell cycle. Cell Systems 13:230–240.e3.

41. Tyson JJ, Novák B. 2015. Models in biology: lessons from modeling regulation of the eukaryotic cell cycle. BMC Biology 13:46.

42. Iorio F, Knijnenburg TA, Vis DJ, Bignell GR, Menden MP, Schubert M, Aben N, Gonçalves E, Barthorpe S, Lightfoot H, Cokelaer T, Greninger P, van Dyk E, Chang H, de Silva H, Heyn H, Deng X, Egan RK, Liu Q, Mironenko T, Mitropoulos X, Richardson L, Wang J, Zhang T, Moran S, Sayols S, Soleimani M, Tamborero D, Lopez-Bigas N, Ross-Macdonald P, Esteller M, Gray NS, Haber DA, Stratton MR, Benes CH, Wessels LFA, Saez-Rodriguez J, McDermott U, Garnett MJ. 2016. A Landscape of Pharmacogenomic Interactions in Cancer. Cell 166:740–754.

43. Min A, Kim JE, Kim YJ, Lim JM, Kim S, Kim JW, Lee KH, Kim TY, Oh DY, Bang YJ, Im SA. 2018. Cyclin E overexpression confers resistance to the CDK4/6 specific inhibitor palbociclib in gastric cancer cells. Cancer Lett 430:123–132.

44. Taylor-Harding B, Aspuria PJ, Agadjanian H, Cheon DJ, Mizuno T, Greenberg D, Allen JR, Spurka L, Funari V, Spiteri E, Wang Q, Orsulic S, Walsh C, Karlan BY, Wiedemeyer WR. 2015. Cyclin E1 and RTK/RAS signaling drive CDK inhibitor resistance via activation of E2F and ETS. Oncotarget 6:696–714.

45. Ahmed TA, Adamopoulos C, Karoulia Z, Wu X, Sachidanandam R, Aaronson SA, Poulikakos PI. 2019. SHP2 drives adaptive resistance to ERK signaling inhibition in molecularly defined subsets of ERK-dependent tumors. Cell reports 26:65–78. e5.

46. Chen J, Bai M, Ning C, Xie B, Zhang J, Liao H, Xiong J, Tao X, Yan D, Xi X. 2016. Gankyrin facilitates follicle-stimulating hormone-driven ovarian cancer cell proliferation through the PI3K/AKT/HIF-1α/cyclin D1 pathway. Oncogene 35:2506–2517.

47. Wang H-Y, Yang S-L, Liang H-F, Li C-H. 2014. HBx protein promotes oval cell proliferation by up-regulation of cyclin D1 via activation of the MEK/ERK and PI3K/Akt pathways. International journal of molecular sciences 15:3507–3518.

48. Del Re M, Crucitta S, Lorenzini G, De Angelis C, Diodati L, Cavallero D, Bargagna I, Cinacchi P, Fratini B, Salvadori B. 2021. PI3K mutations detected in liquid biopsy are associated to reduced sensitivity to CDK4/6 inhibitors in metastatic breast cancer patients. Pharmacological Research 163:105241.

49. Samuels Y, Velculescu VE. 2004. Oncogenic mutations of PIK3CA in human cancers. Cell cycle 3:1221–1224.

50. FDA. 2017. https://www.fda.gov/drugs/resources-information-approved-drugs/palbociclib-ibrance#:∼:text=FDA%20granted%20palbociclib%20accelerated%20approval,based%20therapy%20in%20postmenopausal%20women., *on* FDA. Accessed

51. FDA. 2017. https://www.fda.gov/drugs/resources-information-approved-drugs/ribociclib-kisqali, *on* FDA. Accessed

52. FDA. 2021. https://www.fda.gov/drugs/resources-information-approved-drugs/fda-approves-abemaciclib-endocrine-therapy-early-breast-cancer, *on* FDA. Accessed

53. Kim S, Leong A, Kim M, Yang HW. 2022. CDK4/6 initiates Rb inactivation and CDK2 activity coordinates cell-cycle commitment and G1/S transition. Sci Rep 12:16810.

54. Narasimha AM, Kaulich M, Shapiro GS, Choi YJ, Sicinski P, Dowdy SF. 2014. Cyclin D activates the Rb tumor suppressor by mono-phosphorylation. Elife 3.

55. Yang HW, Cappell SD, Jaimovich A, Liu C, Chung M, Daigh LH, Pack LR, Fan Y, Regot S, Covert M, Meyer T. 2020. Stress-mediated exit to quiescence restricted by increasing persistence in CDK4/6 activation. eLife 9:e44571.

56. Malorni L, Piazza S, Ciani Y, Guarducci C, Bonechi M, Biagioni C, Hart CD, Verardo R, Di Leo A, Migliaccio I. 2016. A gene expression signature of retinoblastoma loss-of-function is a predictive biomarker of resistance to palbociclib in breast cancer cell lines and is prognostic in patients with ER positive early breast cancer. Oncotarget 7:68012–68022.

57. Palafox M, Monserrat L, Bellet M, Villacampa G, Gonzalez-Perez A, Oliveira M, Brasó-Maristany F, Ibrahimi N, Kannan S, Mina L. 2022. High p16 expression and heterozygous RB1 loss are biomarkers for CDK4/6 inhibitor resistance in ER+ breast cancer. nature communications 13:1–20.

58. Yang C, Li Z, Bhatt T, Dickler M, Giri D, Scaltriti M, Baselga J, Rosen N, Chandarlapaty S. 2017. Acquired CDK6 amplification promotes breast cancer resistance to CDK4/6 inhibitors and loss of ER signaling and dependence. Oncogene 36:2255–2264.

59. Li Z, Razavi P, Li Q, Toy W, Liu B, Ping C, Hsieh W, Sanchez-Vega F, Brown DN, Paula AFDC. 2018. Loss of the FAT1 tumor suppressor promotes resistance to CDK4/6 inhibitors via the hippo pathway. Cancer cell 34:893–905. e8.

60. Olanich ME, Sun W, Hewitt SM, Abdullaev Z, Pack SD, Barr FG. 2015. CDK4 Amplification Reduces Sensitivity to CDK4/6 Inhibition in Fusion-Positive Rhabdomyosarcoma. Clin Cancer Res 21:4947–59.

61. Takeshita T, Yamamoto Y, Yamamoto-Ibusuki M, Tomiguchi M, Sueta A, Murakami K, Iwase H. 2018. Clinical significance of plasma cell-free DNA mutations in PIK3CA, AKT1, and ESR1 gene according to treatment lines in ER-positive breast cancer. Mol Cancer 17:67.

62. Iida M, Toyosawa D, Nakamura M, Tsuboi K, Tokuda E, Niwa T, Ishida T, Hayashi S-i. 2020. Decreased ER dependency after acquired resistance to CDK4/6 inhibitors. Breast Cancer 27:963–972.

63. Costa C, Wang Y, Ly A, Hosono Y, Murchie E, Walmsley CS, Huynh T, Healy C, Peterson R, Yanase S. 2020. PTEN loss mediates clinical cross-resistance to CDK4/6 and PI3Kα inhibitors in breast cancer. Cancer discovery 10:72–85.

64. Kumarasamy V, Vail P, Nambiar R, Witkiewicz AK, Knudsen ES. 2021. Functional Determinants of Cell Cycle Plasticity and Sensitivity to CDK4/6 Inhibition. Cancer Research 81:1347–1360.

65. Patel P, Tsiperson V, Gottesman SRS, Somma J, Blain SW. 2018. Dual Inhibition of CDK4 and CDK2 via Targeting p27 Tyrosine Phosphorylation Induces a Potent and Durable Response in Breast Cancer Cells. Mol Cancer Res 16:361–377.

66. Iida M, Nakamura M, Tokuda E, Toyosawa D, Niwa T, Ohuchi N, Ishida T, Hayashi SI. 2019. The p21 levels have the potential to be a monitoring marker for ribociclib in breast cancer. Oncotarget 10:4907–4918.

67. Michaloglou C, Crafter C, Siersbaek R, Delpuech O, Curwen JO, Carnevalli LS, Staniszewska AD, Polanska UM, Cheraghchi-Bashi A, Lawson M, Chernukhin I, McEwen R, Carroll JS, Cosulich SC. 2018. Combined Inhibition of mTOR and CDK4/6 Is Required for Optimal Blockade of E2F Function and Long-term Growth Inhibition in Estrogen Receptor-positive Breast Cancer. Mol Cancer Ther 17:908–920.

68. Goel S, Wang Q, Watt AC, Tolaney SM, Dillon DA, Li W, Ramm S, Palmer AC, Yuzugullu H, Varadan V, Tuck D, Harris LN, Wong K-K, Liu XS, Sicinski P, Winer EP, Krop IE, Zhao JJ. 2016. Overcoming Therapeutic Resistance in HER2-Positive Breast Cancers with CDK4/6 Inhibitors. Cancer Cell 29:255–269.

69. O’Brien NA, McDermott MS, Conklin D, Luo T, Ayala R, Salgar S, Chau K, DiTomaso E, Babbar N, Su F. 2020. Targeting activated PI3K/mTOR signaling overcomes acquired resistance to CDK4/6-based therapies in preclinical models of hormone receptor-positive breast cancer. Breast Cancer Research 22:1–17.

70. Jansen VM, Bhola NE, Bauer JA, Formisano L, Lee KM, Hutchinson KE, Witkiewicz AK, Moore PD, Estrada MV, Sánchez V, Ericsson PG, Sanders ME, Pohlmann PR, Pishvaian MJ, Riddle DA, Dugger TC, Wei W, Knudsen ES, Arteaga CL. 2017. Kinome-Wide RNA Interference Screen Reveals a Role for PDK1 in Acquired Resistance to CDK4/6 Inhibition in ER-Positive Breast Cancer. Cancer Res 77:2488–2499.

71. Turner N, Pearson A, Sharpe R, Lambros M, Geyer F, Lopez-Garcia MA, Natrajan R, Marchio C, Iorns E, Mackay A, Gillett C, Grigoriadis A, Tutt A, Reis-Filho JS, Ashworth A. 2010. FGFR1 amplification drives endocrine therapy resistance and is a therapeutic target in breast cancer. Cancer Res 70:2085–94.

72. Formisano L, Lu Y, Servetto A, Hanker AB, Jansen VM, Bauer JA, Sudhan DR, Guerrero-Zotano AL, Croessmann S, Guo Y, Ericsson PG, Lee K-m, Nixon MJ, Schwarz LJ, Sanders ME, Dugger TC, Cruz MR, Behdad A, Cristofanilli M, Bardia A, O’Shaughnessy J, Nagy RJ, Lanman RB, Solovieff N, He W, Miller M, Su F, Shyr Y, Mayer IA, Balko JM, Arteaga CL. 2019. Aberrant FGFR signaling mediates resistance to CDK4/6 inhibitors in ER+ breast cancer. Nature Communications 10:1373.

73. Alves CL, Ehmsen S, Terp MG, Portman N, Tuttolomondo M, Gammelgaard OL, Hundebøl MF, Kaminska K, Johansen LE, Bak M, Honeth G, Bosch A, Lim E, Ditzel HJ. 2021. Co-targeting CDK4/6 and AKT with endocrine therapy prevents progression in CDK4/6 inhibitor and endocrine therapy-resistant breast cancer. Nature Communications 12:5112.

74. Dean JL, Thangavel C, McClendon AK, Reed CA, Knudsen ES. 2010. Therapeutic CDK4/6 inhibition in breast cancer: key mechanisms of response and failure. Oncogene 29:4018–4032.

75. Hayes TK, Luo F, Cohen O, Goodale AB, Lee Y, Pantel S, Bagul M, Piccioni F, Root DE, Garraway LA, Meyerson M, Johannessen CM. 2019. A Functional Landscape of Resistance to MEK1/2 and CDK4/6 Inhibition in NRAS-Mutant Melanoma. Cancer Research 79:2352–2366.

76. De Leeuw R, McNair C, Schiewer MJ, Neupane NP, Brand LJ, Augello MA, Li Z, Cheng LC, Yoshida A, Courtney SM. 2018. MAPK Reliance via Acquired CDK4/6 Inhibitor Resistance in CancerCDK4/6 Inhibitor Resistance in Cancer. Clinical Cancer Research 24:4201–4214.

77. Haines E, Chen T, Kommajosyula N, Chen Z, Herter-Sprie GS, Cornell L, Wong K-K, Shapiro GI. 2018. Palbociclib resistance confers dependence on an FGFR-MAP kinase-mTOR-driven pathway in KRAS-mutant non-small cell lung cancer. Oncotarget 9:31572.

78. Mo H, Liu X, Xue Y, Chen H, Guo S, Li Z, Wang S, Li C, Han J, Fu M, Song Y, Li D, Ma F. 2022. S6K1 amplification confers innate resistance to CDK4/6 inhibitors through activating c-Myc pathway in patients with estrogen receptor-positive breast cancer. Molecular Cancer 21:171.

79. Robinson AM, Rathore R, Redlich NJ, Adkins DR, VanArsdale T, Van Tine BA, Michel LS. 2019. Cisplatin exposure causes c-Myc-dependent resistance to CDK4/6 inhibition in HPV-negative head and neck squamous cell carcinoma. Cell death & disease 10:1–13.

80. Pandey K, Park N, Park K-S, Hur J, Cho YB, Kang M, An H-J, Kim S, Hwang S, Moon YW. 2020. Combined CDK2 and CDK4/6 inhibition overcomes palbociclib resistance in breast cancer by enhancing senescence. Cancers 12:3566.

81. Hall CR, Bisi JE, Strum JC. 2019. Inhibition of CDK2 overcomes primary and acquired resistance to CDK4/6 inhibitors. Cancer Research 79:4414–4414.

82. Jeselsohn R, Yelensky R, Buchwalter G, Frampton G, Meric-Bernstam F, Gonzalez-Angulo AM, Ferrer-Lozano J, Perez-Fidalgo JA, Cristofanilli M, Gomez H. 2014. Emergence of Constitutively Active Estrogen Receptor-α Mutations in Pretreated Advanced Estrogen Receptor–Positive Breast CancerESR1-Activating Mutations in Advanced Breast Cancer. Clinical cancer research 20:1757–1767.

83. Masri S, Phung S, Wang X, Wu X, Yuan Y-C, Wagman L, Chen S. 2008. Genome-wide analysis of aromatase inhibitor-resistant, tamoxifen-resistant, and long-term estrogen-deprived cells reveals a role for estrogen receptor. Cancer research 68:4910–4918.

84. Vesuna F, Lisok A, Kimble B, Domek J, Kato Y, van der Groep P, Artemov D, Kowalski J, Carraway H, van Diest P. 2012. Twist contributes to hormone resistance in breast cancer by downregulating estrogen receptor-α. Oncogene 31:3223–3234.

85. Johnston S, Saccani-Jotti G, Smith I, Salter J, Newby J, Coppen M, Ebbs S, Dowsett M. 1995. Changes in estrogen receptor, progesterone receptor, and pS2 expression in tamoxifen-resistant human breast cancer. Cancer research 55:3331–3338.

86. Miller TW, Hennessy BT, González-Angulo AM, Fox EM, Mills GB, Chen H, Higham C, García-Echeverría C, Shyr Y, Arteaga CL. 2010. Hyperactivation of phosphatidylinositol-3 kinase promotes escape from hormone dependence in estrogen receptor-positive human breast cancer. J Clin Invest 120:2406–13.

87. Campbell RA, Bhat-Nakshatri P, Patel NM, Constantinidou D, Ali S, Nakshatri H. 2001. Phosphatidylinositol 3-kinase/AKT-mediated activation of estrogen receptor α: a new model for anti-estrogen resistance. Journal of Biological Chemistry 276:9817–9824.

88. McNeil CM, Sergio CM, Anderson LR, Inman CK, Eggleton SA, Murphy NC, Millar EK, Crea P, Kench JG, Alles MC. 2006. c-Myc overexpression and endocrine resistance in breast cancer. The Journal of steroid biochemistry and molecular biology 102:147–155.

89. Planas-Silva MD, Bruggeman RD, Grenko RT, Smith J. 2007. Overexpression of c-Myc and Bcl-2 during progression and distant metastasis of hormone-treated breast cancer. Experimental and molecular pathology 82:85–90.

90. Abukhdeir AM, Vitolo MI, Argani P, De Marzo AM, Karakas B, Konishi H, Gustin JP, Lauring J, Garay JP, Pendleton C. 2008. Tamoxifen-stimulated growth of breast cancer due to p21 loss. Proceedings of the National Academy of Sciences 105:288–293.

91. Massarweh S, Osborne CK, Jiang S, Wakeling AE, Rimawi M, Mohsin SK, Hilsenbeck S, Schiff R. 2006. Mechanisms of tumor regression and resistance to estrogen deprivation and fulvestrant in a model of estrogen receptor–positive, HER-2/neu-positive breast cancer. Cancer research 66:8266–8273.

92. Zhou FH, Downton T, Freelander A, Hurwitz J, Caldon CE, Lim E. 2023. CDK4/6 inhibitor resistance in estrogen receptor positive breast cancer, a 2023 perspective. Frontiers in Cell and Developmental Biology Volume 11–2023.

93. Chandarlapaty S. 2012. Negative Feedback and Adaptive Resistance to the Targeted Therapy of Cancer. Cancer Discovery 2:311–319.

94. Movasat H, Giacopino E, Shahdoost A, Dorri Nokoorani Y, Abrbekouh AH, Tahamtani Y, Shakiba N. 2025. A systems view of cellular heterogeneity: Unlocking the "wheel of fate". Cell Syst 16:101300.

95. Babtie AC, Stumpf MPH. 2017. How to deal with parameters for whole-cell modelling. Journal of The Royal Society Interface 14.

96. Yamada T, Nishiyama M, Oba S, Jimbo HC, Ikeda K, Ishii S, Hong K, Sakumura Y. 2018. Computational Methods for Estimating Molecular System from Membrane Potential Recordings in Nerve Growth Cone. Scientific Reports 8:4559.

97. Spencer SL, Gaudet S, Albeck JG, Burke JM, Sorger PK. 2009. Non-genetic origins of cell-to-cell variability in TRAIL-induced apoptosis. Nature 459:428–432.

98. Ghandi M, Huang FW, Jané-Valbuena J, Kryukov GV, Lo CC, McDonald III ER, Barretina J, Gelfand ET, Bielski CM, Li H. 2019. Next-generation characterization of the cancer cell line encyclopedia. Nature 569:503–508.

99. Migliaccio I, Bonechi M, Romagnoli D, Boccalini G, Galardi F, Guarducci C, Nardone A, Schiff R, Biganzoli L, Malorni L, Benelli M. 2025. Single-cell transcriptomics reveals biomarker heterogeneity linked to CDK4/6 Inhibitor resistance in breast cancer cell lines. npj Breast Cancer 11:82.

100. Asghar US, Barr AR, Cutts R, Beaney M, Babina I, Sampath D, Giltnane J, Lacap JA, Crocker L, Young A, Pearson A, Herrera-Abreu MT, Bakal C, Turner NC. 2017. Single-Cell Dynamics Determines Response to CDK4/6 Inhibition in Triple-Negative Breast Cancer. Clin Cancer Res 23:5561–5572.

101. Taniguchi Y, Choi PJ, Li GW, Chen H, Babu M, Hearn J, Emili A, Xie XS. 2010. Quantifying E. coli proteome and transcriptome with single-molecule sensitivity in single cells. Science 329:533–8.

102. Bar-Even A, Milo R, Noor E, Tawfik DS. 2015. The Moderately Efficient Enzyme: Futile Encounters and Enzyme Floppiness. Biochemistry 54:4969–77.

103. Bengtsson M, Ståhlberg A, Rorsman P, Kubista M. 2005. Gene expression profiling in single cells from the pancreatic islets of Langerhans reveals lognormal distribution of mRNA levels. Genome Res 15:1388–92.

104. Kwon T, Kwon O-S, Cha H-J, Sung BJ. 2019. Stochastic and Heterogeneous Cancer Cell Migration: Experiment and Theory. Scientific Reports 9:16297.

105. Boyeau P, Hong J, Gayoso A, Kim M, McFaline-Figueroa JL, Jordan MI, Azizi E, Ergen C, Yosef N. 2025. Deep generative modeling of sample-level heterogeneity in single-cell genomics. Nature Methods 22:2264–2274.

106. Yip HYK, Shin SY, Chee A, Ang CS, Rossello FJ, Wong LH, Nguyen LK, Papa A. 2024. Integrative modeling uncovers p21-driven drug resistance and prioritizes therapies for PIK3CA-mutant breast cancer. NPJ Precis Oncol 8:20.

