## Supplementary Information for "Systematic Analysis of Network-driven Adaptive Resistance to CDK4/6 and Estrogen Receptor Inhibition using Meta-Dynamic Network Modelling"

^1^Department of Biochemistry and Molecular Biology, Faculty of Medicine, Nursing and Health Sciences, Monash University, Clayton, Victoria 3800, Australia. ^2^Biomedicine Discovery Institute, Monash University, Clayton, Victoria 3800, Australia. ^3^Computational Systems Oncology Program, South Australian immunoGENomics Cancer Institute (SAiGENCI), The University of Adelaide, Adelaide, SA 5005, Australia. ^4^Australian Research Council Centre of Excellence for the Mathematical Analysis of Cellular Systems (MACSYS), Australia.

### **Description of Model Scope and Construction**

To keep the model to a tractable size we selected only two major mitogenic signalling pathways, PI3K-AKT and MAPK, frequently mutated in cancer. For this model we chose the insulin receptor (IR) and the fibroblast growth factor receptor (FGFR) to be representative of the many receptor tyrosine kinases (RTKs) capable of stimulating these pathways. The PI3K-AKT signalling pathway feeds into the cell cycle machinery through two parallel mechanisms. First is through the inactivation of GSK3B and the second is through the promotion of cyclin D transcription by active mTORC1 (*1, 2*). Unphosphorylated GSK3B promotes the degradation of cyclin E, A, and the transcription factor Myc, and its phosphorylation by active AKT deactivates this ability and promotes cell cycle progression (*3, 4*). The transcription of cyclins enables them to bind to and activate their cognate CDKs and similarly promote cell cycle progression. Contained within the PI3K-AKT motif is a negative feedback loop wherein activated ribosomal protein S6 kinase beta (S6K) inhibits insulin receptor mediated signalling (*5*).

The promotion of cell cycle progression by the MAPK cascade has been included in this model in three ways. The first is the direct promotion of cyclin D transcription by the activation of the AP1 transcription factor by ERK (*6*). The second is through the phosphorylation of Myc, which stabilises it and prevents its degradation (*7*). Both phosphorylated and unphosphorylated forms of Myc promote cell cycle progression in many ways. Myc promotes cyclin transcription, promotes the transcription of E2F and enhances E2F’s transcriptional activity and enhances the ability of AKT to phosphorylate p21 and p27 (*8-13*). The third way ERK promotes cell cycle progression is promotion of estrogen receptor (ER) activity (*14*). In particular, ERK enhances the ability of the ER to initiate cyclin D transcription (*15*). There is a negative feedback loop within the MAPK cascade where the activation of ERK causes it to inhibit upstream Ras signalling (*16*). There is also an important crosstalk mechanism that exists between the PI3K-AKT and MAPK signalling pathways; the inhibition of Raf activation by active AKT (*17*).

E2F is thought to be one of the most important transcription factors in pushing the cell cycle towards S-phase as it drives transcription of cyclins E and A (*18, 19*). E2F is inhibited by unphosphorylated Rb, but this inhibition is partially relieved by the monophosphorylation of Rb by the active CDK4/6-cyclin D complex (*20*). Canonically, the partial release of inhibition enables the transcription of cyclin E, which in turn activates CDK2. Active CDK2-cyclin E complexes then hyperphosphorylate Rb, fully relieving the inhibition of E2F and enabling the transcription of cyclin A (*20*). The transcription of cyclin A and its binding to CDK2 represent a point at which the cell has well and truly entered the S-phase of the cell cycle. CDK2-cyclin A complexes have been shown to stabilise the double phosphorylated form of AKT whereas hyperphosphorylated Rb has been shown to be capable of translocating into the cytoplasm and inhibiting the activation of mTORC2 (*21, 22*). In our model p21 and p27 are included as a single species where the phosphorylated form promotes CDK4/6-cyclin D complex formation and the unphosphorylated form inhibits the complex formation of CDK2-cyclin E (*23, 24*).

Due to its importance in ER+BC, ER was also included in our model. Its main function is the promotion of cyclin D transcription, but it is also capable of promoting Myc transcription (*25, 26*). FOXO3 was included due to its ability to promote ER activity by upregulating its transcription (*27*). FOXO3 is regulated by AKT whereby AKT phosphorylates FOXO3, causing it to become localised in the cytosol and unable to regulate transcription (*28*). The network schematic generated from these observations can be seen in **Figure 2**. It should be noted that some protein species from with the mitogenic and cell cycle pathways were excluded/abstracted to keep the overall model to a size that could be effectively explored using the available computational resources.

The following is a brief explanation/justification of each reaction within the model. Any departure from the biological cause-and-effect relationships between proteins was done to reduce model complexity and thus enable greater and more in-depth computational analysis. In general, equations utilise first-order kinetics. This choice was made to reduce the number of parameters for the same reasons we attempted to reduce model complexity, to reduce the model’s computational burden. While utilising more sophisticated rate laws, such as Michaelis-Menten kinetics, may improve the biological accuracy of the model description, we believe the loss is minimal, and worth the gain in our ability to perform more sophisticated analyses. In the future we would like to explore smaller models but utilise more biologically accurate rate laws to precisely determine the effect of rate law choice on model dynamics.

**R1: IR ⬄ pIR**

This reaction describes the dimerization and autophosphorylation of the insulin receptor, stimulated by insulin.

**R2: PI3K ⬄ pPI3K**

This reaction captures the binding of IRS to IR, recruitment of the PI3K regulatory unit and the recruitment of the PI3K catalytic subunit. As IRS is inhibited by active S6K, this component of the reaction is inhibited by S6K. PI3K is also able to be activated by the FGF receptor, though GRB2.

**R3: PIP2 ⬄ PIP3**

PIP2 is phosphorylated by active PI3K to become PIP3.

**R4: PDK1 <=> aPDK1**

PDK1 is recruited to PIP3, bring it into proximity to its targets. This is modelled as an active form of PDK1. As PIP3 is usually far in excess of PDK1, this reaction is modelled as PIP3 catalysing PDK1.

**R5: mTORC2 <=> amTORC2**

mTORC2 is also recruited to PIP3, brining it in proximity to its targets. This is modelled as an active form of mTORC2. As PIP3 is usually far in excess of mTORC2, this reaction is modelled as PIP3 catalysing mTORC2.

**R6: AKT <=> pAKT308**

Akt is phosphorylated at T308 by PDK1.

**R7: pAKT308 <=> ppAKT308473**

Akt is phosphorylated at S473 by mTORC2. Akt phosphorylation has been modelled as a linear progression to reduce complexity. Akt is fully active when it is dual phosphorylated, but it is still active with only a single phosphorylation. This has been incorporated into the model using an AKTscale parameter, which scales the activity of the dual phosphorylated Akt. CDK2-cyclinA has also been demonstrated to phosphorylate the tail of Akt, promoting the dual phosphorylation of Akt, thus CDK2cycA has been modelled as promoting this reaction.

**R8: mTORC1 <=> amTORC1**

Akt inhibits TSC2, which in turn inhibits mTORC1. TSC2 has been excluded from this model, and as such Akt activates mTORC1.

**R9: S6K <=> pS6K**

S6K is phosphorylated and activated by mTORC1.

**R10: FGFR <=> pFGFR**

Similar to IR, this reaction represents the activation of the FGF receptor by the binding of its ligand FGF. E2F is capable of stimulating the expression of FGFR, but as FGFR is not able to be synthesised in this model, this process has been modelled as E2F promoting FGFR activity.

**R11: Ras <=> aRas**

Ras is activated through its recruitment to the membrane, which is achieved by the recruitment of GRB2 to both IR and FGFR. GRB2 has not been included in this model and so the activation of Ras has been modelled as being catalysed by the active form of both receptors. SOS is also involved in the recruitment of Ras and is inhibited by ERK. Thus, as SOS is also not in this model, this reaction has been modelled as being inhibited by ERK.

**R12: Raf <=> aRaf**

Raf is recruited to the membrane by Ras. Raf is also phosphorylated and inhibited by Akt.

**R13: MEK <=> pMEK**

Raf phosphorylates and activates MEK.

**R14: ERK <=> pERK**

MEK phosphorylates and activates ERK.

**R15: ERa <=> aERa**

Like IR and FGFR, the estrogen receptor is activated by the binding of its cognate ligand, estrogen. ER synthesis can also be promoted by FOXO3, and this relationship has been modelled in a similar fashion to the promotion of FGFR activation by E2F. This reaction is also inhibited by the ER drug, which has been modelled as inhibiting the activation of ER by estrogen; the inhibition was modelled by reducing the value of the activation parameter by a fixed fraction.

**R16: => Myc**

Myc synthesis can be promoted by the ER. This promotion can be promoted by ERK, cyclin D and E2F.

**R17: Myc =>**

GSK3B promotes Myc degradation.

**R18: Myc <=> pMyc**

ERK phosphorylates and stabilises Myc.

**R19: GSK3b <=> pGSK3b**

Akt phosphorylates and inhibits GSK3B.

**R20: FOXO3 <=> pFOXO3**

Foxo3 is phosphorylated by Akt, which excludes it from the nucleus and inhibits its transcriptional activity. Thus, pFOXO3 is an inactive form of FOXO3.

**R21: => cycD**

Cyclin D synthesis is independently regulated by mTORC1, ERK, Myc and ER. The mechanisms all vary, where mTORC1 promotes general protein synthesis, ERK and Myc have been shown to promote transcription factors that in turn regulate cyclin D synthesis, and ER can directly regulate cyclin D synthesis. ERK and cyclin D can also promote the regulation of cyclin D synthesis by ER.

**R22: cycD =>**

Cyclins are generally unstable proteins with short half-lives, but this process can be dramatically sped up by GSK3B phosphorylation targeting them for ubiquitin mediated degradation.

**R23: => cycE**

Cyclin E synthesis is directly regulated by E2F and Myc. Myc is also known to promote the activity of E2F.

**R24: cycE =>**

Similar to cyclin D, cyclin E degradation is regulated by GSK3B.

**R25: => cycA**

Similar to cyclin E, cyclin A has been shown to be regulated by E2F and Myc.

**R26: cycA =>**

Cyclin A degradation is regulated by proteins not included in this model.

**R27: CDK46 + cycD <=> CDK46cycD**

The formation of the CDK4/6-cyclin D complex has been shown to be promoted by p21 and p27. The CDK4/6 drug has been modelled to inhibit the formation of this complex; the inhibition was modelled by reducing the value of the association parameter by a fixed fraction.

**R28: CDK2 + cycE <=> CDK2cycE**

Canonically, p21 and p27 are CDK2 inhibitors, and so this relationship has been modelled as p21p27 inhibiting the formation of this complex.

**R29: CDK2 + cycA <=> CDK2cycA**

There is evidence to suggest that Akt phosphorylation can promote the formation of the CDK2-cyclin A complex.

**R30: p2127 <=> pp2127**

Akt can phosphorylate and stabilise p21.

**R31: => E2F**

Myc can promote the synthesis of E2F.

**R32: E2F =>**

The regulation of E2F degradation is performed by proteins not included in this model.

**R33: E2F + Rb <=> RbE2F**

Unphosphorylated Rb binds to E2F inhibiting its ability to regulate transcription.

**R34: Rb <=> pRb**

The CDK4/6-cyclin D complex monophosphorylated E2F, partially relieving the inhibition of Rb on E2F.

**R35: pRb <=> pppRb**

CDK2-cyclin E hyperphosphorylates Rb, leading to its full dissociation from E2F.

### **Error Convergence**

Early experimental observations seemed to suggest that the distribution of protein dynamics was not changing dramatically when we increased the number of model instances being tested. This seemed to suggest that despite the massive number of possible model instances that could be generated given the parameter and initial condition hyperspaces we were searching, the distribution of protein dynamics was fixed well before this number was even close to being reached. To confirm this hypothesis, we calculated the difference between increasingly large populations of model instances to identify if/when the distribution of protein dynamics was converging to a fixed distribution. To measure the convergence, we first recorded the frequency of each dynamic for each protein was measured for N and 2N-sized populations of model instances/parameter sets. We then calculated the squared difference, or error, between the frequencies for each protein species, summed all of the error values and then divided by the number of observations (6 x 50: dynamic categories x number of state variables). In this way we were able to calculate the mean squared error (MSE) as the population size increased and measure the rate of convergence for this particular network topology. See supplementary **Figure S12**.

In addition to the MSE analysis described above, we also repeated the model instance generation and analysis in triplicate. By this we mean we generated 3 groups of 100,000 unique model instances for a total of 300,000 unique model instances. Each group was analysed separately, producing 3 meta-dynamic maps. These distributions of dynamics were then compared. We calculated that the average standard deviation between each distribution of each protein and found it to be 0.000329 or 0.0329% i.e. there is only a 0.0329% difference in the quantity of each protein dynamic on average between replicates. From our MSE analysis and the incredibly high similarity of distributions between completely independent replicates we conclude that the distribution of dynamics has well and truly converged by 100,000 parameter sets.

### **Supplementary Figures**

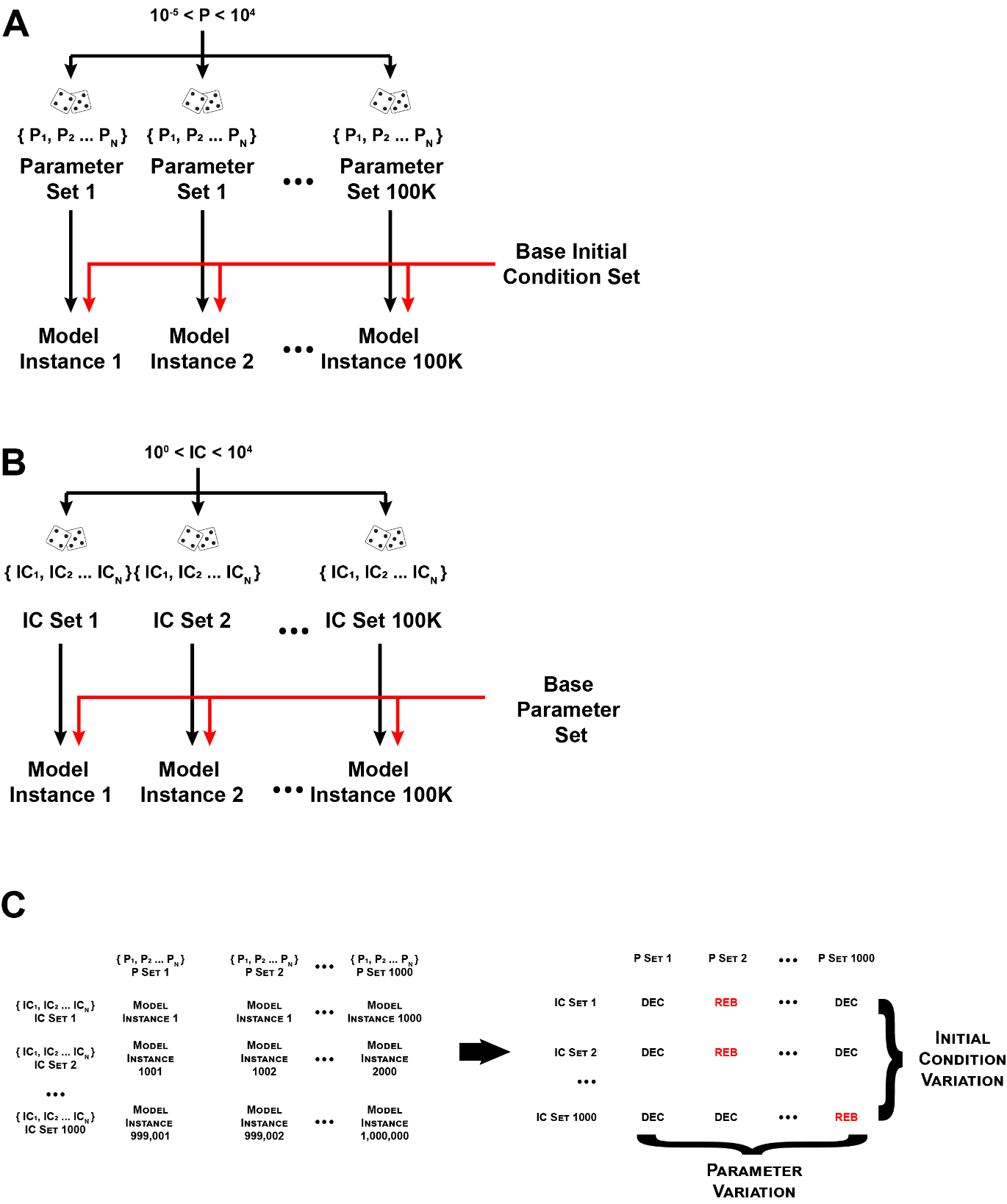

**Supplementary Figure S1:** Methods for generating groups of 100,000 unique model instances. A) Model instances are produced by combining a base set of initial conditions with a unique set of parameters. Parameter values in each unique parameter set are generated by randomly selecting values for parameters from between the range 10^-5^ and 10^4^. B) Model instances are produced by combining a base parameter set with a unique set of initial conditions. Initial condition values in each unique initial condition set are produced by randomly selecting values for initial conditions between the range 10^0^ and 10^4^. C) Measuring the ability of parameter variation versus initial condition variation to induce resistance. First, we selected 1000 model instances from the group of model instances produced in A) that demonstrated decreasing dynamics for one of the key output proteins, and 1000 model instances produced in B) that show the same. We then generated 1,000 new, unique model instances *per* model instance in group A by taking the parameter set of each group A model instance as the base parameter set and combing it with all of the initial condition sets from group B, see left side of panel. This produced a total of 1,000,000 unique model instances. We then simulated each of these model new model instances and determined the protein dynamic of each of the key output proteins. We then calculated the distribution of observed resistance-associated dynamics for the rows and columns. The distribution of the rows is representative of initial condition variation and the distribution of the columns is representative of parameter variation.

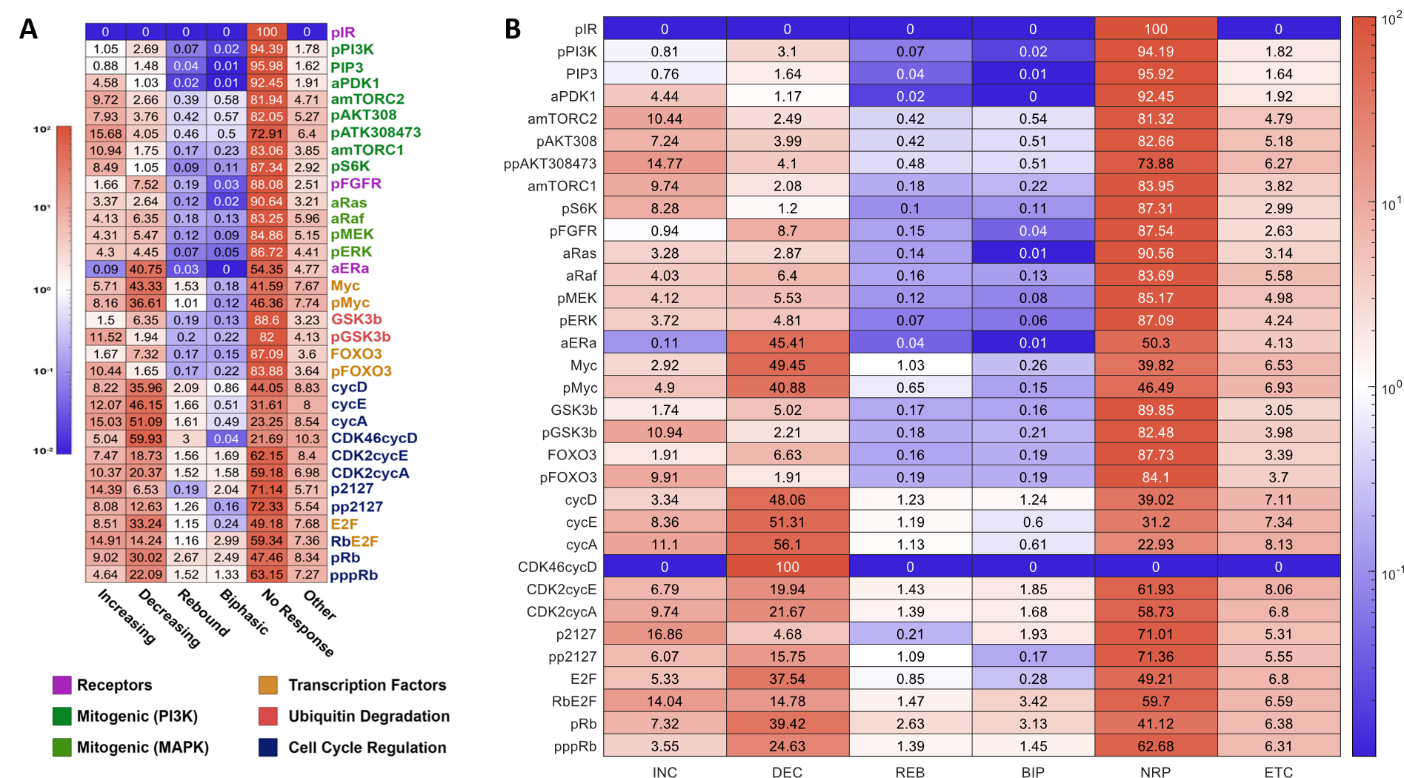

**Supplementary Figure S2.** **A)** Heatmap showing the frequency of each dynamic for a selection of active protein species, across 100,000 model instances (parametric variation) i.e. the network’s meta-dynamic map. Note that CDK4/6cycD shows a sustained decreasing behaviour in only 59.93% of model instances. **B)** Heatmap of dynamic frequencies, filtered for model instances that display strong suppression of CDK46cycD. Filtering in this way has very little effect on the distribution of protein dynamics.

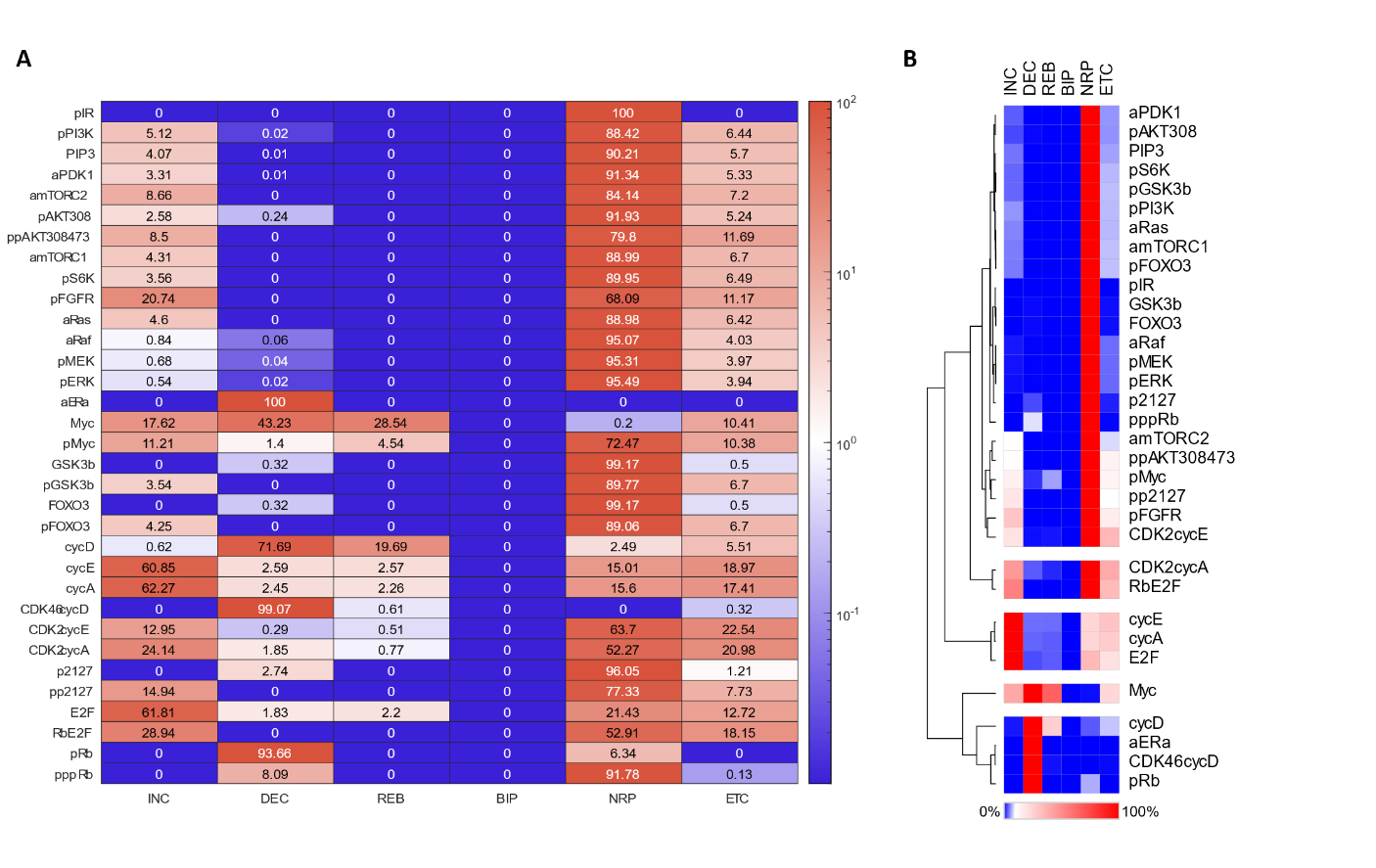

**Supplementary Figure S3**. **A)** Heatmap showing the frequency of each dynamic for a selection of active protein species, across 100,000 model instances when are initial condition values are varied. The base parameter set used was initially randomly selected. Blue represents low frequency and red represents high frequency. **B)** Clustering of the heatmap in A to highlight proteins with similar distributions of dynamic categories.

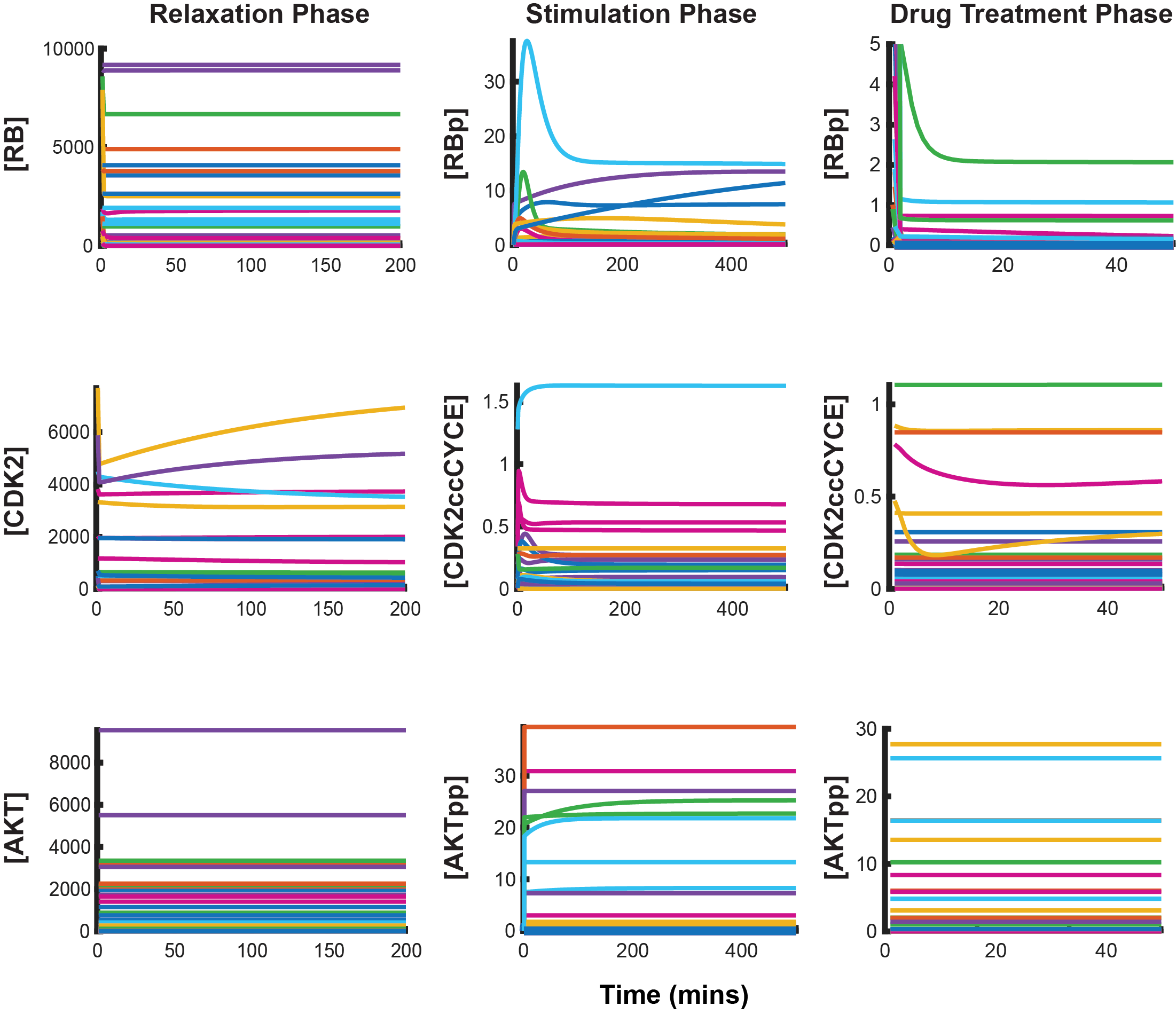

**Supplementary Figure S4**. Varying conserved totals shifts the dynamic equilibria of the relevant protein species. Plots of 50 different model instances, each with an identical base parameter set but with different conserved totals for its constituent protein species. The first column demonstrates the shifted equilibria with no stimulation, the second under (constant) mitogenic stimulation and the third after drug treatment. Variety can be seen in both steady state equilibria and in the qualitative dynamics before stead state is reached.

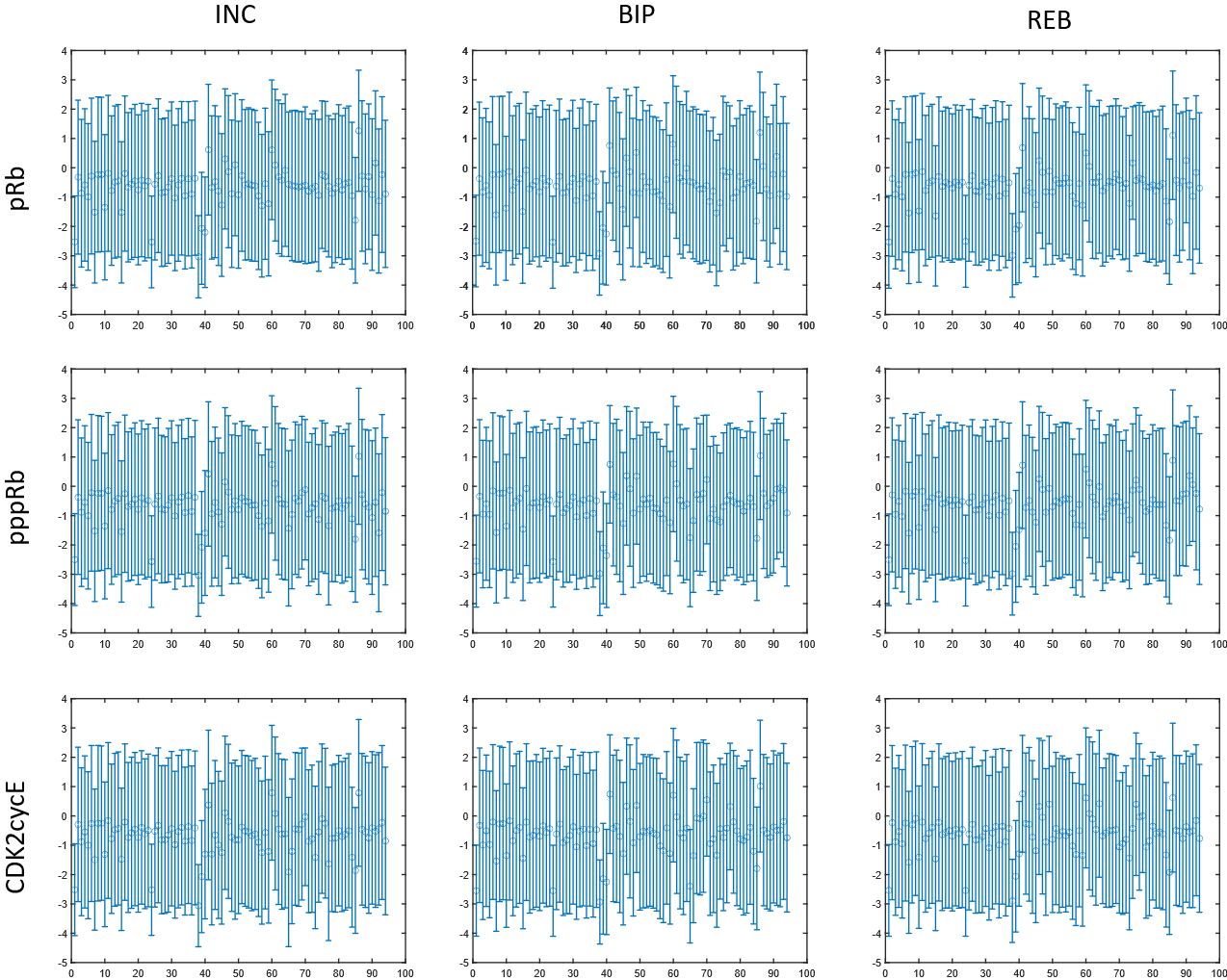

**Supplementary Figure S5.** Mean and standard deviation of the absolute log10 transformed parameter values for each protein-dynamic combination. Y-axis represents the range of possible log10 parameter values, the X-axis represents each individual parameter (in the order they appear in the IQM reaction file).

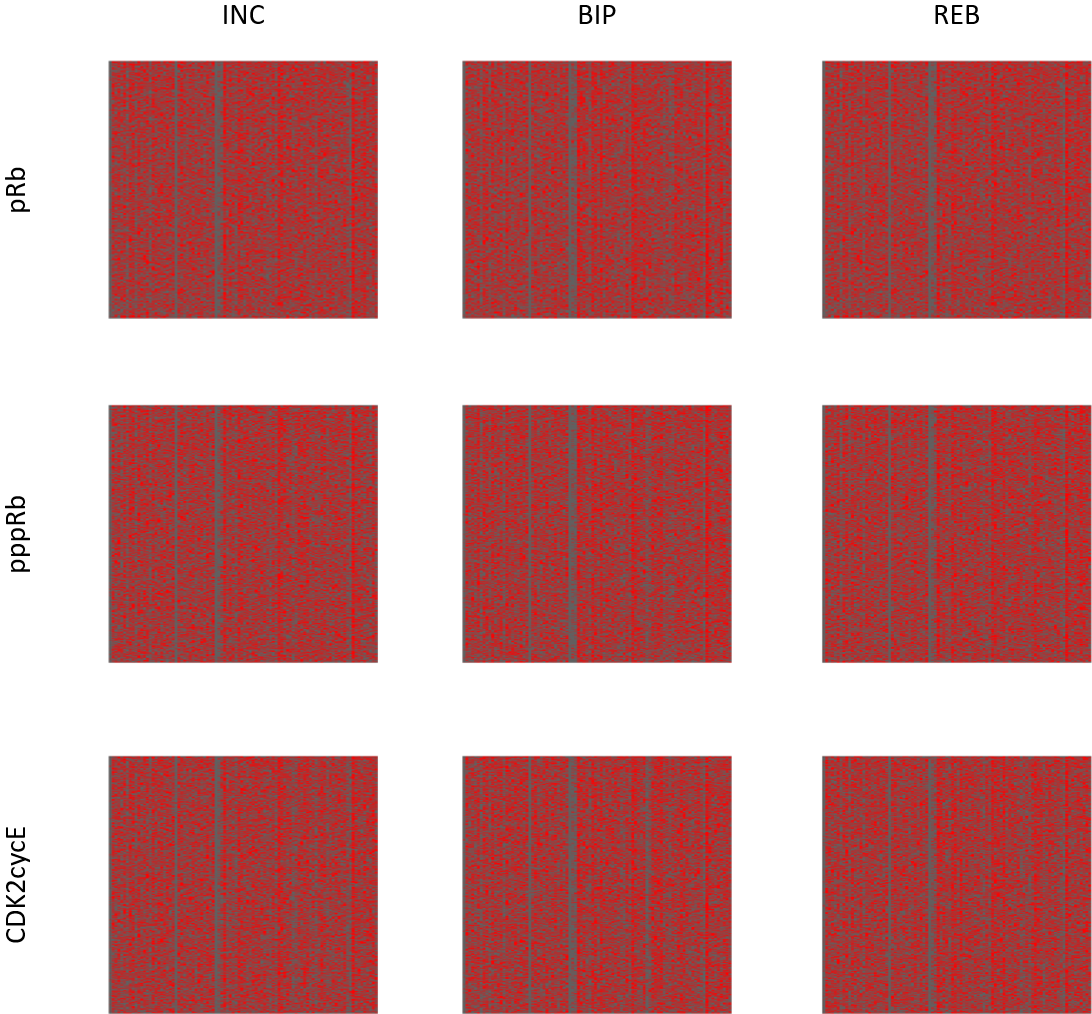

**Supplementary Figure S6.** A) Heatmaps showing the results of the hierarchical clustering of the raw parameter values for each protein-dynamic combination. Very little clustering can be observed, suggesting that there are no patterns in the absolute values of parameters with respect to resistance associated dynamics.

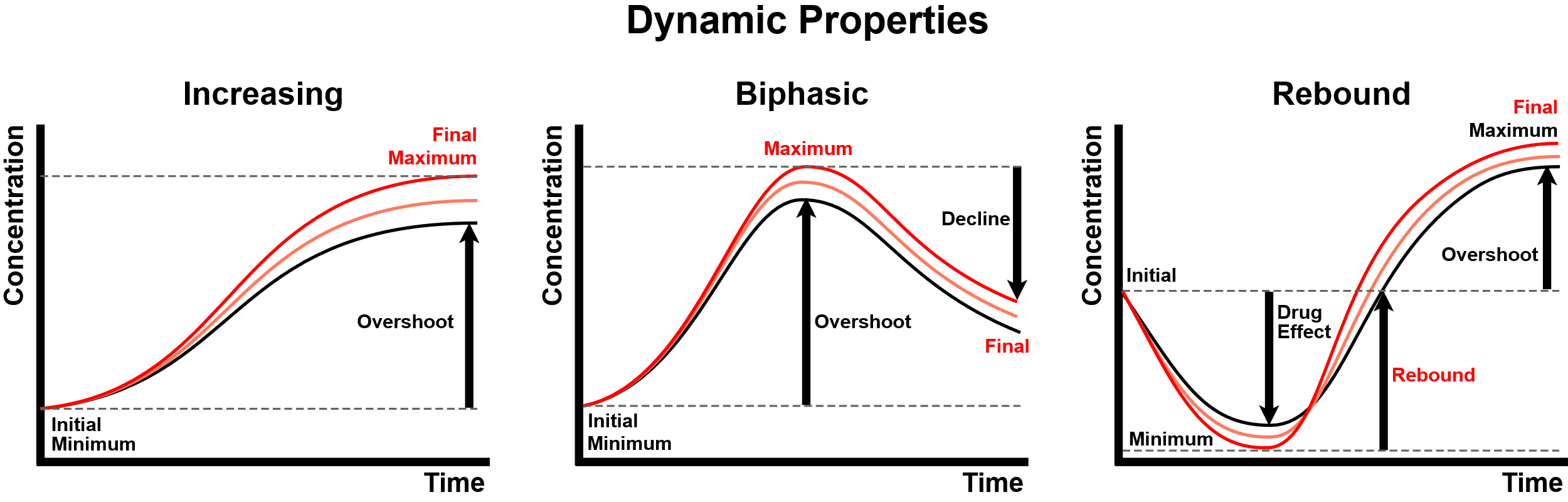

**Figure S7.** Breakdown of the various measurable features of a protein’s dynamic response following drug perturbation. Resistance features (red) are a subset of these dynamic features which are defined by a recovery or increase in the activity of a protein following drug perturbation, represented by the increase in concentration of the active form of a protein. The transition from black to red lines represent an increase in the resistance being displayed.

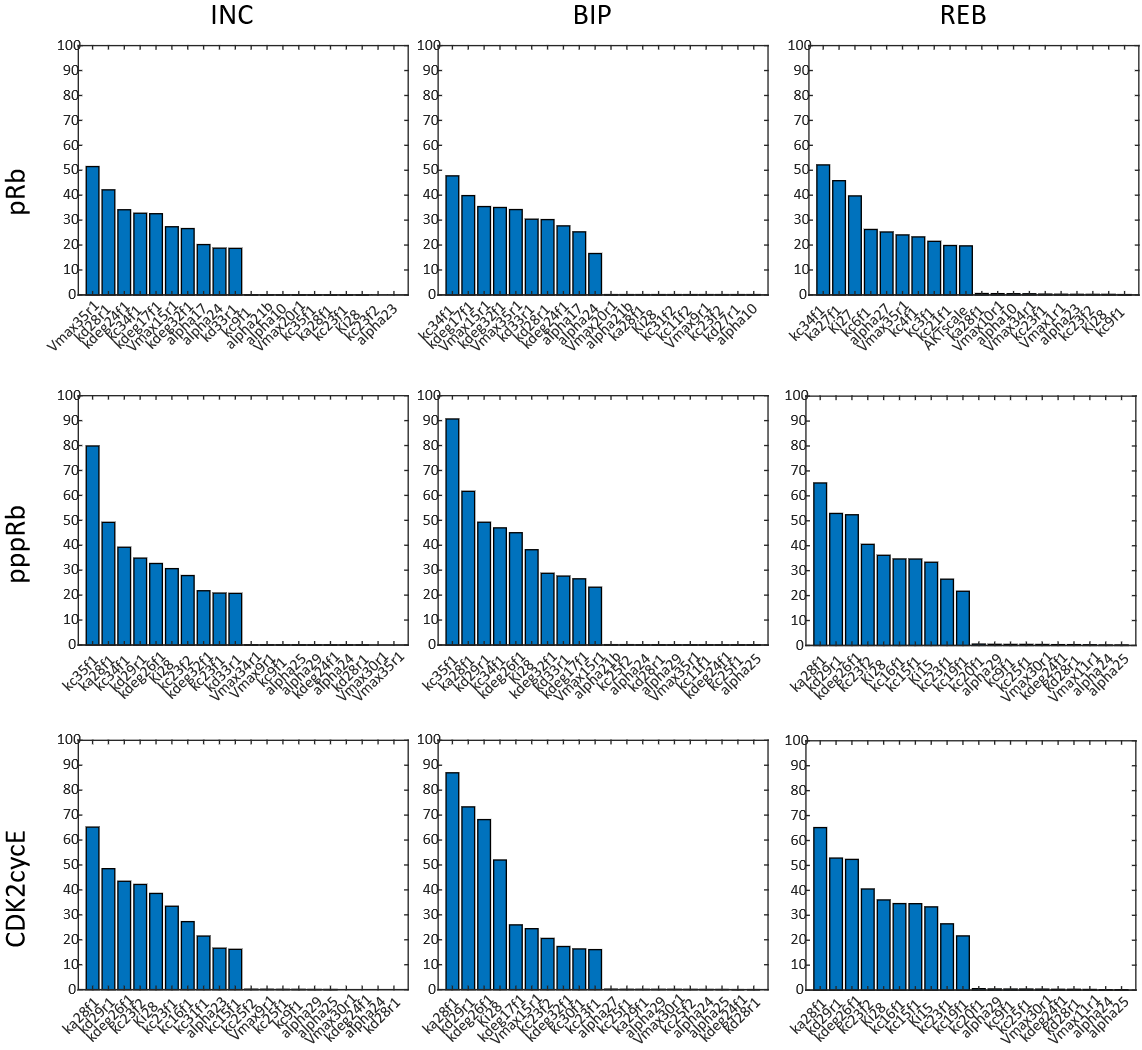

**Figure S8**. Top and bottom 10 ranked parameters for each protein-dynamic combination. Ranking is based on the overall contribution of each parameter to the resistance features for dynamic of each protein-dynamic combination. For biphasic and increasing dynamics, the ability of each parameter to contribute to the maximum concentration and final concentration was measured, and for the rebounding dynamics, the contribution to rebound and final concentration was measured. The y-axis refers to the percentage of the model instances in which the parameters along the x-axis contribute to the relevant resistance features, for that particular protein dynamic-output protein category.

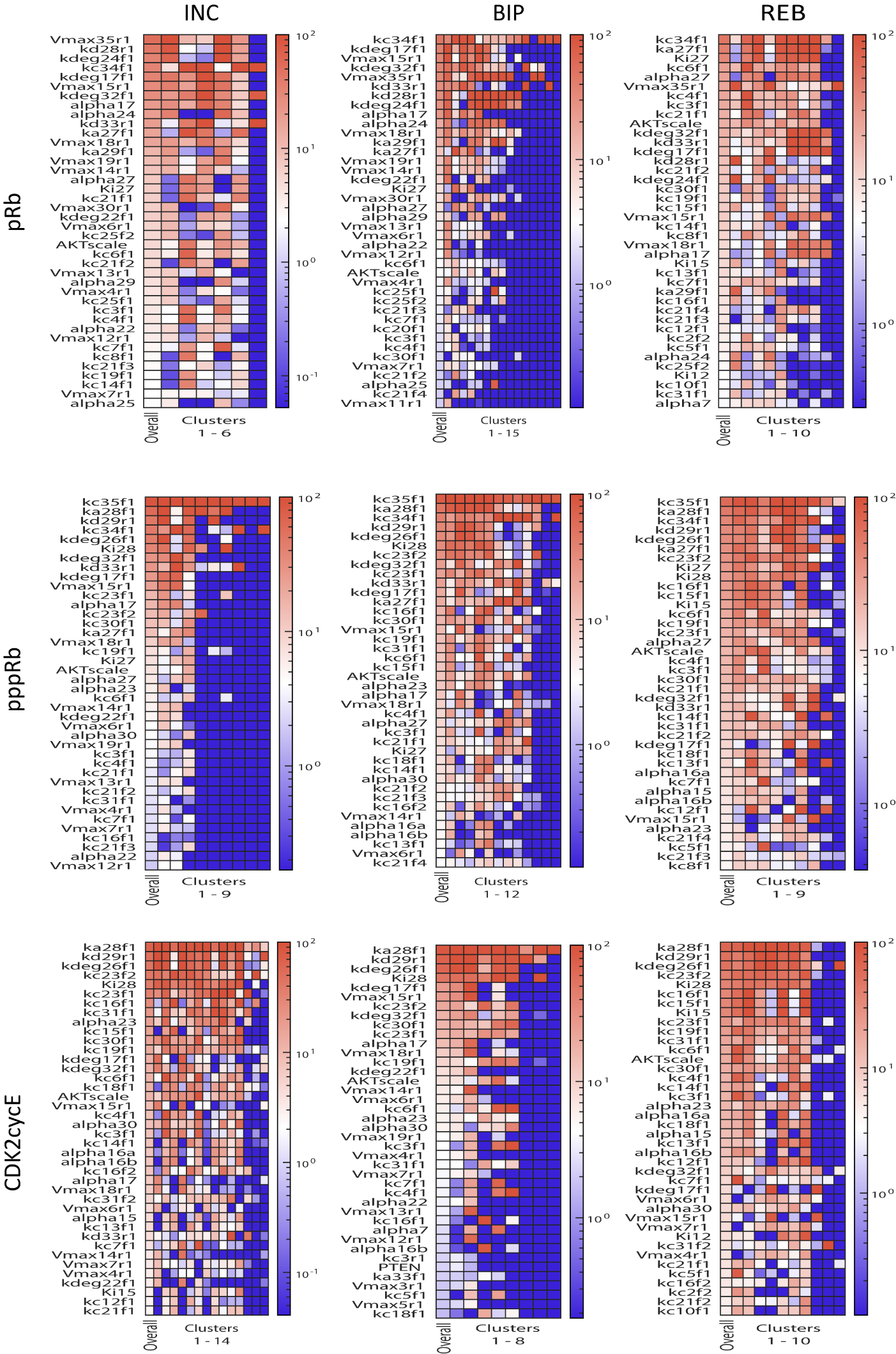

**Figure S9.** Heatmaps comparing the overall ranking of each protein-dynamic combination to the cluster-specific rankings. Rank is determined by how frequently each parameter contributes to the resistance features of the dynamic of each protein-dynamic combination.

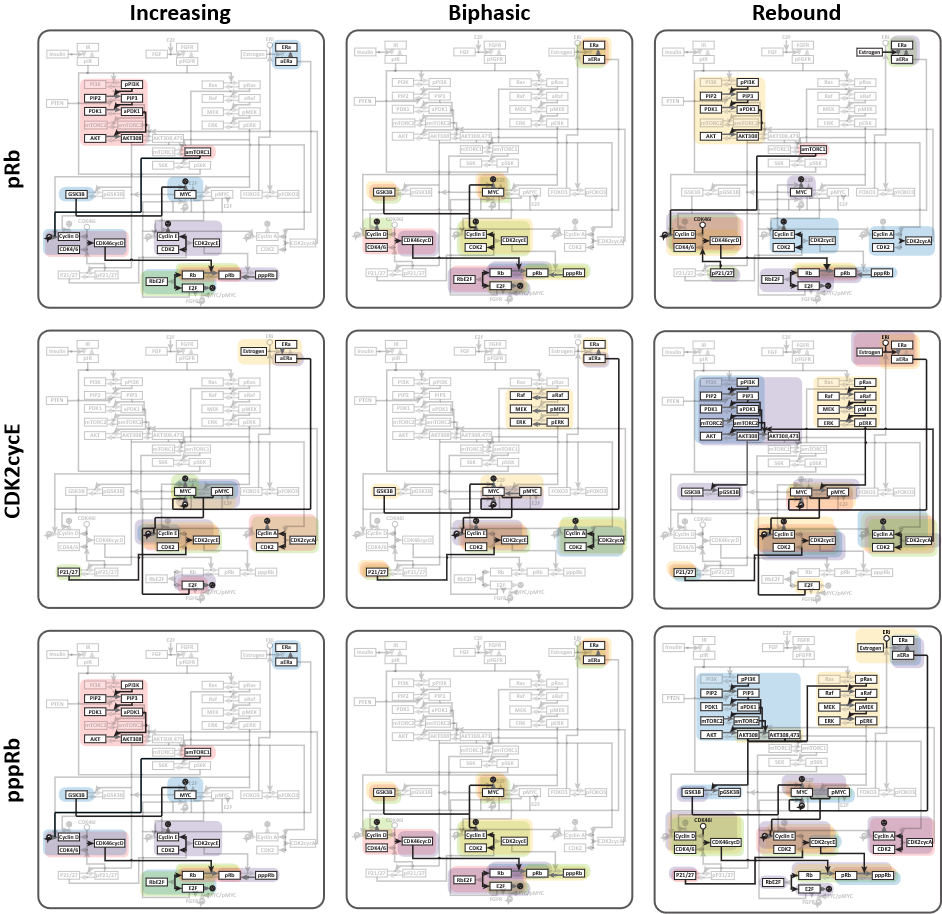

**Figure S10.** Parameter-signature ECC network overlays for each of the 9 protein-dynamic combinations, presented side-by-side for easier comparison of protein-dynamic combinations. Note that Figures S9A-I display the results for the individual protein-dynamic combination in more detail.

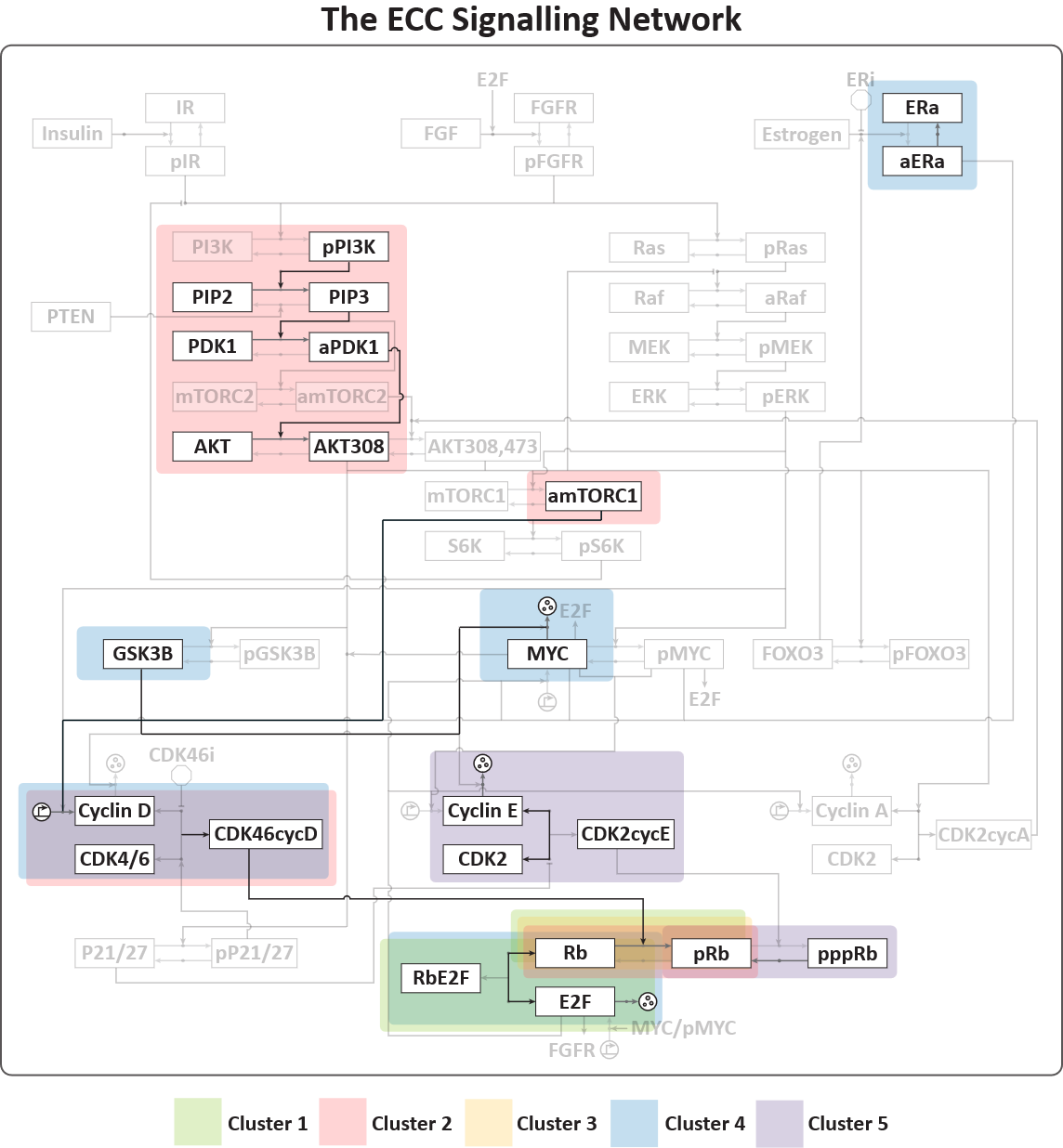

**Figure S10A. Cluster-specific parameter signatures with the ECC network, here displayed for the protein-dynamic pRb-INC combination.** Each figure (Figures S9A-I) is the overlay of the respective cluster-specific parameter signatures with the ECC network, for each protein-dynamic combination. Each figure displays the five largest clusters for each protein-dynamic combination.

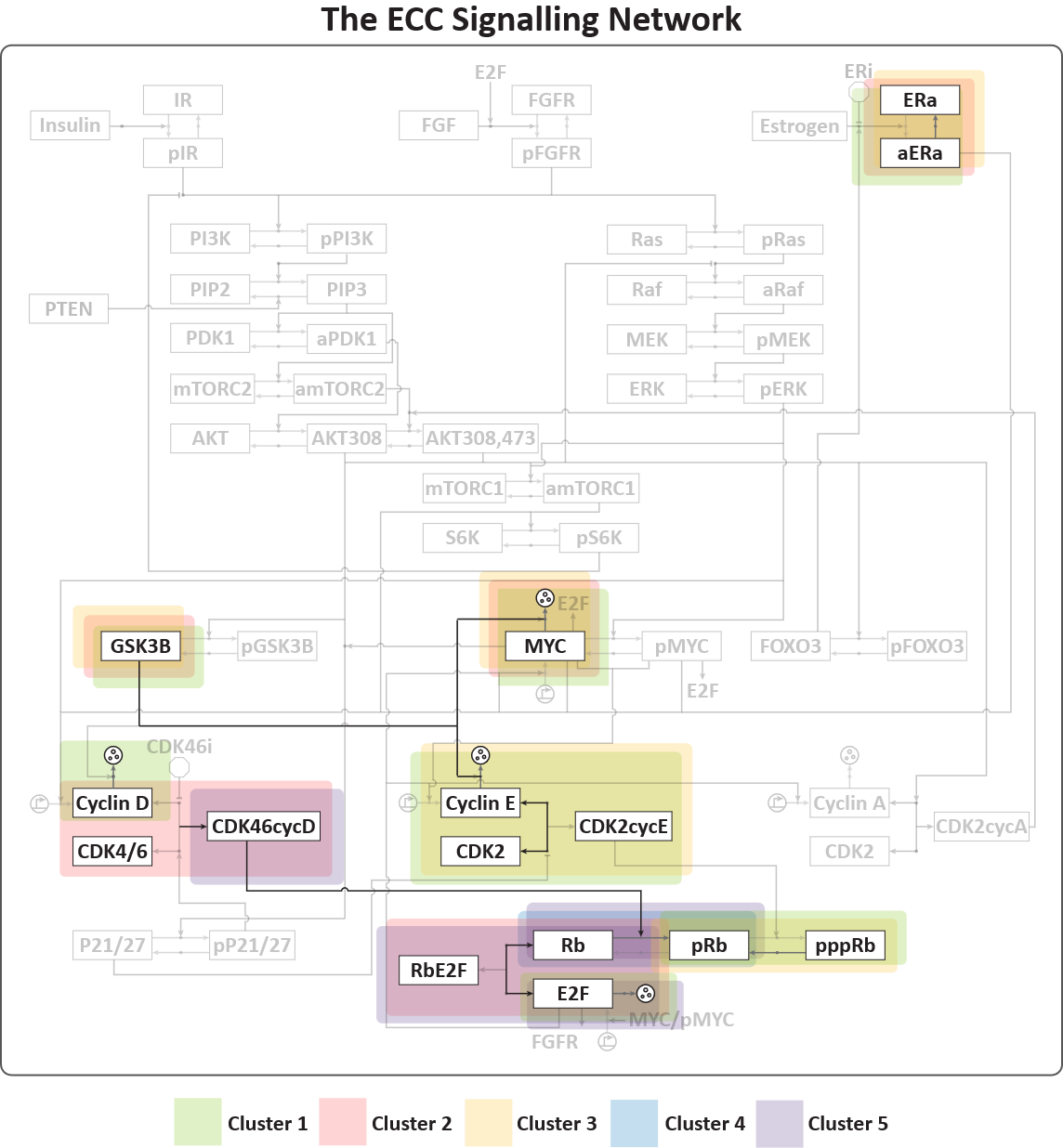

**Figure S10B. Cluster-specific parameter signatures with the ECC network, here displayed for the protein-dynamic pRb-BIP combination.** Each figure (Figures S9A-I) is the overlay of the respective cluster-specific parameter signatures with the ECC network, for each protein-dynamic combination. Each figure displays the five largest clusters for each protein-dynamic combination.

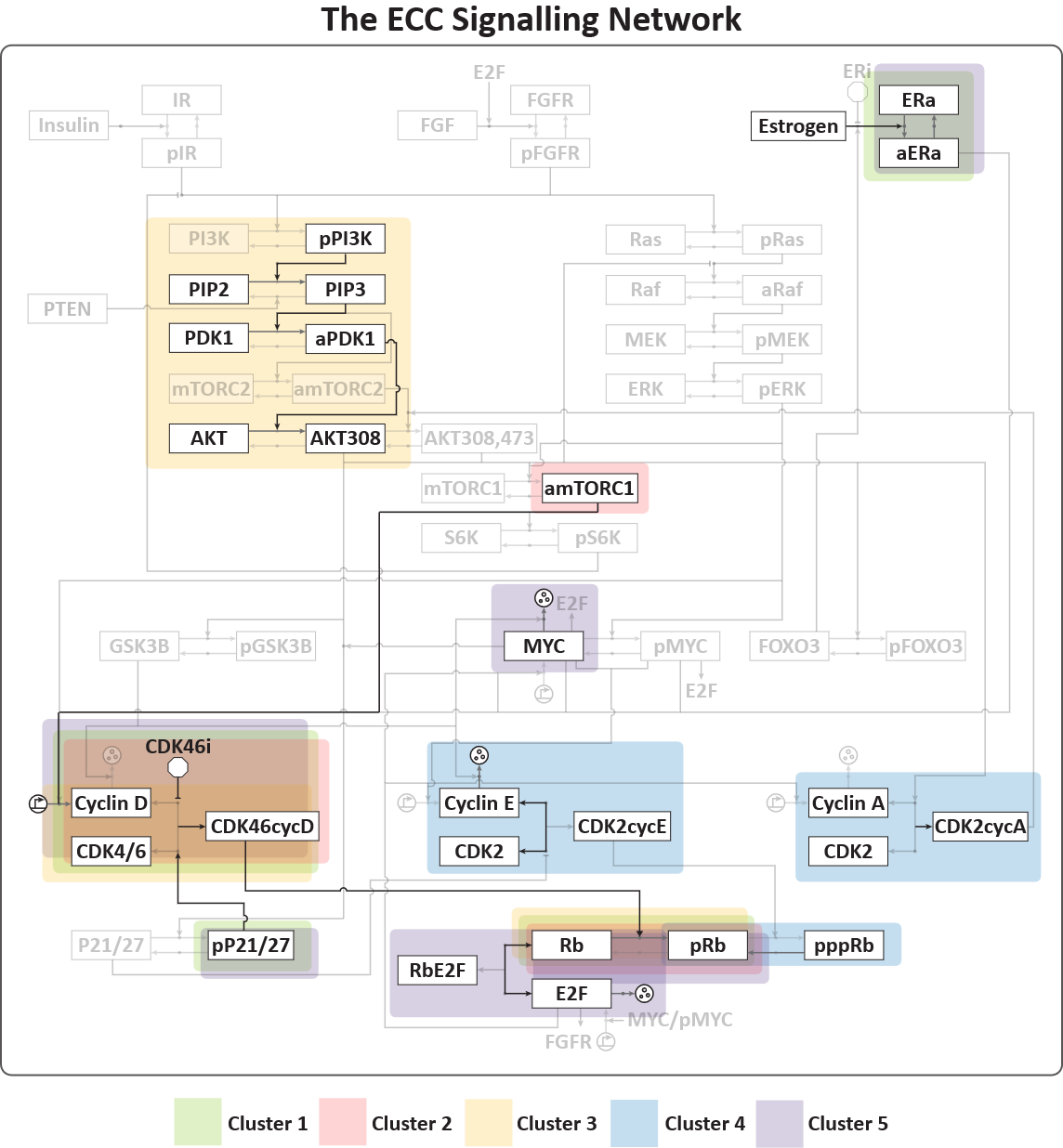

**Figure S10C. Cluster-specific parameter signatures with the ECC network, here displayed for the protein-dynamic pRb-REB combination.** Each figure (Figures S9A-I) is the overlay of the respective cluster-specific parameter signatures with the ECC network, for each protein-dynamic combination. Each figure displays the five largest clusters for each protein-dynamic combination.

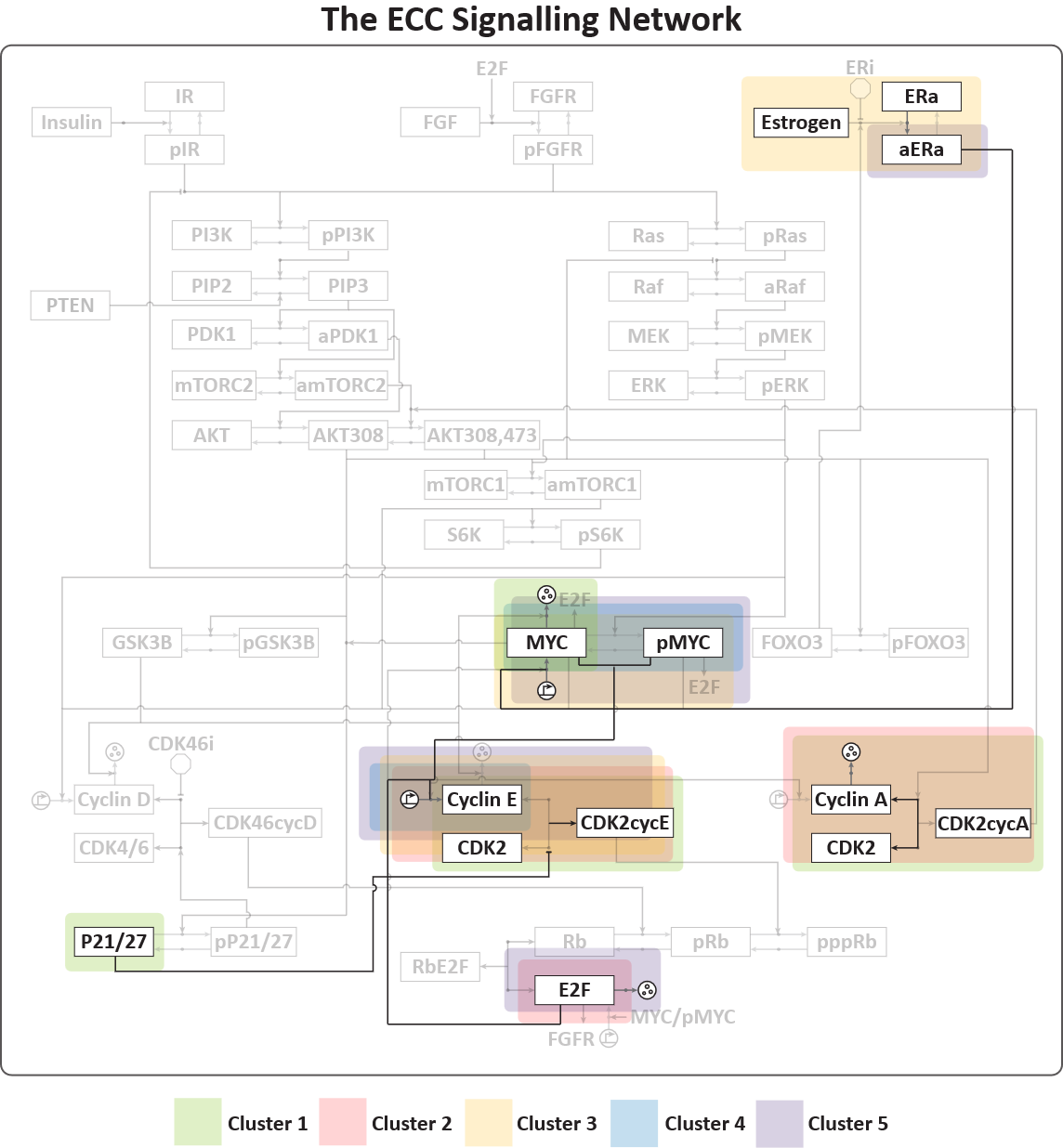

**Figure S10D. Cluster-specific parameter signatures with the ECC network, here displayed for the protein-dynamic CDK2cycE-INC combination.** Each figure (Figures S9A-I) is the overlay of the respective cluster-specific parameter signatures with the ECC network, for each protein-dynamic combination. Each figure displays the five largest clusters for each protein-dynamic combination.

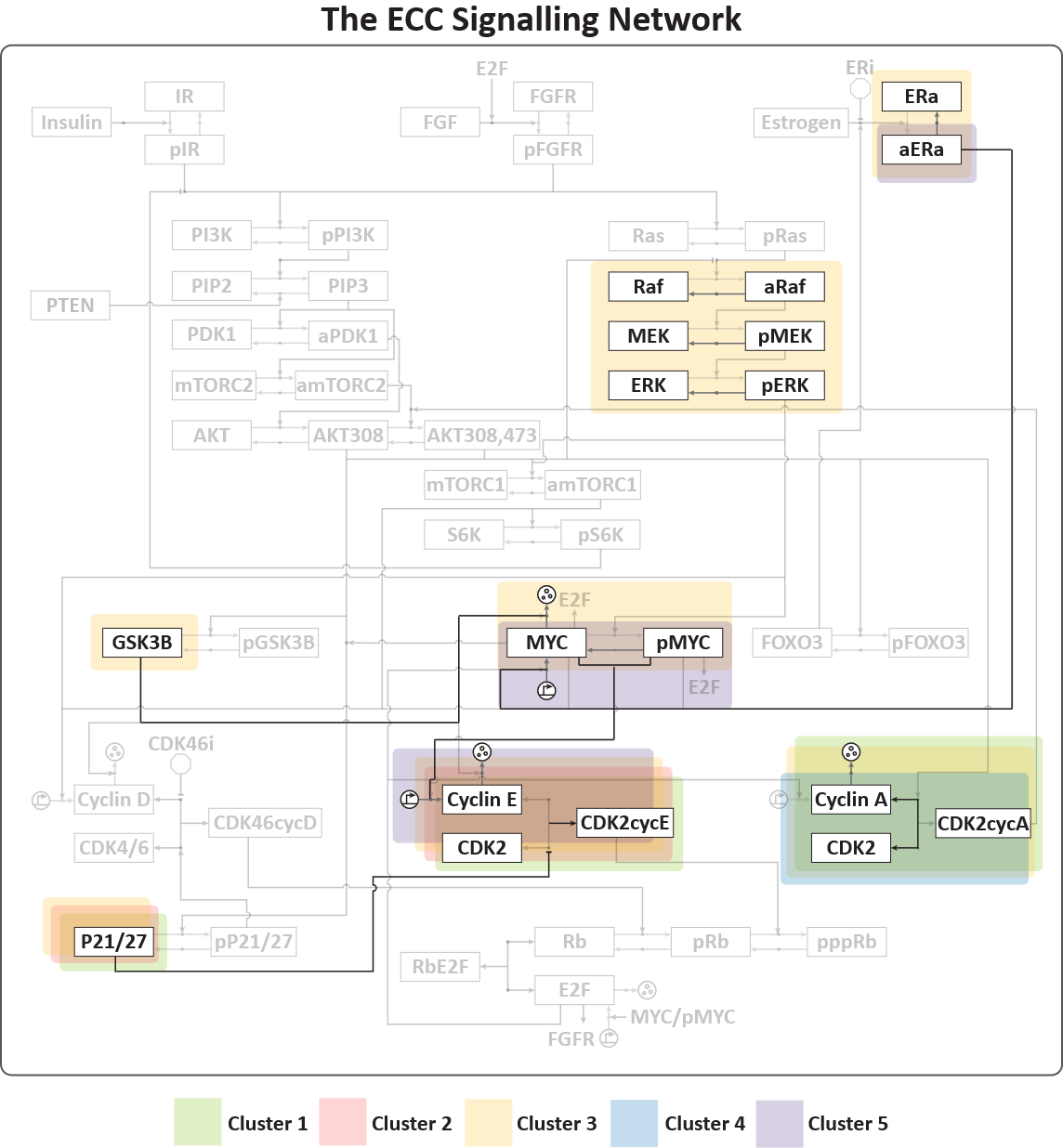

**Figure S10E. Cluster-specific parameter signatures with the ECC network, here displayed for the protein-dynamic CDK2cycE-BIP combination.** Each figure (Figures S9A-I) is the overlay of the respective cluster-specific parameter signatures with the ECC network, for each protein-dynamic combination. Each figure displays the five largest clusters for each protein-dynamic combination.

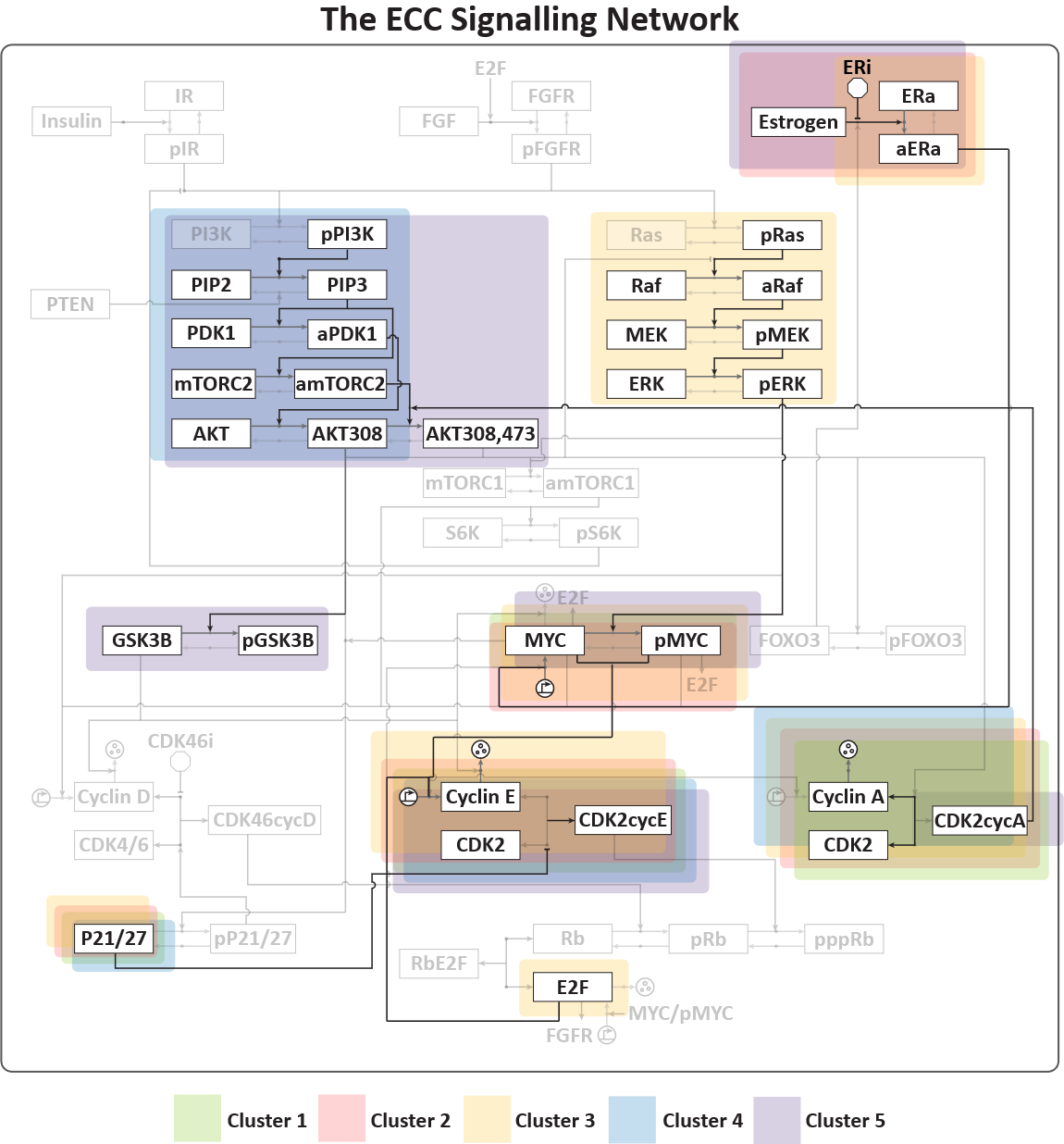

**Figure S10F. Cluster-specific parameter signatures with the ECC network, here displayed for the protein-dynamic CDK2cycE-REB combination.** Each figure (Figures S9A-I) is the overlay of the respective cluster-specific parameter signatures with the ECC network, for each protein-dynamic combination. Each figure displays the five largest clusters for each protein-dynamic combination.

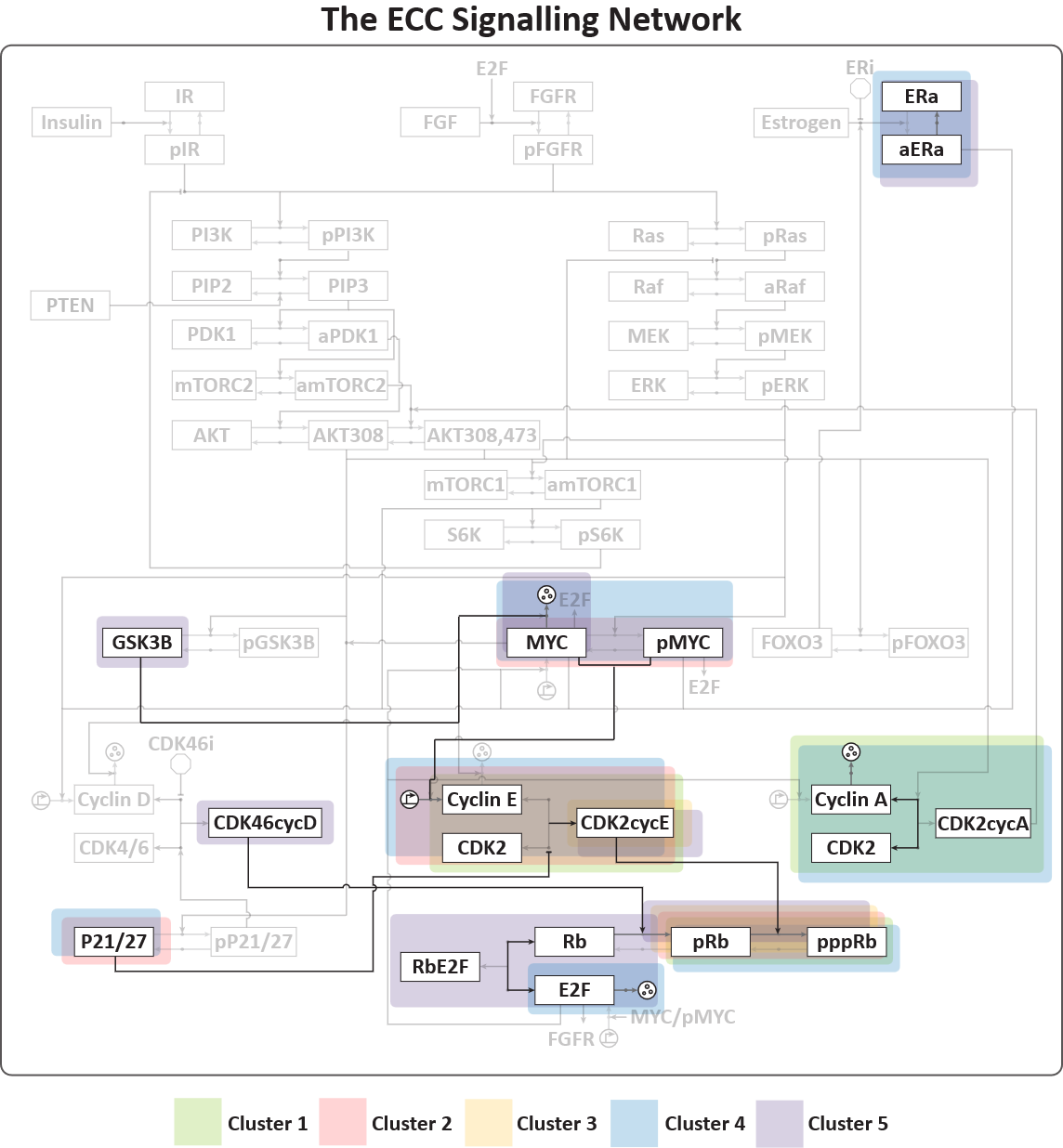

**Figure S10G. Cluster-specific parameter signatures with the ECC network, here displayed for the protein-dynamic pppRb-INC combination.** Each figure (Figures S9A-I) is the overlay of the respective cluster-specific parameter signatures with the ECC network, for each protein-dynamic combination. Each figure displays the five largest clusters for each protein-dynamic combination.

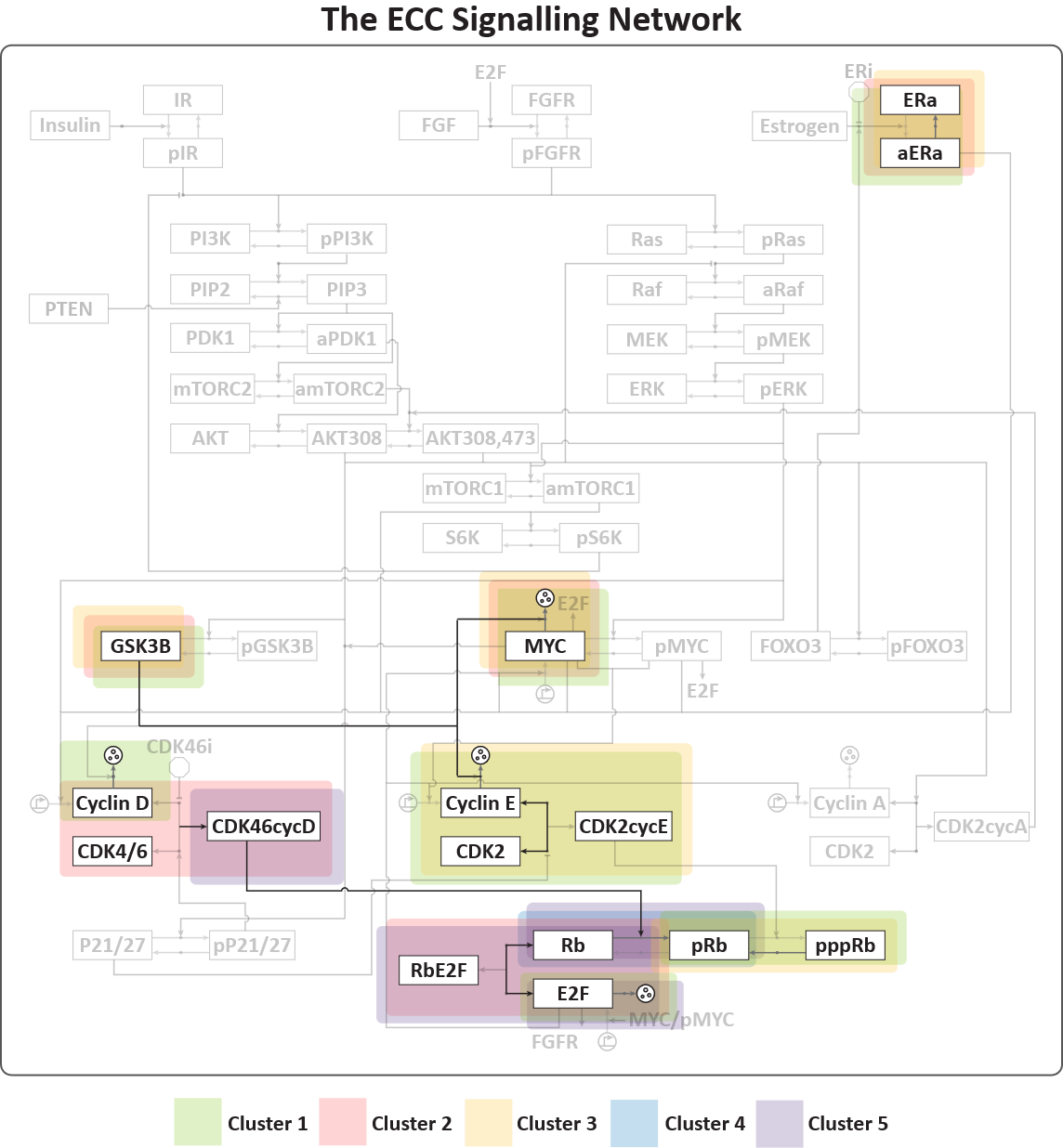

**Figure S10H. Cluster-specific parameter signatures with the ECC network, here displayed for the protein-dynamic pppRb-BIP combination.** Each figure (Figures S9A-I) is the overlay of the respective cluster-specific parameter signatures with the ECC network, for each protein-dynamic combination. Each figure displays the five largest clusters for each protein-dynamic combination.

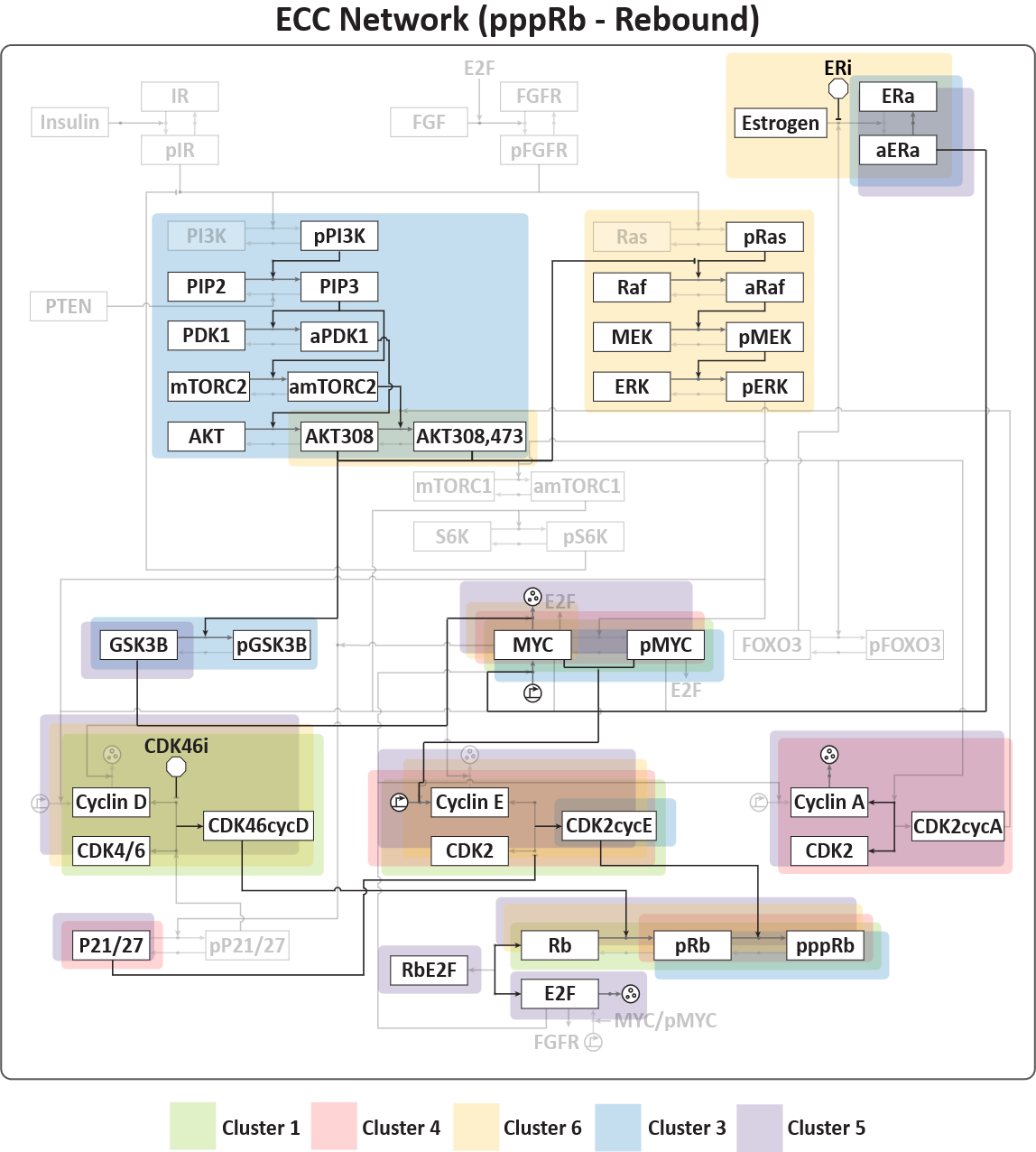

**Figure S10I. Cluster-specific parameter signatures with the ECC network, here displayed for the protein-dynamic pppRb-REB combination.** Each figure (Figures S9A-I) is the overlay of the respective cluster-specific parameter signatures with the ECC network, for each protein-dynamic combination. Each figure displays the five largest clusters for each protein-dynamic combination.

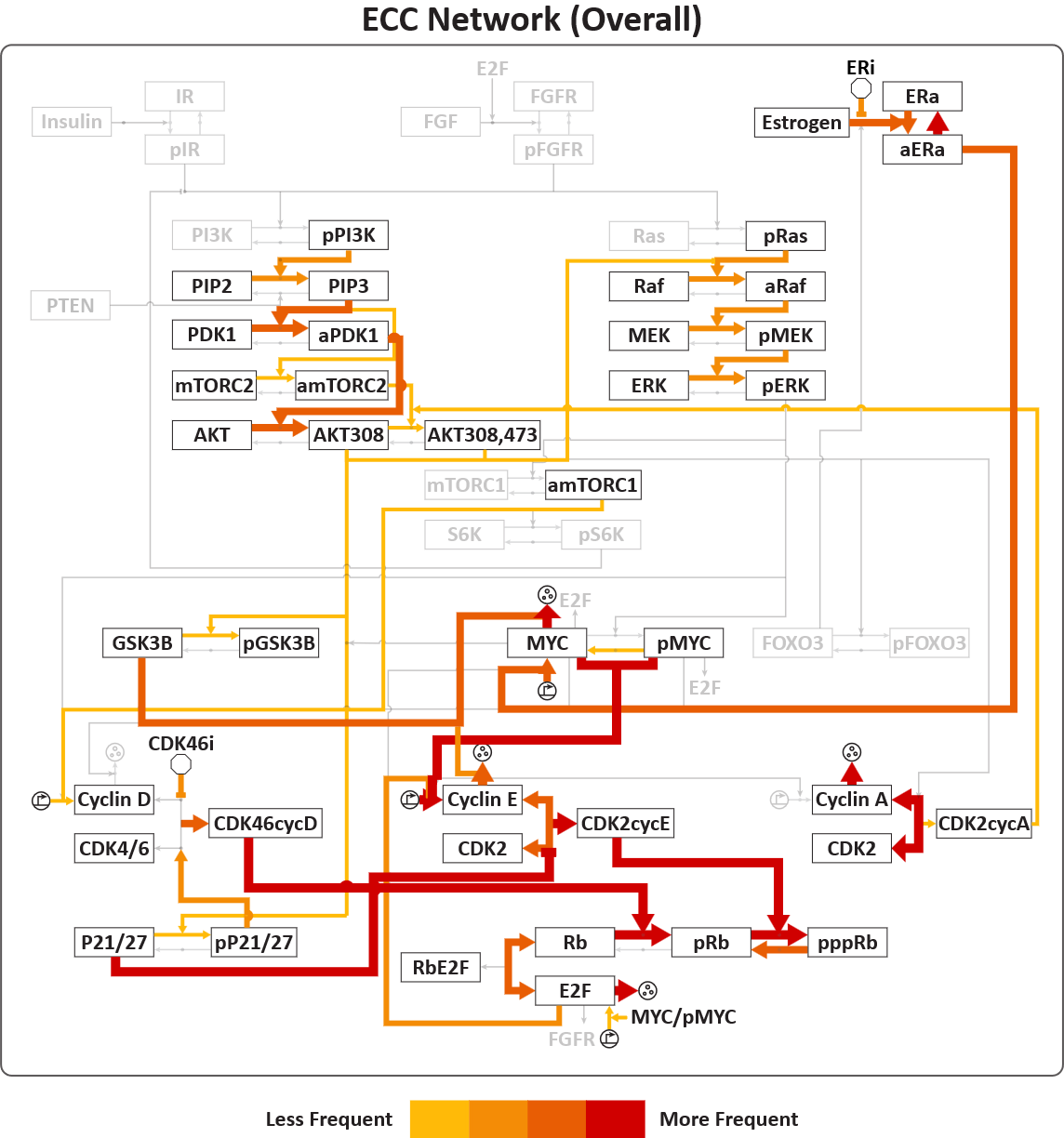

**Figure S11. Summary of all the parameter-signature clusters (subnetworks) that drive resistance identified for 9 protein-dynamic combinations.** This was derived from the individual analysis results in Figures S8 and Figures S9A-I.

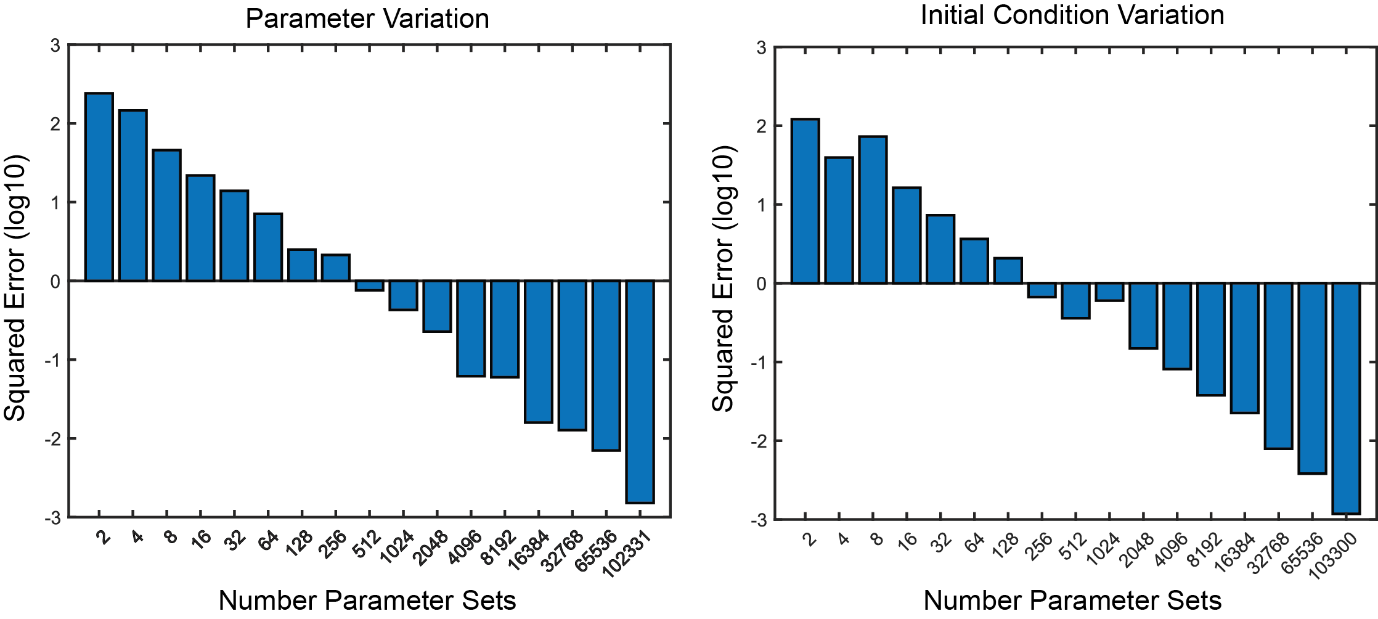

**Figure S12.** Bar graph showing the total change in the frequency of dynamics when the number of model instances is doubled. Specifically, we calculated the Mean Squared Error between the distributions of protein dynamics for model instance populations of size N and 2N. Y-axis is the calculated MSE and is in log10 scale, and the X axis shows the size of population N.

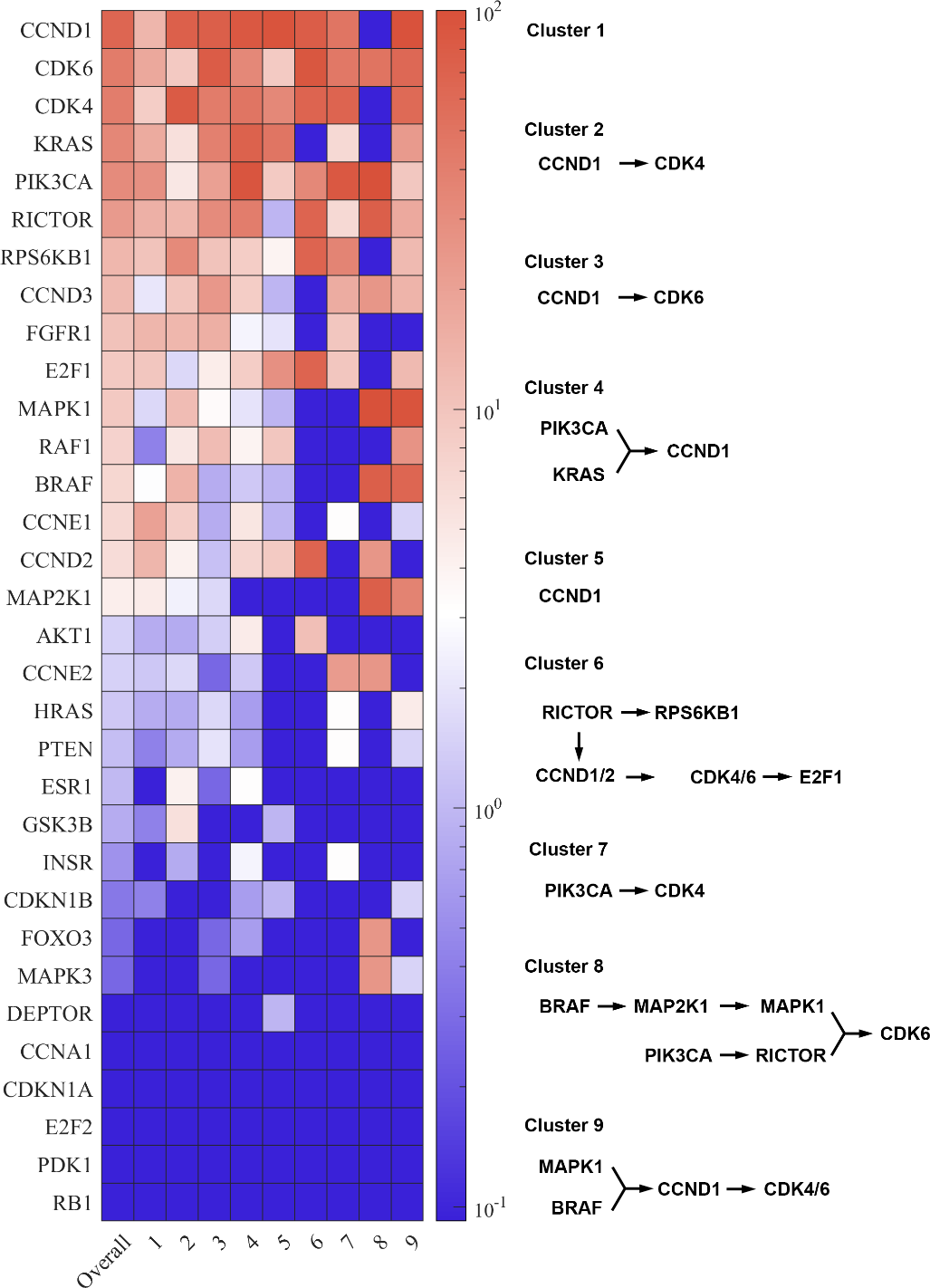

**Figure S13.** Heatmap showing the clustered differential gene-dependencies of hundreds of cancer cell lines. Genes were filtered for moderate dependency and then subject to hierarchical clustering to identify common signatures of gene dependency across approximately 1000 cancer cell lines. The gene signatures identified in this analysis are quasi-similar to the idea of drug resistance signatures identified using our MDN analysis.

### **Supplementary Tables**

**Table S1. Nominal Parameter Units and Values.**

This table represents typical parameter values for use in ODE modelling for each of the different parameter types. It is important to note that these parameters vary quite dramatically from protein interaction to protein interaction and so are an extremely rough estimate. However, they are useful as a base starting point from which to test ODE models.

**Table S2A – J. Parametric resistance signatures for all 9 (resistance) protein-dynamic combinations. Also includes overall parameter contributions to resistance dynamics.**

The contribution of each parameter to the resistance features of each resistance associated protein-dynamic combination was measured and subjected to a modified hierarchical clustering. In this clustering process, clusters were recursively clustered until clusters contained at least one parameter that represented at least 80% of the model instances comprising the cluster. Additionally, the clusters had to contain at least 1% of the total number of model instances to ensure the clusters represented a robust parametric resistance signature. Parameters were considered to contribute to the resistance signature if they occurred in more than 50% of the model instances within the cluster. The final result is a list of resistance driving parametric signatures for each resistance associated protein-dynamic combination.

Each table shows the identified clusters (resistance signatures) for each resistance associated protein-dynamic combination, the frequency at which they occur, as a percentage of the total number of model instances within each protein-dynamic, and the parameters that comprise each resistance signature.

**Table S3. Table containing the known resistance mechanisms and the respective resistance-driving parameters identified by our MDN analysis.**

**Supplementary Information file.** Detailed description of Model Scope and Construction

### **Supplementary Files**

**Model File S1. Model – Biochemical Reaction format.**

**Model File S2. Model – ODE format.**

**Model File S3. Model - SBML format.**

### **Supplementary References.**

1.	D. A. E. Cross, D. R. Alessi, P. Cohen, M. Andjelkovich, B. A. Hemmings, Inhibition of glycogen synthase kinase-3 by insulin mediated by protein kinase B. *Nature* **378**, 785-789 (1995)10.1038/378785a0).

2.	J. Averous, B. D. Fonseca, C. G. Proud, Regulation of cyclin D1 expression by mTORC1 signaling requires eukaryotic initiation factor 4E-binding protein 1. *Oncogene* **27**, 1106-1113 (2008); published online Epub2008/02/01 (10.1038/sj.onc.1210715).

3.	M. Welcker, J. Singer, K. R. Loeb, J. Grim, A. Bloecher, M. Gurien-West, B. E. Clurman, J. M. Roberts, Multisite Phosphorylation by Cdk2 and GSK3 Controls Cyclin E Degradation. *Molecular Cell* **12**, 381-392 (2003); published online Epub2003/08/01/ (<https://doi.org/10.1016/S1097-2765(03)00287-9>).

4.	R. C. Sears, The life cycle of C-myc: from synthesis to degradation. *Cell Cycle* **3**, 1133-1137 (2004); published online EpubSep (

5.	J. Zhang, Z. Gao, J. Yin, M. J. Quon, J. Ye, S6K directly phosphorylates IRS-1 on Ser-270 to promote insulin resistance in response to TNF-(alpha) signaling through IKK2. *J Biol Chem* **283**, 35375-35382 (2008); published online EpubDec 19 (10.1074/jbc.M806480200).

6.	K. Balmanno, S. J. Cook, Sustained MAP kinase activation is required for the expression of cyclin D1, p21Cip1 and a subset of AP-1 proteins in CCL39 cells. *Oncogene* **18**, 3085-3097 (1999); published online EpubMay 20 (10.1038/sj.onc.1202647).

7.	T. K. Hayes, N. F. Neel, C. Hu, P. Gautam, M. Chenard, B. Long, M. Aziz, M. Kassner, K. L. Bryant, M. Pierobon, R. Marayati, S. Kher, S. D. George, M. Xu, A. Wang-Gillam, A. A. Samatar, A. Maitra, K. Wennerberg, E. F. Petricoin, H. H. Yin, B. Nelkin, A. D. Cox, J. J. Yeh, C. J. Der, Long-Term ERK Inhibition in KRAS-Mutant Pancreatic Cancer Is Associated with MYC Degradation and Senescence-like Growth Suppression. *Cancer Cell* **29**, 75-89 (2016); published online Epub2016/01/11/ (<https://doi.org/10.1016/j.ccell.2015.11.011>).

8.	K. D. Hanson, M. Shichiri, M. R. Follansbee, J. M. Sedivy, Effects of c-myc expression on cell cycle progression. *Mol Cell Biol* **14**, 5748-5755 (1994); published online EpubSep (10.1128/mcb.14.9.5748-5755.1994).

9.	G. Leone, J. DeGregori, R. Sears, L. Jakoi, J. R. Nevins, Myc and Ras collaborate in inducing accumulation of active cyclin E/Cdk2 and E2F. *Nature* **387**, 422-426 (1997); published online Epub1997/05/01 (10.1038/387422a0).

10.	R. Beier, A. Bürgin, A. Kiermaier, M. Fero, H. Karsunky, R. Saffrich, T. Möröy, W. Ansorge, J. Roberts, M. Eilers, Induction of cyclin E–cdk2 kinase activity, E2F-dependent transcription and cell growth by Myc are genetically separable events. *The EMBO journal* **19**, 5813-5823 (2000).

11.	G. Leone, R. Sears, E. Huang, R. Rempel, F. Nuckolls, C. H. Park, P. Giangrande, L. Wu, H. I. Saavedra, S. J. Field, M. A. Thompson, H. Yang, Y. Fujiwara, M. E. Greenberg, S. Orkin, C. Smith, J. R. Nevins, Myc requires distinct E2F activities to induce S phase and apoptosis. *Molecular Cell* **8**, 105-113 (2001)10.1016/S1097-2765(01)00275-1).

12.	G. Bretones, M. D. Delgado, J. León, Myc and cell cycle control. *Biochimica et Biophysica Acta (BBA) - Gene Regulatory Mechanisms* **1849**, 506-516 (2015); published online Epub2015/05/01/ (<https://doi.org/10.1016/j.bbagrm.2014.03.013>).

13.	Y. Li, D. Dowbenko, L. A. Lasky, AKT/PKB phosphorylation of p21Cip/WAF1 enhances protein stability of p21Cip/WAF1 and promotes cell survival. *Journal of Biological Chemistry* **277**, 11352-11361 (2002).

14.	M. R. Bratton, J. W. Antoon, B. N. Duong, D. E. Frigo, S. Tilghman, B. M. Collins-Burow, S. Elliott, Y. Tang, L. I. Melnik, L. Lai, J. Alam, B. S. Beckman, S. M. Hill, B. G. Rowan, J. A. McLachlan, M. E. Burow, Gαo potentiates estrogen receptor α activity via the ERK signaling pathway. *J Endocrinol* **214**, 45-54 (2012)10.1530/JOE-12-0097).

15.	M. Marino, F. Acconcia, F. Bresciani, A. Weisz, A. Trentalance, Distinct nongenomic signal transduction pathways controlled by 17β-estradiol regulate DNA synthesis and cyclin D1 gene transcription in HepG2 cells. *Molecular biology of the cell* **13**, 3720-3729 (2002).

16.	C. Dong, S. B. Waters, K. H. Holt, J. E. Pessin, SOS phosphorylation and disassociation of the Grb2-SOS complex by the ERK and JNK signaling pathways. *J Biol Chem* **271**, 6328-6332 (1996); published online EpubMar 15 (10.1074/jbc.271.11.6328).

17.	S. Zimmermann, K. Moelling, Phosphorylation and regulation of Raf by Akt (protein kinase B). *Science* **286**, 1741-1744 (1999).

18.	Y. Geng, E. N. Eaton, M. Picon, J. M. Roberts, A. S. Lundberg, A. Gifford, C. Sardet, R. A. Weinberg, Regulation of cyclin E transcription by E2Fs and retinoblastoma protein. *Oncogene* **12**, 1173-1180 (1996).

19.	B. Henglein, X. Chenivesse, J. Wang, D. Eick, C. Bréchot, Structure and cell cycle-regulated transcription of the human cyclin A gene. *Proceedings of the National Academy of Sciences of the United States of America* **91**, 5490-5494 (1994)10.1073/pnas.91.12.5490).

20.	A. M. Narasimha, M. Kaulich, G. S. Shapiro, Y. J. Choi, P. Sicinski, S. F. Dowdy, Cyclin D activates the Rb tumor suppressor by mono-phosphorylation. *eLife* **3**, e02872 (2014); published online Epub2014/05/29 (10.7554/eLife.02872).

21.	P. Liu, Z. Wang, W. Wei, Phosphorylation of Akt at the C-terminal tail triggers Akt activation. *Cell Cycle* **13**, 2162-2164 (2014).

22.	J. Zhang, K. Xu, P. Liu, Y. Geng, B. Wang, W. Gan, J. Guo, F. Wu, Y. R. Chin, C. Berrios, Evan C. Lien, A. Toker, James A. DeCaprio, P. Sicinski, W. Wei, Inhibition of Rb Phosphorylation Leads to mTORC2-Mediated Activation of Akt. *Molecular Cell* **62**, 929-942 (2016); published online Epub2016/06/16/ (<https://doi.org/10.1016/j.molcel.2016.04.023>).

23.	J. LaBaer, M. D. Garrett, L. F. Stevenson, J. M. Slingerland, C. Sandhu, H. S. Chou, A. Fattaey, E. Harlow, New functional activities for the p21 family of CDK inhibitors. *Genes & development* **11**, 847-862 (1997).

24.	G. C. Bao, J.-G. Wang, A. Jong, Increased p21 expression and complex formation with cyclin E/CDK2 in retinoid-induced pre-B lymphoma cell apoptosis. *FEBS Letters* **580**, 3687-3693 (2006); published online Epub2006/06/26/ (<https://doi.org/10.1016/j.febslet.2006.05.052>).

25.	M. D. Planas-Silva, Y. Shang, J. L. Donaher, M. Brown, R. A. Weinberg, AIB1 enhances estrogen-dependent induction of cyclin D1 expression. *Cancer research* **61**, 3858-3862 (2001).

26.	C. Jiang, M. Ito, V. Piening, K. Bruck, R. G. Roeder, H. Xiao, TIP30 interacts with an estrogen receptor α-interacting coactivator CIA and regulates c-myc transcription. *Journal of Biological Chemistry* **279**, 27781-27789 (2004).

27.	K. Belguise, S. Guo, G. E. Sonenshein, Activation of FOXO3a by the green tea polyphenol epigallocatechin-3-gallate induces estrogen receptor α expression reversing invasive phenotype of breast cancer cells. *Cancer research* **67**, 5763-5770 (2007).

28.	M. Dobson, G. Ramakrishnan, S. Ma, L. Kaplun, V. Balan, R. Fridman, G. Tzivion, Bimodal regulation of FoxO3 by AKT and 14-3-3. *Biochimica et Biophysica Acta (BBA)-Molecular Cell Research* **1813**, 1453-1464 (2011).
