## Supplementary material for "Systematic Analysis of Network-driven Adaptive Resistance to CDK4/6 and Estrogen Receptor Inhibition using Meta-Dynamic Network Modelling": Model File 4

### Early Cell Cycle Model - ODEs

Anthony Hart

December 2025

#### 1 State Variables

$$\frac{d[\text{IR}]}{dt} = -R1$$
$$\frac{d[\text{pIR}]}{dt} = +R1$$

$$\frac{d[\text{PI3K}]}{dt} = -R2$$
$$\frac{d[\text{pPI3K}]}{dt} = +R2$$

$$\frac{d[\text{PIP2}]}{dt} = -R3$$
$$\frac{d[\text{PIP3}]}{dt} = +R3$$

$$\frac{d[\text{PDK1}]}{dt} = -R4$$
$$\frac{d[\text{aPDK1}]}{dt} = +R4$$

$$\frac{d[\text{mTORC2}]}{dt} = -R5$$
$$\frac{d[\text{amTORC2}]}{dt} = +R5$$

$$\frac{d[\text{AKT}]}{dt} = -R6$$
$$\frac{d[\text{pAKT308}]}{dt} = +R6 - R7$$
$$\frac{d[\text{ppAKT308473}]}{dt} = +R7$$

$$\frac{d[\text{mTORC1}]}{dt} = -R8$$
$$\frac{d[\text{amTORC1}]}{dt} = +R8$$

$$\frac{d[\text{S6K}]}{dt} = -R9$$
$$\frac{d[\text{pS6K}]}{dt} = +R9$$

$$\frac{d[\text{FGFR}]}{dt} = -R10$$
$$\frac{d[\text{pFGFR}]}{dt} = +R10$$

$$\frac{d[\mathrm{Ras}]}{dt} = -R11$$
$$\frac{d[\mathrm{aRas}]}{dt} = +R11$$

$$\frac{d[\mathrm{Raf}]}{dt} = -R12$$
$$\frac{d[\mathrm{aRaf}]}{dt} = +R12$$

$$\frac{d[\mathrm{MEK}]}{dt} = -R13$$
$$\frac{d[\mathrm{pMEK}]}{dt} = +R13$$

$$\frac{d[\mathrm{ERK}]}{dt} = -R14$$
$$\frac{d[\mathrm{pERK}]}{dt} = +R14$$

$$\frac{d[\mathrm{ERa}]}{dt} = -R15$$
$$\frac{d[\mathrm{aERa}]}{dt} = +R15$$

$$\frac{d[\mathrm{Myc}]}{dt} = +R16 - R17 - R18$$
$$\frac{d[\mathrm{pMyc}]}{dt} = +R18$$

$$\frac{d[\mathrm{GSK3b}]}{dt} = -R19$$
$$\frac{d[\mathrm{pGSK3b}]}{dt} = +R19$$

$$\frac{d[\mathrm{FOXO3}]}{dt} = -R20$$
$$\frac{d[\mathrm{pFOXO3}]}{dt} = +R20$$

$$\frac{d[\mathrm{cycD}]}{dt} = +R21 - R22 - R27$$
$$\frac{d[\mathrm{cycE}]}{dt} = +R23 - R24 - R28$$
$$\frac{d[\mathrm{cycA}]}{dt} = +R25 - R26 - R29$$

$$\frac{d[\mathrm{CDK46}]}{dt} = -R27$$
$$\frac{d[\mathrm{CDK2}]}{dt} = -R28 - R29$$

$$\frac{d[\mathrm{CDK46cycD}]}{dt} = +R27$$
$$\frac{d[\mathrm{CDK2cycE}]}{dt} = +R28$$
$$\frac{d[\mathrm{CDK2cycA}]}{dt} = +R29$$

$$\frac{d[p21\_27]}{dt} = -R30$$

$$\frac{d[pp21\_27]}{dt} = +R30$$

$$\frac{d[\mathrm{E2F}]}{dt} = +R31 - R32 - R33$$

$$\frac{d[\mathrm{RbE2F}]}{dt} = +R33$$

$$\frac{d[\mathrm{Rb}]}{dt} = -R33 - R34$$

$$\frac{d[\mathrm{pRb}]}{dt} = +R34 - R35$$

$$\frac{d[\mathrm{pppRb}]}{dt} = +R35$$

#### 2 ODEs

$$R1 = (k_{c1f1}\,\text{Ins}\,\text{IR}) - (V_{\max,1r1}\,p\text{IR}) \qquad \{\text{reversible}\}$$

$$R2 = \left(\frac{k_{c2f1}\,p\text{IR}\,\text{PI3K}}{1+(p\text{S6K}/K_{i2})} + k_{c2f2}\,p\text{FGFR}\,\text{PI3K}\right) - (V_{\max,2r1}\,p\text{PI3K}) \qquad \{\text{reversible}\}$$

$$R3 = (k_{c3f1}\,p\text{PI3K}\,\text{PIP2}) - ((V_{\max,3r1} + k_{c3r1}\,\text{PTEN})\,\text{PIP3}) \qquad \{\text{reversible}\}$$

$$R4 = (k_{c4f1}\,\text{PIP3}\,\text{PDK1}) - (V_{\max,4r1}\,a\text{PDK1}) \qquad \{\text{reversible}\}$$

$$R5 = \left(\frac{k_{c5f1}\,\text{PIP3}\,\text{mTORC2}}{1+(ppp\text{Rb}/K_{i5})}\right) - (V_{\max,5r1}\,a\text{mTORC2}) \qquad \{\text{reversible}\}$$

$$R6 = (k_{c6f1}\,a\text{PDK1}\,\text{AKT}) - (V_{\max,6r1}\,p\text{AKT308}) \qquad \{\text{reversible}\}$$

$$R7 = \left(k_{c7f1}\,a\text{mTORC2}\,p\text{AKT308}\left(1+\alpha_7\,\text{CDK2cycA}\right)\right) - (V_{\max,7r1}\,pp\text{AKT308473}) \qquad \{\text{reversible}\}$$

$$R8 = ((k_{c8f1}\,p\text{AKT308} + \text{AKTscale}\,k_{c8f1}\,pp\text{AKT308473})\,\text{mTORC1}) - (V_{\max,8r1}\,a\text{mTORC1}) \qquad \{\text{reversible}\}$$

$$R9 = (k_{c9f1}\,a\text{mTORC1}\,\text{S6K}) - (V_{\max,9r1}\,p\text{S6K}) \qquad \{\text{reversible}\}$$

$$R10 = (k_{c10f1}\,\text{FGF}\,\text{FGFR})(1+\alpha_{10}\text{E2F}) - (V_{\max,10r1}\,p\text{FGFR}) \qquad \{\text{reversible}\}$$

$$R11 = \left(\frac{k_{c11f1}\,p\text{FGFR} + k_{c11f2}\,p\text{IR}}{1+(p\text{ERK}/K_{i11})}\,\text{Ras}\right) - (V_{\max,11r1}\,a\text{Ras}) \qquad \{\text{reversible}\}$$

$$R12 = \left(\frac{k_{c12f1}\,a\text{Ras}\,\text{Raf}}{1+(p\text{AKT308}/K_{i12}) + (pp\text{AKT308473}/(\text{AKTscale}\,K_{i12}))}\right) - (V_{\max,12r1}\,a\text{Raf}) \qquad \{\text{reversible}\}$$

$$R13 = (k_{c13f1}\,a\text{Raf}\,\text{MEK}) - (V_{\max,13r1}\,p\text{MEK}) \qquad \{\text{reversible}\}$$

$$R14 = (k_{c14f1}\,p\text{MEK}\,\text{ERK}) - (V_{\max,14r1}\,p\text{ERK}) \qquad \{\text{reversible}\}$$

$$R15 = \left(\frac{k_{c15f1}\,\text{Est}\,\text{ERa}(1+\alpha_{15}\text{FOXO3})}{1+(E_{i0}/K_{i15})}\right) - (V_{\max,15r1}\,a\text{ERa}) \qquad \{\text{reversible}\}$$

$$R16 = k_{c16f1}\,a\text{ERa}(1+\alpha_{16a}p\text{ERK} + \alpha_{16b}\text{cycD}) + k_{c16f2}\text{E2F}$$

$$R17 = k_{\text{deg}17f1}\,\text{Myc}(1+\alpha_{17}\text{GSK3b})$$

$$R18 = (k_{c18f1}\,p\text{ERK}\,\text{Myc}) - (V_{\max,18r1}\,p\text{Myc}) \qquad \{\text{reversible}\}$$

$$R19 = ((k_{c19f1}\,p\text{AKT308} + \text{AKTscale}\,k_{c19f1}\,pp\text{AKT308473})\text{GSK3b}) - (V_{\max,19r1}\,p\text{GSK3b}) \qquad \{\text{reversible}\}$$

$$R20 = ((k_{c20f1}\,p\text{AKT308} + \text{AKTscale}\,k_{c20f1}\,pp\text{AKT308473})\text{FOXO3}) - (V_{\max,20r1}\,p\text{FOXO3}) \qquad \{\text{reversible}\}$$

$$R21 = k_{c21f1}a\text{mTORC1} + k_{c21f2}p\text{ERK} + k_{c21f3}(\text{Myc} + p\text{Myc}) + k_{c21f4}a\text{ERa}(1+\alpha_{21a}p\text{ERK} + \alpha_{21b}\text{cycD})$$

$$R22 = k_{\text{deg}22f1}\,\text{cycD}(1+\alpha_{22}\text{GSK3b})$$

$$R23 = k_{c23f1}\text{E2F}(1+\alpha_{23}(\text{Myc} + p\text{Myc})) + k_{c23f2}(\text{Myc} + p\text{Myc})$$

$$R24 = k_{\text{deg}24f1}\,\text{cycE}(1+\alpha_{24}\text{GSK3b})$$

$$R25 = k_{c25f1}\text{E2F}(1+\alpha_{25}(\text{Myc} + p\text{Myc})) + k_{c25f2}(\text{Myc} + p\text{Myc})$$

$$R26 = k_{\text{deg}26f1}\,\text{cycA}$$

$$R27 = \left(\frac{k_{a27f1}\,\text{CDK46}\,\text{cycD}(1+\alpha_{27}pp\text{21\_27})}{1+(\text{CDK46i}_0/K_{i27})}\right) - (k_{d27r1}\,\text{CDK46cycD}) \qquad \{\text{reversible}\}$$

$$R28 = \left( \frac{k_{a28f1}\,\mathrm{CDK2\,cycE}}{1 + (p21\_27/K_{i28})} \right) - (k_{d28r1}\,\mathrm{CDK2cycE}) \qquad \{\mathrm{reversible}\}$$

$$R29 = k_{a29f1}\,\mathrm{CDK2\,cycA}\,(1 + \alpha_{29} p\mathrm{AKT308} + \mathrm{AKTscale}\,\alpha_{29} pp\mathrm{AKT308473}) - (k_{d29r1}\,\mathrm{CDK2cycA}) \qquad \{\mathrm{reversible}\}$$

$$R30 = ((k_{c30f1} p\mathrm{AKT308} + \mathrm{AKTscale}\,k_{c30f1}\,pp\mathrm{AKT308473})\,p21\_27(1 + \alpha_{30}(\mathrm{Myc} + p\mathrm{Myc}))) \\ - (V_{\mathrm{max},30r1}\,pp21\_27) \qquad \{\mathrm{reversible}\}$$

$$R31 = k_{c31f1}(\mathrm{Myc} + p\mathrm{Myc}) + k_{c31f2}\mathrm{E2F}$$

$$R32 = k_{\mathrm{deg}32f1}\,\mathrm{E2F}$$

$$R33 = k_{a33f1}\mathrm{E2FRb} - (k_{d33r1}\mathrm{RbE2F}) \qquad \{\mathrm{reversible}\}$$

$$R34 = k_{c34f1}\mathrm{CDK46cycD\,Rb} - (V_{\mathrm{max},34r1}\,p\mathrm{Rb}) \qquad \{\mathrm{reversible}\}$$

$$R35 = k_{c35f1}\mathrm{CDK2cycE}\,p\mathrm{Rb} - (V_{\mathrm{max},35r1}\,ppp\mathrm{Rb}) \qquad \{\mathrm{reversible}\}$$
